# Ferroptosis assassinates tumor (FAST)

**DOI:** 10.1101/2021.10.04.463002

**Authors:** Tao Luo, Yile Wang, Jinke Wang

## Abstract

In 2020, nearly 20 million peoples got cancer and nearly 10 million peoples died of cancer, indicating the current therapies do not meet the cancer treatment and cancer remains a great threat to human health and life. New therapies are still in urgent demand. In a recent study, we developed a new effective cancer therapy, gene-interfered ferroptosis therapy (GIFT), by combining cancer cell-specific knockdown of two iron efflux genes (FPN and LCN2) with iron nanoparticles (FeNPs). GIFT shows wide antitumor activity, high cancer specificity, certain cancer eradication potential, and biosafety. To further improve the therapy, we here develop an updated GIFT named as Ferroptosis ASsassinates Tumor (FAST) by knocking down five additional ferroptosis-resistance genes (FSP1, FTH1, GPX4, SLC7A11, NRF2). As a result, we found that FAST showed more significant antitumor activity than GIFT. Especially, FAST eradicated three different types of tumors (leukemia, colon cancer and lung metastatic melanoma) from over 50 percent of cancer mice, making the mice to survive up to 250 days without tumor relapse. FAST also significantly inhibited and prevented growth of spontaneous breast cancer and improved survival in mice. Additionally, FAST showed high pan-antitumor efficacy, high cancer specificity, and in vivo safety.

---

According to the International Agency for Research on Cancer (IARC) of World Health Organization (WHO), in 2020, as many as 19.29 million people were newly diagnosed as cancer and about 9.96 million people died of cancer in the world wide, and as many as 4.57 million people were newly diagnosed as cancer and about 3 million people died of cancer in China. Cancer is therefore a major stress for public well-being and the most dreadful disease. Cancer is still a major burden of disease and a leading cause of death worldwide. Cancer therapies such as surgery, chemotherapy, radiotherapy, targeted therapy, and immunotherapy have evolved considerably in recent decades and substantially improve the quality of life and survival of patients with cancer. However, the current cancer therapies are still challenged by several key limitations such as low response, drug resistance, side effects, and recurrence. Therefore, there is still a huge gap between the efficacy of cancer treatment and the desire of patients for health and life. Cancer treatment urgently needs breakthrough technology.

Ferroptosis is a cell death form induced by iron-dependent lipid peroxidation (*1, 2*), in which ferrous iron (Fe^2+^) produces hydroxyl radicals (•OH) via Fenton reaction (*3, 4*). Since coined, the antitumor effects of ferroptosis has been widely investigated in variant cancers (*5*). Various compounds for inhibiting or depleting system x_c_^−^, GPX4 and CoQ10 are identified as ferroptosis inducers (*6*); however, these inducers are still challenged by side effects and drug resistance (*7–9*). Iron oxide nanoparticles (IONPs) can generate iron ions (Fe^2+^/Fe^3+^) in lysosomes when internalized in cells and the generated Fe^2+^ can produce •OH by Fenton reaction, which potentially induce ferroptosis (*10–12*). However, cells can overcome this potential damage by positively maintaining the iron and redox homeostasis (*13–17*). To overcome this limitation, we recently developed an effective cancer therapy, gene-interfered ferroptosis therapy (GIFT)(*18*), by combining cancer cell-specific knockdown of two iron efflux genes, FPN and LCN2 (*19–23*), with iron nanoparticles (FeNPs) (*17*). GIFT is in fact a gene interference-enhanced ferroptosis. We found that GIFT had wide antitumor activity, high cancer specificity, certain cancer eradication potential, and biosafety in treating variant cancer cells and tumors in mice.

To further improve the therapy, we here develop an updated GIFT named as Ferroptosis Assassinates Tumor (FAST) by knocking down five additional genes related with iron and redox homeostasis (FSP1, FTH1, GPX4, SLC7A11, NRF2)(*24–28*). As a result, we found that FAST showed more excellent antitumor activity than GIFT. Especially, FAST eradicated three different types of tumors (leukemia, colon cancer and lung metastatic melanoma) from over 50 percent of cancer mice, making the mice to survive up to 250 days without tumor relapse. FAST also significantly inhibited growth of spontaneous breast tumor and improved survival in mice. Additionally, FAST showed pan-antitumor activity, high cancer specificity, and in vivo biosafety. In mechanism, FAST destroyed cancer cells by a disaster ferroptosis induced by violent lipid peroxidation resulted from ROS burst that is produced by the FeNPs-released Fe^2+^.

## Results

### Oxidative stress, iron storage suppression-dependent anti-tumor effect

In our recent GIFT (*18*), cancer cells were killed by a combination of DMSA-coated Fe_3_O_4_ nanoparticles (FeNPs) with cancer cell-specific knockdown of two iron efflux genes (FPN and LCN2) (Fig. 1A). The cancer cell-specific knockdown of FPN and LCN2 is realized by controlling expression of miRNAs targeting the two genes with a NF-κB-specific promoter that consists of a NF-κB decoy and a minimal promoter (DMP). Because NF-κB is constitutively over-activated in cancers but not in normal cells (*29, 30*), DMP can drive the effector gene to express only in cancer cells (*31–33*). Because there are variety of ferroptosis-resistance genes in cells, we deduced the antitumor effect of GIFT may be further improved by knocking down other ferroptosis-resistance genes. To verify the speculation, we selected to knock down five new genes distributed in glutathione-dependent pathway (GPX4, SLC7A11, NRF2) (*24–26*), COQ-dependent pathway (FSP1) (*27*), and iron storage (FTH1) (*28*) (Fig. 1A). We designed microRNAs targeting these genes of both human and mouse used to human and mouse cells. respectively (Table S1).

**Fig. 1.**
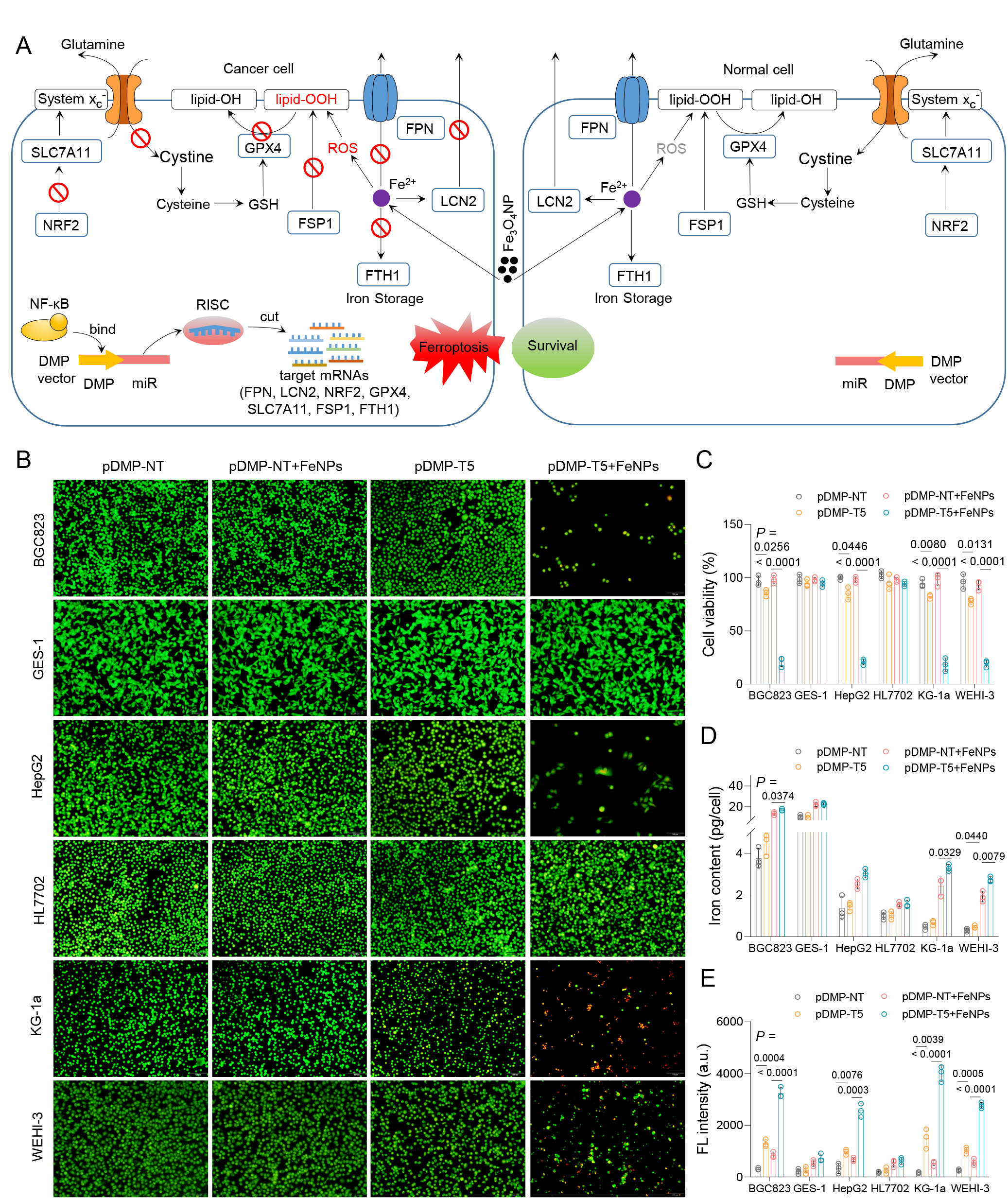
Treatment of tumor and normal cells with pDMP-T5 and FeNPs. Cells (BGC823, GES-1, HepG2, HL7702, KG-1a, WEHI-3) were transfected by DMP vectors overnight, then incubated with or without 50 μg/mL FeNPs for 72 h. (A) Schematic diagram of FAST eradicating tumor. (B) Cells stained by AO&EB. (C) Cell viability. The images and viability of cells incubated with FeNPs for 24 h and 48 h are shown in the figs. 1–6, in which the data of more treatments as controls are shown. Cell viability was detected by the CCK8 assay. (D and E) Iron content (D) and ROS levels (E) of cells incubated with 50 μg/mL FeNPs for 48 h. All values are mean ± s.d. (n = 3 wells). All statistical significance is labeled. pDMP-NT, a plasmid coding an artificial microRNA targeting no transcripts (NT); pDMP-T5, a plasmid coding artificial microRNAs targeting five target (T) genes including FSP1, FTH1, GPX4, SLC7A11, and NRF2.

To demonstrate the potential effects of these genes on cancer cells, we first treated cells with the gene-interfering vectors (GIVs) expressing microRNAs targeting these genes under the control of DMP. The results indicated that both GIVs and FeNPs (50 μg/mL) alone had little effect on cell viability, but the combination of GIVs with FeNPs produced significant time-dependent cytotoxicity to cancer cells (figs. S1– 4). To combine the effect of each genes, we then constructed a DMP-controlled microRNA expression vector that can co-express microRNAs targeting five genes (FSP1, FTH1, GPX4, SLC7A11, and NRF2) (named pDMP-T5, in which T means target). As a result, when combined with FeNPs, pDMP-T5 showed more significant time-dependent cytotoxicity in four cancer cells (HepG2, BGC823, KG-1a, WEHI-3) than GIVs expressing microRNA targeting a single ferroptosis-resistance gene (Fig. 1B and C; figs. S1–4), which was revealed by both acridine orange and ethidium bromide (AO&EB) dual staining and CCK-8 assay. It was found that pDMP-T5 alone also showed a low killing effect to cancer cells, but exerted little effect on the growth of normal cells (HL77202 and GES-1), even combined with FeNPs (figs. S5 and 6). The detection of mRNA level revealed that the expression of five ferroptosis-resistance genes were all significantly knocked down in cancer cells by pDMP-T5 (fig. S7). However, the expression of these target genes was not knocked down by pDMP-T5 in normal cells HL7702 and GES-1 (fig. S7). The measurement of intracellular iron content and total ROS levels indicated that in comparison with the co-treatment of pDMP-NT and FeNPs, the co-treatment of pDMP-T5 and FeNPs slightly increased the intracellular iron contents (Fig. 1D) but significantly increased the ROS levels (Fig. 1E) in four cancer cells (BGC823, HepG2, KG-1a, WEHI-3). Such changes of intracellular iron contents and ROS levels agree with the fact that pDMP-T5 mainly targets to gene producing antioxidants of glutathione (NRF2, SLC7A11, GPX4) and COQ (FSP1), which resist ferroptosis by preventing lipid peroxidation (*27*). Only FTH1, a constituent of ferritin that stores iron (*28*), is targeted by pDMP-T5. The the co-treatment of pDMP-T5 and FeNPs did not changed the intracellular iron contents and ROS levels in normal cells HL7702 and GES-1 (Fig. 1D and E). These results demonstrate that the combination of pDMP-T5 and FeNPs can specifically induce significant ferroptosis of cancer cells by simultaneously producing ROS and knocking down five ferroptosis-resistance genes.

To explore the in vivo anti-tumor effects of pDMP-NT and FeNPs, we then cloned the DMP-T5 fragment into adeno-associated virus (AAV) to prepare a recombinant virus (rAAV-T5). The mouse leukemia cell WEHI-3 was subcutaneously transplanted into BALB/c mice to make tumor-bearing mice. When the average tumor volume of treatment group reaches 150 mm^3^, the WEHI-3 xenograft mice were injected with virus (1×10^10^ vg/mouse) and FeNPs (3 mg/kg) together in the tail vein every two days (fig. S8A). After two injections, only the rAAV-T5+FeNP treatment significantly inhibited tumor growth (fig. S8B–D), and the mice weight remained stable during the study period (fig. S8E). rAAV-T5 or FeNPs alone did not reduce the size and weight of tumor and had little antitumor effect in vivo (fig. S8B–D). To evaluate the distribution of rAAV and the silencing efficiency of target genes in treated mice, we detected the abundance of GIV DNA and mRNA levels of five ferroptosis-resistance genes (FSP1, FTH1, GPX4, SLC7A11, and NRF2) in various tissues by qPCR. The results showed that the GIV DNA distributed in all detected tissues, especially in tumor and liver (fig. S8F). The expression of five target genes were only significantly knocked down in tumor (fig. S8G–K). These data indicated that rAAV-T5 was safe in vivo and had significant antitumor effect when combined with FeNPs.

### The anti-tumor activity of FAST

Many studies have reported that iron overload in cells can cause oxidative stress through the Fenton reaction (*34, 35*). Besides iron storage, iron efflux is also an important pathway to maintain cellular iron homeostasis. The functions of both FPN and LCN2 in cells are related to the efflux of iron ions (*19–23*). In our previous study, we found the expression of the two genes were significant up-regulated when cells were treated with FeNPs (*17*). Once the two genes were knocked down and FeNPs were administered, the iron ion output from cancer cells would be severely hindered, thus leading significant ferroptosis of cancer cells in vitro and in vivo (*18*). Therefore, to further enhance the killing effect on cancer cells, we integrated DMP-miFPN and DMP-miLCN2 (pDMP-T2) into pDMP-T5 to form a stronger ferroptosis-inducing vector (pDMP-T7). As a result, compared with the pDMP-T2 and pDMP-T5, pDMP-T7 resulted in more significant increase of the iron contents and total ROS levels in HepG2 and KG-1a cells post FeNPs administration, but not in HL7702 cell (Fig. 2A and B). The wider anti-tumor effect of pDMP-T7 were further characterized by AO&EB dual staining and CCK-8 assays in four cancer cells (Hepg2, A549, MDA-MB-453, KG-1a) and three normal cells (HL7702, MRC5, MCF-12A). As a result, almost all cancer cells were killed at 72 h upon co-treatment of pDMP-T7 and FeNPs, but normal cells were not significantly affected at any treatment time (figs. S9– 16). Thus, with the synergy of FeNPs, the killing effect of pDMP-T7 was stronger and more efficient than pDMP-T2 and pDMP-T5. Notably, pDMP-T7 alone also had more significant time-dependent cytotoxicity than pDMP-T2 and pDMP-T5, which even caused more than half of MDA-MB-453 to die at 72 h (fig. S16). These results indicated that the inhibition of iron efflux can promote pDMP-T7 to eliminate cancer cells. To verify the pan-anticancer activity of co-treatment of pDMP-T7 and FeNPs, we used the crystal violet staining (CVS) assay to evaluate the antitumor activity of FAST in a variety of cancer cells representing different hematological and solid tumors in human and mouse, including two leukemia cells (KG-1a and WEHI-3) and ten solid tumor cells (A549, PANC-1, SKOV3, MDA-MB-453, BGC-823, KYSE30, B16F10, HepG2, HeLa, and CT26). The results revealed that the combination of pDMP-T7 with FeNPs produced the significant time-dependent killing effects in all detected tumor cells (figs. S17 and 19). To investigate the cancer cell specificity of pDMP-T7, we next treated six normal human and mouse cells (HL7702, MRC5, GES-1, L929, MCF-12A, and NIH-3T3). The results showed that all these cells were survival when treated by pDMP-T7 and FeNPs, even at 72 h (figs. S18 and 19). To further validate the key role of NF-κB activation in FAST, we induced these cells with TNFα (a NF-κB activator) before pDMP-T7 transfection. Upon FeNPs administration, these TNFα-induced cells were also significantly killed by the combination of DMP-T7 and FeNPs (figs. S18 and 19). It suggests that pDMP-T7 is a cancer cell-specific GIV and the combination of DMP-T7 and FeNPs is a cancer cell-specific killer. Thus, we named the co-treatment of DMP-T7 and FeNPs as FAST (Ferroptosis ASsassinates Tumor). The principle of FAST is schematically illustrated in Fig. 1A.

**Fig. 2.**
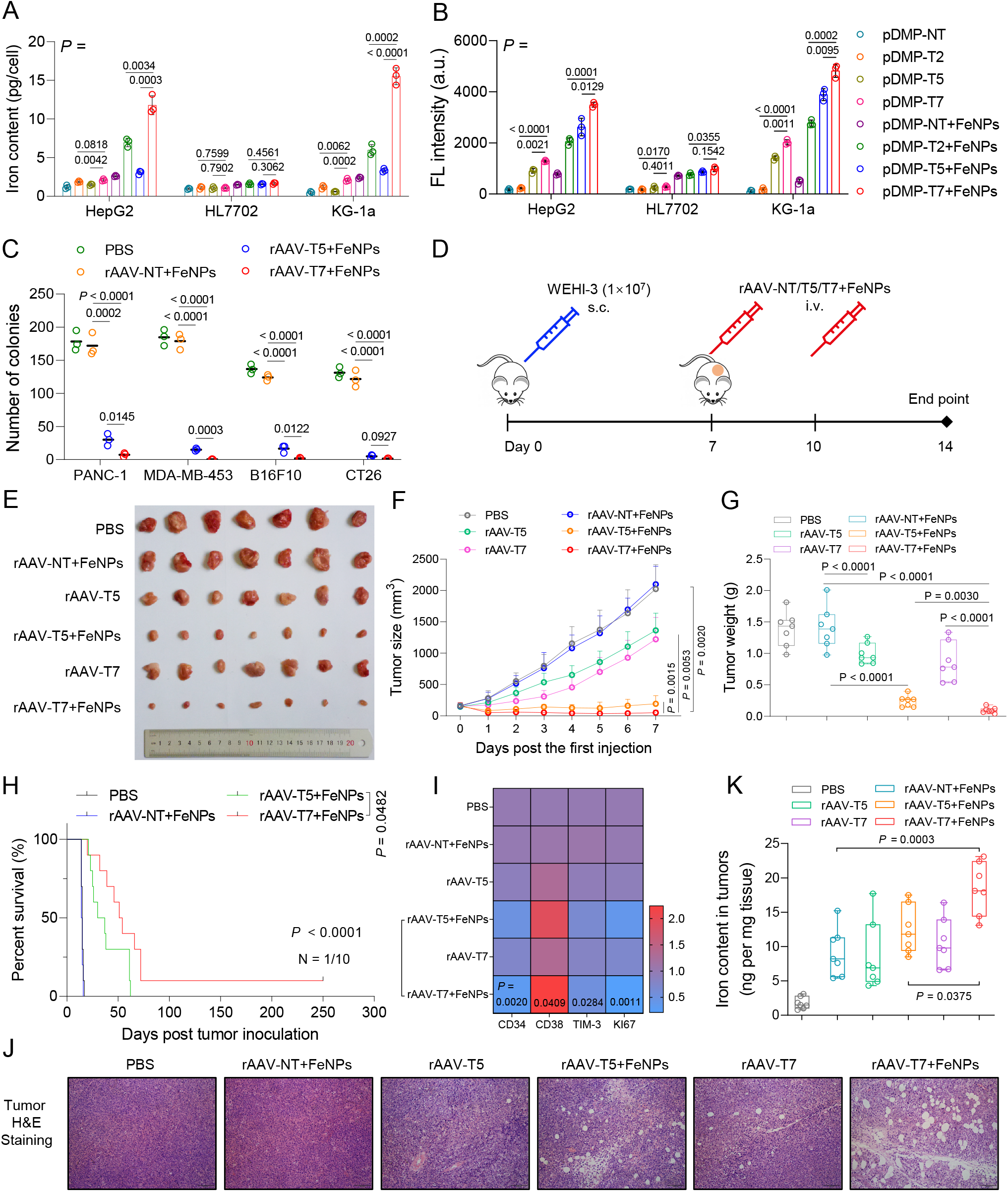
Anti-tumor effects of FAST and other treatments in cancer cells and the WEHI-3 xenograft mice. Cells (HepG2, HL7702, KG-1a) were transfected by pDMP-NT/T2/T5/T7 overnight respectively, and then incubated with or without 50 μg/mL FeNPs for 48 h. (A and B) Comparisons of cellular iron content (A) and total ROS levels (B) post treatment. A and B use a same set of symbols. All data are presented as mean ± s.d (n = 3 wells). pDMP-NT, a plasmid coding an artificial microRNA targeting no transcripts (NT); pDMP-T2, a plasmid coding artificial microRNAs targeting two target (T) genes including FPN and LCN2; pDMP-T5, a plasmid coding artificial microRNAs targeting five target genes including FSP1, FTH1, GPX4, SLC7A11, and NRF2; pDMP-T7, a plasmid coding artificial microRNAs targeting seven target genes including FSP1, FTH1, GPX4, SLC7A11, NRF2, FPN, and LCN2. (C) Quantitative clone formation assays. Cells (PANC-1, MDA-MB-453, B16F10, CT26) were infected with rAAV-NT/T5/T7 at the dose of 1×10^5^ vg per cell for 24 h and then incubated with 50 μg/mL FeNPs for another 48 h. PBS, cells just treated with phosphate buffered saline (PBS) as a control. After treatment, 200 cells were seeded into 6-well plate and cultured until colonies were clearly visible (< 50 cells) (about two weeks). Each treatment was conducted in triplicates. The images of cell clones are shown in fig. S20. rAAV-NT/T5/T7, rAAVs packaged with pDMP-NT, pDMP-T5, and pDMP-T7. (D) Schematics of animal treatment (WEHI-3 xenograft mice). s.c., subcutaneous injection; i.v., intravenous injection. (E) Tumor imaging. (F) Tumor growth curve. All data are presented as mean ± s.d (n = 7 mice). (G) Tumor weight. (n = 7 mice). (H) Kaplan-Meier survival curve (n = 10 mice). The statistical significance was analyzed by the log-rank test. (I) Expression of stemness- and proliferation-related marker genes. The relative expression detected by qPCR (2^−ΔΔt^) was shown by heatmap. The *p* values between rAAV-T5+FeNPs and rAAV-T7+FeNPs are given. (J) Representative H&E-stained section of tumors. (K) Iron content in tumors of mice in different treatment groups (n =7 mice). By these comparative treatments of cells and mice, the treatment of pDMP/rAAV-T7+FeNPs that shows the best anti-tumor effect was defined as FAST, referring to Ferroptosis ASsassinates Tumor.

To explore the in vivo anti-tumor effects, we also cloned DMP-T7 into AAV to prepare a recombinant virus (rAAV-T7). All rAAVs were tested by infecting four cancer cell lines (PANC-1, MDA-MB-453, B16F10, CT26). Consistent with the cell viability experiments, rAAV-T7 showed a more significant anti-tumor activity than rAAV-T5. The colony formation assays showed that FAST could nearly eradicate the colony formation capability and tumorigenicity of various tumor cells (Fig. 2C; fig. S20). Next, the WEHI-3 xenograft mice model was used to evaluate the anti-tumor effects of FAST (Fig. 2D). The tumor-bearing mice were intravenously administered twice with PBS, rAAV-NT+FeNPs, rAAV-T5, rAAV-T5+FeNPs, rAAV-T7, and rAAV-T7+FeNPs, respectively (1×10^10^ vg/mouse and 3 mg/kg FeNPs, n = 7). The results indicated that rAAV-T7+FeNPs was the most active therapeutic intervention in delaying tumor progression (Fig. 2E–G) and extending survival (Fig. 2H) in all treatment groups. Additionally, the expression of genes as stemness (CD34, CD38, TIM-3) and proliferation (Ki67)-related markers in tumors were detected (Fig. 2I). The heatmap showed that CD38 was upregulated but CD34, TIM-3, and Ki-67 were down-regulated in tumors by FAST. It suggested that the stemness and proliferation of tumor were significantly inhibited by FAST, which explained the reason for the shrinkage of tumor. The hematoxylin and eosin (H&E) staining of tumor tissue slices further proved the superior antitumor activity of FAST (Fig. 2J). In general, FAST showed the better antitumor activity than rAAV-T5+FeNPs (Fig. 2E–J). To further characterize the treatment, we detected the iron content, abundance of rAAV DNA and mRNA of seven target genes (FSP1, FTH1, GPX4, NRF2, SLC7A11, FPN and LCN2) and RELA in various tissues. Compared with rAAV-NT+FeNPs, FAST significantly increased the iron content in tumors (Fig. 2K), but not in other tissues (fig. S21A). The rAAV DNA distributed in all detected tissues, especially in liver and tumor (fig. S21B). Seven target genes were only knocked down significantly in tumor (fig. S21C) and RELA was only highly expressed in tumor (fig. S21D). These data showed the cancer specificity and NF-κB activity dependence of FAST therapy. Additionally, all mice showed no significant pathological tissue damage in major organs (heart, liver, spleen, lung, and kidney) (fig. S22A) and lose of body weight (fig. S22B) during the treatment, indicating the biosafety of FAST.

To further explore the safety of FAST, we also treated the healthy BALB/c mice with FAST. The healthy mice were randomly divided into two groups and injected three times with PBS and rAAV-T7+FeNPs (5×10^10^ vg/mouse and 3 mg/kg FeNPs, n = 5) via tail vein, respectively (fig. S23A). Compared the PBS group, the FAST treatment produced no significant effect on the body weight (fig. S23B), blood biochemical markers (white blood cells, red blood cells, platelet, and hemoglobin), hepatotoxicity markers (alanine aminotransferase, aspartate aminotransferase, and alkaline phosphatase), and kidney injury markers (blood urea nitrogen, creatinine, and uric acid) (fig. S23C) of mice. The FAST treatment also produced no significant effect on the external appearance and size (fig. S24A) and tissue structure (fig. S24B) of the main organs of mice. Especially, compared the PBS group, the FAST treatment showed no significant effect on the weight of liver and spleen of mice (fig. S24C). Additionally, although the rAAV distributed in all detected tissues (fig. S24D), it did not affect the expression of all target genes and NF-κB/RELA in these tissues (fig. S24E). Therefore, the used dose of FAST reagents had high biosafety, which was then used in following animal experiments.

### Ferroptosis dependence of FAST antitumor activity

Ferroptosis was primarily characterized by condensed mitochondrial membrane densities and corresponding volume reduction than normal mitochondria (*36*). To clarify the form of cell death, we observed the morphology of HepG2 (a solid tumor cell) and KG-1a (a hematological cell) at 48 h post FAST treatment by Transition electron microscopy (TEM). In the two cells, both the FAST and erastin (a typical ferroptosis inducer) treatments induced the mitochondria shrinking and mitochondrial membrane density increase (fig. S25); however, FeNPs (50 μg/mL) alone did not change the mitochondrial morphology like DMSO (fig. S25). The results directly demonstrated that the mechanism underlying FAST was ferroptosis. Ferroptosis is a unique form of cell death that is different from apoptosis, autophagy, and necrosis. Its occurrence depends on severe lipid peroxidation relying on ROS generation and iron overload (*2*). As revealed by the above experiments, the FAST treatment resulted in the significant increase of both iron contents and total ROS level in HepG2 and KG-1a (Fig. 2A and B). To further confirm whether the catastrophic ROS accumulation was related with lipid peroxidation, we next detected the lipid peroxidation using a lipid oxidation indicator, C11-BODIPY. The results indicated that the FAST treatment most significantly elevated the lipid peroxidation from 24 h to 72 h in HepG2 (fig. S26; Fig. 3A and B). The elevated lipid peroxidation could be reverted by the ferroptosis inhibitors (Fer1, DFO, and NAC), but not by apoptosis inhibitor (ZVAD), necroptosis inhibitor (Nec1s), and autophagy inhibitor (BA1) (fig. S26; Fig. 3C). Moreover, the elevated lipid peroxidation could be most significantly reverted by the co-treatment of DFO, Fer1, NAC (DFN) (fig. S26; Fig. 3C). The similar lipid peroxidation was also obtained in other cancer cells (A549, KG-1a, MDA-MB-453, PANC-1) (Fig. 3A; figs. S27 and 28). However, the FAST treatment showed no similar lipid peroxidation effect on normal cells (MCF-12A, MRC-5, HL7702, GES-1) (Fig. 3A; figs. S29 and 30). Nevertheless, when these normal cells were induced by TNFα (a NF-κB stimulator), the FAST treatment produced the similar lipid peroxidation in these cells (figs. S29 and 30). These data indicated the cancer cell specificity and NF-κB activity dependence of FAST (figs. S26-30). As a positive control, erastin also induced significant lipid peroxidation in cancer cells; however, it also induced significant lipid peroxidation in normal cells (fig. S25; figs. S27-30), indicating its strong side effects on normal cells. The ferroptosis dependence of cancer cell-specific antitumor activity of FAST was also confirmed by the CCK-8 assay of cell proliferation (Fig. 3D) and colony formation assays (Fig. 3E; fig. S31). In two assays, each single ferroptosis inhibitor (DFO, Fer1, NAC) recovered limited cell viability and a few colonies, whereas cocktail of multiple ferroptosis inhibitor (DFN) rescued most cell viability and colonies (Fig. 3D and E; fig. S31). This is consistence with the mechanism that FAST inhibits multiple anti-ferroptosis pathways by knocking down key genes responsible for these pathways. Interestingly, the autophagy inhibitor BA1 rescued about 30% of viability of the FAST-treated cancer cells (Fig. 3D and E; fig. S31), suggesting that autophagy partially contributed to the antitumor activity of FAST. This was also confirmed by existence of autophagosomes in the FAST-treated cancer cells (fig. S32; fig. S33). This is consistence with the reports that lipid peroxidation can induce autophagosome formation and ferroptosis is closely related with autophagy (*37*). Finally, we checked ferroptosis dependence of FAST in vivo using the WEHI-3 xenograft tumor model. The tumor-bearing mice were intravenously administered three times with rAAV-NT+FeNPs, rAAV-T7+FeNPs, and rAAV-T7+FeNPs+NAC (5×10^10^ vg/mouse Virus, 3 mg/kg FeNPs, n = 10), respectively (Fig. 3F). The ferroptosis inhibitor NAC (1 g/L) was added in the drinking water. As a result, the FAST treatment significantly inhibited tumor growth and prolonged survival (60% survival rate) (Fig. 3G and H); however, NAC significantly offset the FAST effects (Fig. 3G and H), indicating that FAST inhibits tumor by ferroptosis in vivo, consistent with the in vitro results. Importantly, we found that 60% FAST-treated mice lived tumor-free up to 250 days without tumor relapse, indicating that tumors were eradicated from these mice by FAST.

**Fig. 3.**
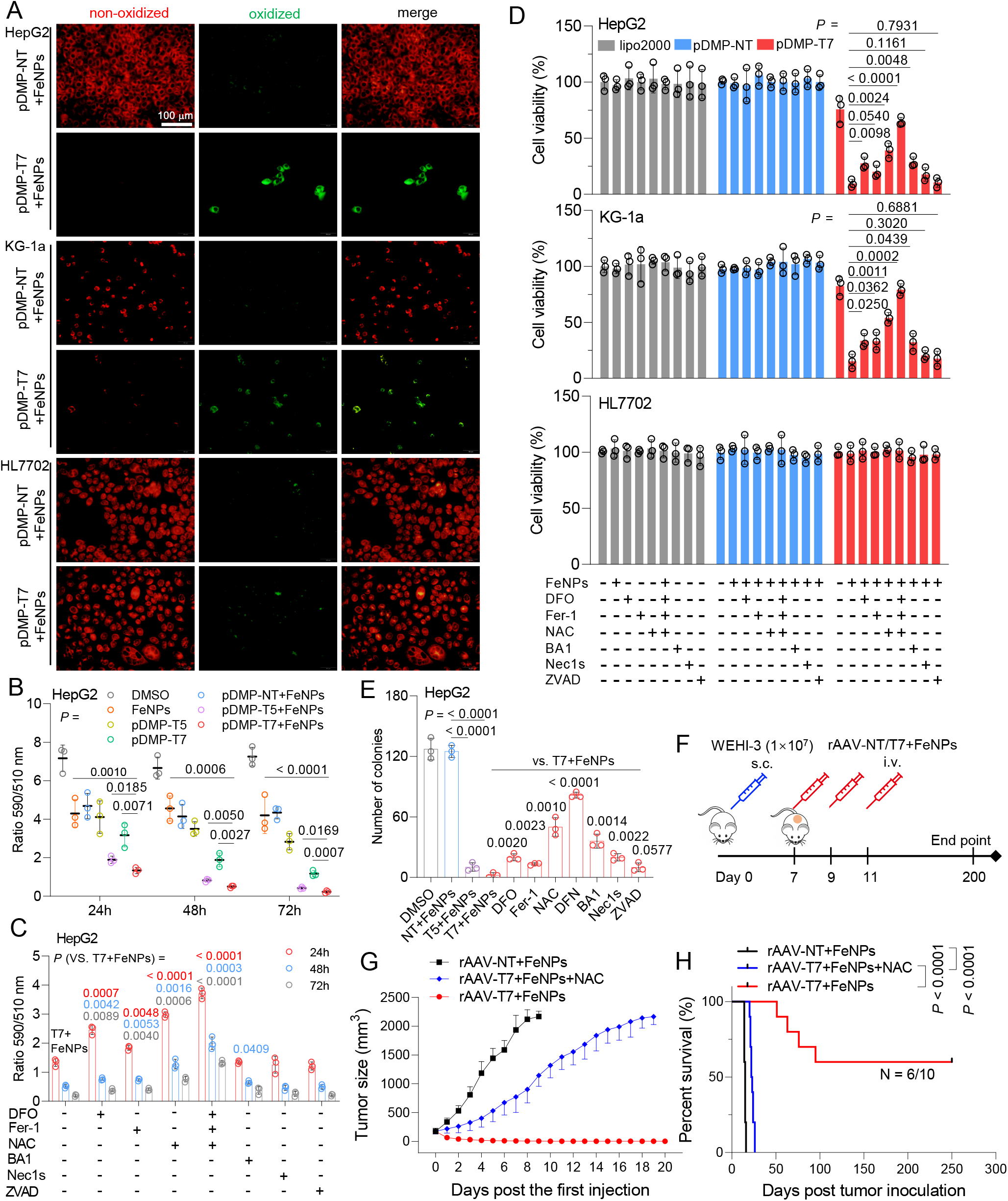
FAST-induced cancer cell death by ferroptosis. (A) Lipid ROS imaging. Lipid ROS production was detected by C11-BODIPY and imaged by fluorescence microscope. Red, reduced dye; green, oxidized dye. Only the representative images of HepG2, KG-1a, and HL7702 that were incubated with 50 μg/mL FeNPs for 72 h post pDMP-NT/T7 transfection are shown here. More detailed pictures (including 5 cancer cells and 4 normal cells) and quantified results are shown in figs. 26-30. (B) and (C) The quantified data of lipid peroxidation in HepG2 at 24, 48, and 72 h. The lipid peroxidation in cells were determined by quantitating the fluorescence intensities analyzed by ImageJ software and calculating the ratio of intensity in 590 to 510 channels. Fer-1, Ferrostain-1 (1 μM); DFO, deferoxamine (100 μM); NAC, N-acetylcysteine (1 mM); DFN (mixture of 1 μM Fer1, 100 μM DFO, and 1 mM NAC); BA1, Bafilomycin A1 (1 nM); Nec1s, Necrostatin-1s (10 μM); ZVAD, ZVAD-FMK (50 μM). (D) Cell viability of HepG2, HL7702 and KG-1a when co-incubated with FeNPs and cell death inhibitors for 72 h post plasmid transfection overnight. (E) Quantitative clone formation assays of the treated HepG2 when exposed to cell death inhibitors. Each treatment was conducted in triplicates. The images of cell clones are shown in fig. 31. (F) Schematics of animal treatment (WEHI-3 xenograft mice). s.c., subcutaneous injection; i.v., intravenous injection. (G) Tumor growth curve (n = 10 mice). (H) Kaplan-Meier survival curve (n = 10 mice). The statistical significance was analyzed by the log-rank test.

### Antitumor activity of FAST in variant tumors

To further confirm in vivo antitumor effect of FAST therapy, we established a colon cancer model by subcutaneously injecting CT26 cells in BALB/c mice. When the tumor volume reached around 150 mm^3^, the tumor-bearing mice were intravenously administered three times with rAAV-NT/T7+FeNPs (5×10^10^ vg/mouse Virus, 3 mg/kg FeNPs) (Fig. 4A). As a result, the body weight of mice remained stable during the treatment (Fig. 4B); however, the tumor size was remarkably shrunk by FAST (Fig. 4C and D). The tumor weight was significantly decreased (Fig. 4E) and the survival was greatly improved by FAST (Fig. 4F). Excitingly, 60% of mice survived and lived over 250 days without tumor relapse (Fig. 4F). Further detections of tissues revealed that the the iron content was significantly increased by FAST in tumor (fig. S34A), rAAV-T7 distributed in all detected tissues (fig. S34B), the expression of seven target genes was significantly knocked down by FAST only in tumors (fig. S34C) due to NF-κB expression only in tumors (fig. S34D), and the expression of stemness and proliferation markers was significantly decreased in tumors by the FAST treatment (fig. S34E). In addition, there was no significant splenomegaly and hepatomegaly (fig. S35A-C) and changes of tissue structure (fig. S35D), blood biochemical markers (fig. S35E), and liver and kidney injury markers (fig. S35F) after the FAST treatment in this tumor model, further indicating the safety of FAST.

**Fig. 4.**
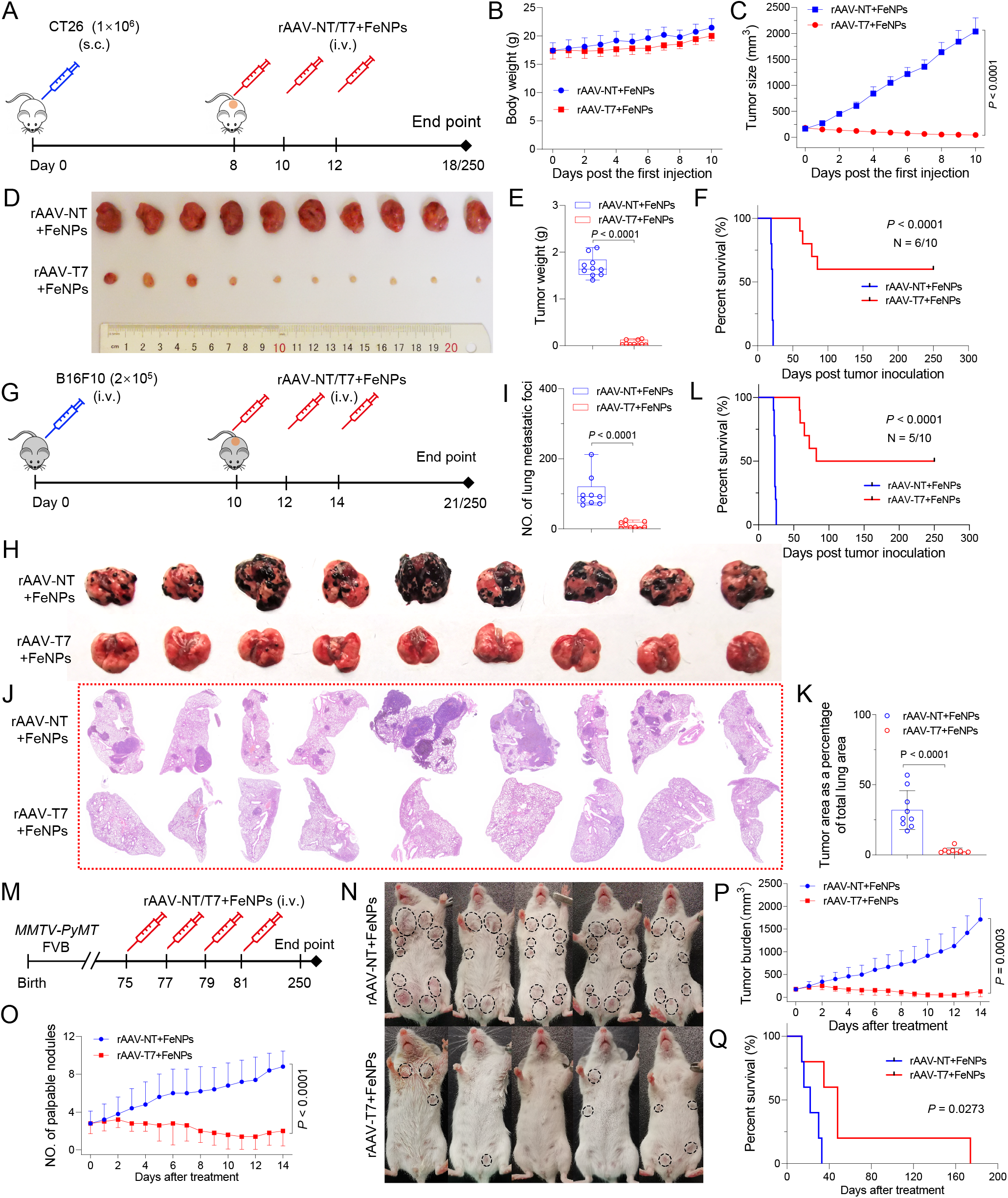
The antitumor effects of FAST in three tumor models of mice. (A-F) The in vivo antitumor effects of FAST in the colon cancer model of mice. The model was constructed by subcutaneously injecting the CT26 cells (n = 10 mice). (A) Schematics of animal treatment. s.c. subcutaneous injection; i.v., intravenous injection. (B) Average body weight. (C) Tumor growth curve. (D) Tumor imaging. (E) Tumor weight. Data are shown as mean± s.d (n = 10 mice). (F) Kaplan-Meier survival curve (n = 10 mice). The statistical significance was analyzed by the log-rank test. (G-L) The in vivo antitumor effects of FAST in the pulmonary metastatic melanoma model. The model was constructed by intravenously injecting the B16F10 cells. (G) Schematics of animal treatment. (H) Lung imaging. (I) Quantified B16F10 lung metastatic-like tumor foci. Data are shown as mean± s.d (n = 9 mice). (J) H&E-stained lung section imaging. (K) Tumor area as a percent of total lung area. Data are shown as mean± s.d (n = 9 mice). (L) Kaplan-Meier survival curve (n = 10 mice). The statistical significance was analyzed by the log-rank test. (M-Q) The in vivo antitumor effects of FAST in spontaneous breast cancer model (n = 5 mice). (M) Schematics of animal treatment. (N) Representative image showing gross appearance of tumors. Dotted-line circles demarcate palpable mammary tumor nodules. (O) Comparison of the number of palpable tumor nodules. (P) Comparison of total tumor burden. Tumor burden was calculated by summating the volume of every tumor nodule per mouse. Data are presented as mean± s.d. (n = 5 mice). (Q) Kaplan-Meier survival curve. The statistical significance was analyzed by the log-rank test.

To determine whether the FAST therapy could similarly treat metastatic tumors, we then treated a lung metastatic melanoma model made by intravenously injecting B16F10 cells in C57BL/6J female mice with FAST. The tumor-bearing mice were intravenously administered three times with rAAV-NT/T7+FeNPs (5×10^10^ vg/mouse Virus, 3 mg/kg FeNPs) (Fig. 4G). The results indicated that the FAST treatment significantly reduced tumor burden of lung assessed by both metastatic-like tumor foci number (Fig. 4H and I) and tumor area as a percentage of total lung area (Fig. 4J and K). The FAST treatment prevented the body weight loss in compared with those treated by rAAV-NT+FeNPs (fig. S36A). The FAST treatment significantly decreased splenomegaly (fig. S36B) and the weight of spleen (fig. S36C) and lung (fig. S36D). Tyrosinase-related protein 1 (Tyrp1) is a melanocyte-specific biomarker whose expression in the lung is restricted to B16F10, which provides a quantitative measure of tumor development (*38*). The FAST treatment significantly decreased the tyrosinase expression in lung tissue (fig. S36E) and the melanin in lung extract (fig. S36F). Although the rAAV distributed in all tissues (fig. S36G), the FAST treatment produced no significant changes in tissue structure (fig. S36H), liver weight (fig. S36I), blood biochemical markers (fig. S36J), and liver and kidney injury markers (fig. S36K), indicating the safety of FAST. Most importantly, the FAST treatment significantly improved the survival of mice, making 50% of mice surviving and living over 250 days without tumor relapse (Fig. 4L).

To explore whether FAST have the antitumor activity in spontaneous cancer model, we treated the *MMTV-PyMT* transgenic mice with FAST, which spontaneously develops metastatic breast cancer. Once the tumor burden reached ~150 mm^3^, the mice were randomly divided into two groups and injected rAAV-NT+FeNPs and rAAV-T7+FeNPs via tail vein every other day, respectively (Fig. 4M). The results exhibited a significant reduction in overall tumor burden and a delay in the development of palpable tumor nodules compared with the control mice (Fig. 4N–P; fig. S37A). Histologic analyses exhibited the FAST treatment effectively blocked the occurrence of lung metastasis and tumor invasion (fig. S37B-D). Tissue detection revealed that rAAV-T7 distributed in all tissues (fig. S37E) and the tumor iron content was increased by the FAST treatment (fig. S37F). Similarly, the FAST treatment produced no significant changes in other tissue structures (fig. S37G) and body weight (fig. S37H). Critically, the overall survival was significantly improved by the FAST treatment in compared with the control, making one mouse (n=1/4) surviving and living up to 174 days (Fig. 4Q). To further confirm the anti-tumor effect of FAST in this spontaneous cancer model, we performed a new animal experiment with more mice, in which two groups each contained 8 mice were respectively treated with PBS (the control group) and FAST (the treatment group) (fig. S38A) at the same time points (Fig. 4M). However, a new group of 8 mice was treated with FAST one week ahead (the prevention group) (fig. S38A), by which we expect to know if FAST could prevent cancer because no tumor can be detectable at that time. As a result, we found that the the treatment group obtained the similar anti-tumor effect (fig. S38B-G). Additionally, the prevention group indeed obtained the prevention effect (fig. S38B-G), making 37.5% mice (n=3/8) survive to 100 days without tumors.

To further explore the anti-tumor effect of FAST, we finally established a new cancer model by subcutaneously injected human hepatoma cell HepG2 to nude mice (fig. S39A). We then treated the mice with PBS, rAAV-NT+FeNPs, and rAAV-T7+FeNPs, respectively. The results revealed that FAST also showed significantly anti-tumor effect on this tumor model, indicating that FAST can also inhibit the human tumor growth in living body (fig. S39B-D). However, the anti-tumor effect of FAST on this model is not as marked as those on other models. Because nude mouse has no adaptive immunity, we thus deduced that the immunity may also contribute to the anti-tumor effect of FAST as a recent study reported that ferroptosis is immunogenic in mice (*39*). To verify the deduction, we first re-challenged the the mice survived up to 250 days in above experiments with cancer cells same as or different from the cancer cells used to established tumor models (fig. S40A-C). As a result, we found that mice died with the same survival days (fig. S40A-C), suggesting that no immunogenicity was elicited by ferroptosis of tumors in mice with three different tumors. To further verify this conclusion, we performed a new animal experiment, in which the mice were subcutaneously injected with PBS, CT26 cells, and the FAST-pretreated CT26 cells, respectively (fig. S40D). As a result, the mice injected with CT26 cells died at 21 days post injection; however, no mice injected with FAST-pretreated CT26 cells grew tumors until 40 days post injection (fig. S40E). We then re-challenged the PBS- and FAST-pretreated CT26 cell-injected mice with fresh CT26 cells. As a result, we found that the mice in two groups died at the same time (fig. S40E), also indicating that no immunogenicity was intrigued by ferroptosis of cancer cells. In these experiments, the body weight of mice was not obviously affected by these treatments, showing no toxicity of FAST-pretreated cancer cells to mice (fig. S40F). Altogether, these data indicated that FAST has no significant immunogenicity, suggesting that FAST inhibits tumors just by ferroptosis in mechanism. Because many studies reported that ferroptosis triggers the release of pro-inflammatory damage-associated molecular patterns (DAMPs) related to the activation of innate immune system such as surface-exposed calreticulin (CRT) and secreted high mobility group box-1 protein (HMGB1)(*40–43*), we finally detected the release of these DAMPs in the FAST-treated cells to further explore the immunogenicity of FAST. The results indicate that CRT was exposed on surface of the FAST-treated cancer cells (fig. S41A and B) and HMGB1 was released in the culture medium of the FAST-treated cancer cell (fig. S41C). However, compared with ferroptosis inducer erastin, FAST induced low level of CRT and HMGB1 (fig. S41). The low DAMPs inducement may result in non-immunogenicity of FAST in mice. It should be noted that erastin also strongly induced DAMPs in normal cells but FAST did not (fig. S41B and C). The immunogenicity of FAST should be further explored in later.

## Conclusion

We here develop a new cancer therapy, ferroptosis-assassinating tumor (FAST), by combining iron oxide nanoparticles with cancer-selective knockdown of seven ferroptosis-resistance genes. FAST induced significant ferroptosis in variety of cancer cells but had little effect on normal cells. FAST also showed notable anti-tumor efficacy in five kinds of tumor-bearing mice with high biosafety. FAST realizes its anti-tumor efficacy by a gene interference-enhanced ferroptosis in mechanism.

## Materials and methods

### Vector construction

The universal miRNA expression vector pDMP-miR was optimized based on pCMV-miR which was previously constructed by our laboratory by replacing the CMV promoter with a decoy minimal promoter (DMP). DMP, a chemically synthesized NF-κB-specific promoter, contains a NF-κB response sequence (5′-GGG AAT TTC CGG GGA CTT TCC GGG AAT TTC CGG GGA CTT TCC GGG AAT TTC C-3′) and a minimal promoter sequence (5′-TAG AGG GTA TAT AAT GGA AGC TCG ACT TCC AG-3′). The miRNAs targeting human or murine FSP1, FTH1, GPX, NRF2, SLC7A11 were designed into two sets for the degradation of different targeted gene regions by BLOCK-iT™ RNAi Designer (https://rnaidesigner.thermofisher.com/rnaiexpress/), respectively (Table S1). Oligonucleotide pairs synthesized by Sangon Biotech (Shanghai, China) (Table S2) were denatured and then annealed to obtain double-stranded DNA (dsDNA), which were then linked with the linear pDMP-miR vector cleaved with BsmBI. The generated miRNA expression vectors targeting the FSP1, FTH1, GPX, NRF2 and SLC7A11 were named as pDMP-miFSP1, pDMP-miFTH1, pDMP-miGPX4, pDMP-miNRF2 and pDMP-miSLC7A11, respectively. Due to the differences in gene sequences between species, pDMP expression vectors for human and mouse were specifically constructed. By cell viability screening figs. 1-6), two set of miRNAs for target gene, of which the more cytotoxic was selected for co-expressing vector construction. Then, the plasmid co-expressing miRNAs targeting the FSP1, FTH1, GPX, NRF2 and SLC7A11 was obtained by In-fusion cloning (Takara), named pDMP-T5. The DMP-FPN and DMP-LCN2 fragment were amplified from pDMP-T2 and ligated into pDMP-T5 to get pDMP-T5. pDMP-T2 was obtained from our previous work. AS a negative control vector, the miR-NT fragment was synthesized according to the sequence of plasmid pcDNA™ 6.2-GW/EmGFP-miR-Neg and inserted into pDMP-miR, named pDMP-NT.

The DMP-T5 (DMP-miFSP1-DMP-miFTH1-DMP-miGPX4-DMP-miNRF2-DMP-miSLC7A11) and DMP-T7 (DMP-miFSP1-DMP-miFTH1-DMP-miGPX4-DMP-miNRF2-DMP-miSLC7A11-DMP-miFPN-DMP-miLCN2) sequences were cleaved from pDMP-T5 and pDMP-T7, respectively. By using the MluI (upstream) and AfeI (downstream) restriction sites, the cleaved fragments were cloned into pAAV-MCS (VPK-410, Stratagene) to construct the pAAV-T5 and pAAV-T7 vectors, respectively. The DMP-NT fragment was also copied from pDMP-NT and inserted into pAAV-MCS to get pAAV-NT vector. Vectors were detected with PCR amplification and verified by DNA sequencing.

### Nanoparticles, cells and culture

The DMSA-coated Fe_3_O_4_ magnetic nanoparticle (FeNPs) were provided by the Biological and Biomedical Nanotechnology Group of the State Key Lab of Bioelectronics, Southeast University, Nanjing, China. This nanoparticle was characterized by our previous study (*17*). Cells used in this research included KG-1a (human acute myeloid leukaemia cells), WEHI-3 (mouse acute mononuclear leukaemia cells), HEK-293T (human fetal kidney cells), HepG2 (human liver cancer cells), A549 (human lung cancer cells), HeLa (human cervical cancer cells), SKOV3 (human ovarian cancer cells), PANC-1 (human pancreatic cancer cells), MDA-MB-453 (human breast cancer cells), B16F10 (mouse melanoma cells), BGC-823 (Human gastric adenocarcinoma cells), KYSE30 (human esophageal carcinoma cells), CT26 (mouse colon cancer cells), HL7702 (human normal hepatocytes), L929 (mouse fibroblast), NIH-3T3 (mouse embryonic fibroblast), MRC5 (human embryonic fibroblasts), GES-1 (human gastric mucosal epithelial cells), and MCF-12A (human breast epithelial cells). KG-1a, SKOV3, MCF-12A cell lines were acquired from American Type Culture Collection (ATCC). WEHI-3, HEK-293T, HepG2, A549, HeLa, PANC-1, MDA-MB-453, B16F10, BGC-823, KYSE30, CT26, HL7702, L929, NIH-3T3, MRC-5, and GES-1 cell lines were obtained from the cell resource center of Shanghai Institutes for Biological Sciences, Chinese Academy of Sciences. Leukaemia cell lines, KG-1a, and WEHI-3, were cultured in Iscove’s Modified Dulbecco’s Medium (IMEM) (Gibco). HEK-293T, HepG2, HeLa, PANC-1, MDA-MB-453, B16F10, MRC-5, L929 and NIH-3T3 cells were cultured in Dulbecco’s Modified Eagle Medium (DMEM) (Gibco). A549, SKOV-3, BGC-823, KYSE30, CT26, MCF-12A and HL7702 cells were cultured in Roswell Park Memorial Institute (RPMI) 1640 medium (Gibco). All media were supplemented with 10% fetal bovine serum (HyClone), 100 units/mL penicillin (Thermo Fisher), and 100 μg/mL streptomycin (Thermo Fisher). Cells were incubated at 37 °C in a humidified incubator containing 5% CO_2_.

### AO&EB staining

Cell transfection was performed with plasmids using Lipofectamine 2000 (Thermo Fisher Scientific) according to the manufacturer’s instruction. Briefly, cells (1×10^5^) were seeded into 24-well plates overnight, then transfected with 500 ng of various plasmids including pDMP-NT, pDMP-miFSP1, pDMP-miFTH1, pDMP-miGPX4, pDMP-miNRF2, pDMP-miSLC7A11, pDMP-T2, pDMP-T5 and pDMP-T7. The mouse and human cells were transfected with vectors targeting to mouse and human genes, respectively. The transfected cells were cultured overnight and then incubated with or without 50 μg/mL FeNP for 24 h, 48 h, 72 h. After FeNPs administration, all cells were stained with acridine orange and ethidium bromide (AO&EB, Solarbio) following the manufacturer’s instruction. Live cells will appear uniformly green, necrotic cells will stain orange. Cells were imaged under a fluorescence microscope (IX51, Olympus) to observe numbers of live and dead cells.

### Cytotoxicity assays

Cell viability was estimated according to Cell counting Kit-8 (CCK-8) and Crystal Violet staining (CVS) assays carried out independently. Cell suspension (5000 cells/well) were dispensed in a 96-well plate and cultured at 37°C in 5% CO_2_ overnight. Cells were then transfected overnight with various pDMP-miR expression vectors (200 ng) including all the vectors used for the above AO&EB staining to screen the more cytotoxic miRNAs. The transfected cells were cultured overnight and then incubated with or without 50 μg/mL FeNP for 24 h, 48 h, 72 h.

For viability assays for testing various inhibitors, cells (5000 cells/well) were also seeded in 96-well plate and cultured overnight. Then cells were transfected with 200 ng pDMP-NT/T7 overnight. Next, the treated cells were co-incubated with 50 μg/mL FeNPs and the indicated inhibitors including ferrostatin-1 (Fer-1) (Sigma, SML0583) (1 μM), deferoxamine (DFO) (ApexBio, B6068) (100 μM), N-acetylcysteine (NAC) (Sigma, A9165) (1 mM), DFN (mixture of 1 μM Fer1, 100 μM DFO and 1 mM NAC), ZVAD-FMK (ZVAD) (ApexBio, A1902) (50 μM), Necrostatin-1s (Nec1s) (BioVision, 2263-1) (10 μM), Bafilomycin A1 (BA1) (Sigma, B1793) (1 nM), and cells were cultured for another 72 h. After FeNPs administration, 10 μL of CCK-8 solution (Biosharp) was added to each test well and incubate the plate for 1 h in the incubator. The absorbance at 450 nm was measured using a microplate reader (BioTek).

For the CVS assay, 12 cancer cell lines and 6 normal cell lines were measured post FAST therapy at 24, 48 and 72 h. Normal cell lines were first induced with or without 10 ng/mL TNFα (Sigma-Aldrich) for 1 h before pDMP-NT/T7 transfection. Then all cell lines were incubated with 50 μg/mL FeNPs for 24, 48 and 72 h. The treated cells were fixed with 4% paraformaldehyde for 10 minutes, and rinse twice with distilled water for 3 min. Next, cells were stained with crystal violet (CV, Sigma, C6158) at 0.02% (w/v) for 5 min at room temperature. Each treatment was conducted in triplicates. After washing and observation, CV was eluted with 0.1 M sodium citrate in 50% (v/v) ethanol, and recorded absorbance values at 585 nm to obtain quantitative results.

### ROS measurement

Cells were treated with FeNPs as previously described. Briefly, cells (5000 cells/well) were also seeded in 96-well plate and cultured for overnight growth. Cells were then transfected with 200 ng of various plasmids including pDMP-NT, pDMP-T2, pDMP-T5, and pDMP-T7. The transfected cells were cultured overnight and then incubated with or without 50 μg/mL of FeNPs for another 24 h, 48 h, 72 h. To evaluate cellular total ROS levels, the treated cells were stained with 2′,7′-dichlorodihydrofluorescein diacetate (DCFH-DA, 10 μM) at 48 h using the Reactive Oxygen Species Assay Kit (Beyotime) according to the manufacturer’s instructions. ROS changes indicated by fluorescence shift was detect using a fluorescence microplate reader (BioTek). The lipid peroxidation of 5 cancer cells (HepG2, A549, PANC-1, KG-1a, MDA-MB-453) and 4 normal cells (GES-1, HL7702, MCF-12A, MRC5) post FAST therapy were observed at 72 h. Normal cell lines were first induced with or without 10 ng/mL TNFα for 1 h before plasmids transfection. Cells were exposed to erastin (10 μM) (Sigma-Aldrich) for 8 h as a positive control. To exam the effect of cell death inhibitors on cellular lipid peroxidation, the transfected HepG2 cell was co-incubated with FeNPs and one of the indicated inhibitors (Fer-1, DFO, NAC, DFN, ZVAD, Nec1s, BA1) for 24 h, 48 h, and 72 h. After FeNPs administration, the lipid peroxidation of HepG2 was measured by BODIPY® 581/591 C11 using the Image-iT™ Lipid Peroxidation Kit (Thermo Fisher) and imaged by fluorescence microscope (IX51, Olympus) using traditional 590 nm and 510 nm emission filters with a 40X objective, respectively. The lipid peroxidation in cells were determined by quantitating the fluorescence intensities analyzed with ImageJ 1.51j8 software and calculating the ratio of intensity in 590 to 510 channels.

### Iron content measurement

The average intracellular iron content and iron content of tissues were measured by ICP-MS (Agilent Technologies 7700, USA). The measurement procedure can be summarized as the following: (i) Cells were treated with FeNPs as previously described. Intracellular iron was determined 48 h post FeNP administration. Cells were washed with PBS, digested, counted and precipitated by centrifugation. Tissues were weighed and transferred to the 5 mL centrifuge tubes. (ii) The cell precipitation or tissues were then added a certain amount of 65% nitric acid and heated for complete digest. (iii) Iron standard solutions (GSB 04-1726-2004, Beijing) with different concentrations (0, 0.1, 0.2, 0.5, 1, 2, and 5 μg/mL) were prepared to establish the standard curve for ICP-MS measurement. The intracellular iron content was reported as average iron content per cell. The iron content of tissue was reported as iron content per mg tissue. Each experiment was repeated in triplicates.

### Virus preparation

HEK293T cells were seeded into 75 cm^2^ flasks at a density of 5×10^6^ cells per flask and cultivated for overnight. Cells were then co-transfected with two helper plasmids (pHelper and pAAV-RC; Stratagene) and one of the pAAV plasmids (pAAV-NT, pAAV-T5, pAAV-T7) using Lipofectamine 2000 according to the manufacturer’s instructions. Cells were cultured for another 72 h. The cells and media were collected and kept at −80 °C overnight. The cells and media were then incubated in 37 °C water bath for 2 h. This freeze-thaw process was totally repeated three times. The 1/10 volume of pure chloroform was added to the cell lysate and the mixture was vigorously shaken at 37 °C for 1 h. The mixture was added NaCl to a final concentration of 1 M and shaken until NaCl dissolved. The mixture was centrifuged at 15,000 revolutions per minute (rpm) at 4 °C for 15 min and the supernatant was collected. The supernatant was added PEG8000 at a final concentration of 10% (w/v) and shaken until PEG8000 dissolved. The mixture was centrifuged at 15,000 rpm at 4 °C for 15 min. The supernatant was discarded and the pellet was dissolved into PBS. DNase and RNase were added to a final concentration of 1 μg/mL to the dissolved pellet. The mixture was incubated at room temperature for 30 min. The mixture was extracted once with chloroform (1:1 volume) and the aqueous layer that contained the purified virus was transferred to a new tube. Titers of AAVs were determined by qPCR using the primers AAV-F/R (Table S3). Quantified viruses were aliquoted and kept at −80 °C for later use. The obtained viruses were named as rAAV-NT, rAAV-T5, rAAV-T7.

### Clone formation assay

Cells were infected with the viruses including rAAV-NT, rAAV-T5, rAAV-T7 at the dose of 1×10^5^ vg per cell for 24 h and then incubated with 50 μg/mL FeNPs for another 48 h. For inhibitor groups, the infected cells were co-incubated with FeNPs and indicated inhibitors for 48 h. After treatment, cells were pipetted into 6-well plates at a density of 200 cells/well and cultured until colonies were clearly visible (> 50 cells). The test well was fixed with 4% paraformaldehyde (Sangon Biotech, China), stained with crystal violet (CV, Sigma, C6158) at 0.02% (w/v) for 5 min at room temperature and counted using a light microscope. Each treatment was conducted in triplicates.

### Transmission Electron Microscope (TEM)

KG-1a and HepG2 cells were treated with DMSO (48 h, 0.1%), FeNPs (48 h, 50 μg/mL), Erastin (8 h, 10 μM), pDMP-T7+FeNPs (plasmid transfection overnight, then incubated with 50 μg/mL FeNPs for 48 h), respectively. Then cells were fixed in in suspension with 2.5% glutaraldehyde in 0.1 M calcodyate buffer (pH 7.4) after harvesting, overnight at 4 °C. Cell pellets were post-fixed with 2% osmium tetraoxide dissolved in 0.1 M cacodylate and 1.5% potassium ferrocyanide at room temperature for 1 h. Cells were dehydrated with a graded series of ethanol, and embedded in epoxy Epon812 (EMS, 14120). Then the areas containing cells were cut into ultrathin sections and stained with uranyl acetate and lead citrate and observed on transmission electron microscope (Carl Zeiss microscopy GmbH, Jena, Germany).

### MDC staining

Dansylcadaverine (MDC) is a fluorescent pigment and an eosinophilic stain, which is usually used to detect the formation of autophagosomes. Cells were transfected with pDMP-T7 overnight and then incubated with or without FeNPs for 72 h. Rapamycin (RAP, 500 nM) (Aladdin) was added to culture medium for 12 h as a positive control to induce cell autophagy. Bafilomycin A1 (BA1, 1 nM), a typical autophagy inhibitor, was co-incubated with FeNPs (72 h, 50 μg/mL), RAP (12 h, 500 nM) and Erastin (8 h, 10 μM), respectively. After incubation, the cells were stained using MDC stain kit (Solarbio) and DAPI (Sigma-Aldrich) following the manufacturer’s instruction. Then, the treated cells were rinsed three times with PBS and observed using a fluorescence microscope (Olympus IX51, Tokyo, Japan).

### Quantitative PCR

Total RNA was isolated from cell lines at 48 h post incubation with FeNPs or mouse tissues using TRIzol™ (Invitrogen) according to the manufacturer’s protocol. The complementary DNA (cDNA) was generated using the FastKing RT kit (Tiangen) according to the manufacturer’s instruction. The genomic DNA (gDNA) was extracted from various tissues of mice using the TIANamp Genomic DNA Kit (TIANGEN). Amplification for the genes of interest from cDNA and gDNA was performed by quantitative PCR (qPCR) using the Hieff qPCR SYBR Green Master Mix (Yeasen). The primers used for qPCR are shown in the Table S3. Triplicate samples per treatment were evaluated on an ABI Step One Plus (Applied Biosystems). Relative mRNA transcript levels were compared to the GADPH internal reference and calculated as relative quantity (RQ) according to the following equation: RQ = 2^−ΔΔCt^. Virus DNA abundance were normalized to the GADPH internal reference and calculated according to the following equation: RQ = 2^−ΔCt^. Tyrosinase in lung of the pulmonary metastatic melanoma model was detected by qPCR using the primers Murine Tyrp1 F/R. The results are normalized to mouse GAPDH and analyzed by 2^−ΔCt^ method. All qPCR primers were verified specific using melting curve analysis and were listed in Table S3. All experiments were performed in triplicates.

### Animal treatments

Four-week-old female BALB/c, four-week-old male BALB/c nude mice and ten-week-old C57BL/6J female mice were purchased from the Changzhou Cavens Laboratory Animal Co. Ltd (China). Female *MMTV-PyMT* transgenic mice (FVB/N) were purchased from The Jackson Laboratory (#002374). All animal experiments in this study followed the guidelines and ethics of the Animal Care and Use Committee of Southeast University (Nanjing, China). Tumor growth was monitored by volume measurement with calipers. Tumor volumes were calculated using formula V = (ab^2^)/2, where a is the longest diameter and b is the shortest diameter. The mice were euthanized when the tumor size reached 2000 mm^3^ or the body weight loss was greater than 20% of the predosing weight. Various tissues (including heart, liver, spleen, lung, kidney, and tumor tissues) were collected for further analysis. Five animal models were performed.

Three batches of animal experiments were performed in the WEHI-3 xenografted model on BALB/c mice. WEHI-3 xenografts were generated by subcutaneously transplantation with 1×10^7^ WEHI-3 cells into inner thighs. Mice were bred for 7 days for tumor formation. In the first batch of animal experiment, the tumor-bearing mice of WEHI-3 cell were randomly divided into six treatment groups (PBS, FeNPs, rAAV-NT, rAAV-NT+FeNPs, rAAV-T5, rAAV-T5+FeNPs; n=7). The mice were intravenously injected twice every two days with PBS, FeNPs, rAAV-NT, rAAV-NT+FeNPs, rAAV-T5, rAAV-T5+FeNPs, respectively. The dose of viruses and FeNPs were 1×10^10^ vg/mouse and 3 mg/kg body weight, respectively. To simplify the drug administration, rAAV and FeNPs were mixed together and intravenously injected to mice one time in this batch of animal experiment. Tumor size and mice weight were measured every day. Mice were euthanized and photographed on the seventh day post FeNPs injection.

In the second batch of animal experiment, the tumor-bearing mice of WEHI-3 cell were randomly divided into six treatment groups (PBS, rAAV-NT+FeNPs, rAAV-T5, rAAV-T5+FeNPs, rAAV-T7, rAAV-T7+FeNPs; n = 7). The mice were intravenously injected twice every two days with PBS, rAAV-NT+FeNPs, rAAV-T5, rAAV-T5+FeNPs, rAAV-T7, rAAV-T7+FeNPs. The dose of viruses and FeNPs were 1×10^10^ vg/mouse and 3 mg/kg body weight, respectively. Virus and FeNPs were injected as mixture. Tumor size and mice weight were measured every day. Mice were euthanized and photographed on the seventh day post FeNPs injection. Various tissues including heart, liver, spleen, lung, kidney, and tumor were harvested for H&E analysis, virus DNA, gene expression and iron content detection. To further study the survival rate of mice using this therapy, the Kaplan-Meier method was used. The tumor-bearing mice of WEHI-3 cell were randomly divided into four treatment groups (PBS, rAAV-NT+FeNPs, rAAV-T5+FeNPs, rAAV-T7+FeNPs; n = 10). Mice were euthanized and recorded when the tumor size reached 2000 mm^3^.

In the third batch of animal experiment, the tumor-bearing mice of WEHI-3 cell were randomly divided into three treatment groups (rAAV-NT+FeNPs, rAAV-T7+FeNPs+NAC, rAAV-T7+FeNPs; n = 10). The mice were intravenously injected three times every other day with rAAV-NT+FeNPs, rAAV-T7+FeNPs+NAC, rAAV-T7+FeNPs, respectively. The dose of viruses and FeNPs were 5×10^10^ vg/mouse and 3 mg/kg body weight, respectively. Virus and FeNPs were injected as mixture. Mice in the groups of rAAV-T7+FeNPs+NAC were administered NAC in their drinking water at 1 g/L. The body weight of the mice and tumor size were monitored daily. The Kaplan-Meier method was used to analyze the mice survival over time. Mice were euthanized and recorded when the tumor size reached 2000 mm^3^.

For the safety assessment, ten BALB/c female mice were randomly divided into two treatment groups (PBS and rAAV-T7+FeNPs; n = 5). The mice were intravenously administered three times every other day with PBS and rAAV-T7+FeNPs, respectively. The dose of virus and FeNPs were 5×10^10^ vg/mouse and 3 mg/kg body weight, respectively. Virus and FeNPs were injected as mixture. The body weight of the mice was monitored daily. Mice were euthanized on the seventh day post first injection. Blood and serum samples from each group were collected for routine blood test and serum biochemical parameter detection. Various tissues including heart, liver, spleen, lung, and kidney were photographed and harvested for H&E analysis.

Two batches of animal experiments were performed in the CT26 xenografted model on BALB/c mice. CT26 xenografts were generated on BALB/c mice by subcutaneously transplantation with 1×10^6^ CT26 cells into inner thighs. The mice were bred for 8 days for tumor formation. In the first batch of animal experiment, the tumor-bearing mice were randomly divided into four treatment groups (rAAV-NT+FeNPs-1, rAAV-NT+FeNPs-2, rAAV-T7+FeNPs-1, rAAV-T7+FeNPs-2; n = 10). Mice were intravenously administered every other day three times with rAAV-NT+FeNPs (rAAV-NT+FeNPs-1 and rAAV-NT+FeNPs-2) and rAAV-T7+FeNPs (rAAV-T7+FeNPs-1 and rAAV-T7+FeNPs-2), respectively. The dose of viruses and FeNPs were 5×10^10^ vg/mouse and 3 mg/kg body weight, respectively. Virus and FeNPs were injected as mixture. The body weight of the mice and tumor size were monitored daily. Mice in groups (rAAV-NT+FeNP-1, rAAV-NT+FeNPs-1; n = 10) were euthanized and photographed on the tenth day post first injection. Blood and serum samples from each group were collected for routine blood test and serum biochemical parameter detection. Various tissues including heart, liver, spleen, lung, kidney, and tumor were harvested for H&E analysis and virus DNA and iron content detection. The weight of liver and spleen were weighted and recorded. The mice in the other two groups (rAAV-NT+FeNPs-2 and rAAV-T7+FeNPs-2; n = 10) were used for survival study. Mice were euthanized when the tumor size reached 2000 mm^3^. In the second batch of animal experiment, the mice were randomly divided into three groups (PBS, rAAV-NT+FeNPs, rAAV-T7+FeNPs; n = 6). CT26 cells were pre-treated with rAAV-NT+FeNPs and rAAV-T7+FeNPs for 24 h, respectively. After infection, the treated cells (1×10^6^) were collected and subcutaneously transplantation into inner thighs. The control group was simultaneously injected with 100 μl of PBS into inner thighs. 40 days post tumor inoculation, the CT26 cells (1×10^6^) were subcutaneously transplantation into inner thighs again in PBS and rAAV-T7+FeNPs groups. The body weight of the mice and tumor size were monitored daily. The Kaplan-Meier method was used to analyze the mice survival over time. Virus and FeNPs were injected as mixture. The body weight of the mice and tumor size were monitored daily.

Pulmonary metastatic melanoma model was established on C57BL/6J female mice by intravenously injection 2×10^5^ B16F10 cells. The mice were bred for 10 days for tumor formation. The tumor-bearing mice were randomly divided into four treatment groups (rAAV-NT+FeNPs-1, rAAV-NT+FeNPs-2, rAAV-T7+FeNPs-1, rAAV-T7+FeNPs-2). Mice were intravenously administered every other day three times with rAAV-NT+FeNPs (rAAV-NT+FeNPs-1 and rAAV-NT+FeNPs-2) and rAAV-T7+FeNPs (rAAV-T7+FeNPs-1 and rAAV-T7+FeNPs-2), respectively. The dose of viruses and FeNPs were 5×10^10^ vg/mouse and 3 mg/kg body weight, respectively. Virus and FeNPs were injected as mixture. The body weight of the mice was monitored daily. Mice in groups (rAAV-NT+FeNPs-1 and rAAV-T7+FeNP-1; n = 9) were euthanized and photographed on the eleventh day post first injection. Blood and serum samples from each group were collected for routine blood test and serum biochemical parameter detection. Lung, liver and spleen of mice were weighted and recorded. Various tissues including heart, liver, spleen, lung, and kidney were harvested for H&E analysis, virus DNA, gene expression and iron content detection. The mice in the other two groups (rAAV-NT+FeNP-2 and rAAV-T7+FeNP-2; n = 10) were used for survival study. Mice were euthanized when the body weight loss was greater than 20% of the predosing weight.

A HepG2 tumor xenograft model was established on BALB/c nude mice by subcutaneously transplantation with 1×10^7^ HepG2 cells into inner thighs. The mice were bred for 12 days for tumor formation. When the average tumor size reached ~150 mm^3^, the tumor-bearing mice were randomly divided into three treatment groups (PBS, rAAV-NT+FeNPs, rAAV-T7+FeNPs; n = 6). Mice were intravenously administered every other day four times with 100 μl of PBS, rAAV-NT+FeNPs and rAAV-T7+FeNPs, respectively. The dose of viruses and FeNPs were 5×10^10^ vg/mouse and 3 mg/kg body weight, respectively. The body weight and tumor size of the mice was monitored daily. The Kaplan-Meier method was used to analyze the mice survival over time.

Two batches of animal experiments were performed in female *MMTV-PyMT* transgenic mice, which was spontaneous breast cancer model. In the first batch of animal experiment, starting at 10 weeks after birth, the number of tumor nodules and volume of tumor burden (sum volume of all tumor nodules) in mice were observed and measured daily. When the tumor burden reached ~150 mm^3^, the mice were randomly divided into two groups (rAAV-NT+FeNPs and rAAV-T7+FeNPs; n = 5). Mice were intravenously administered every other day four times with rAAV-NT+FeNPs and rAAV-T7+FeNPs, respectively. The dose of viruses and FeNPs were 5×10^10^ vg/mouse and 3 mg/kg body weight, respectively. Virus and FeNPs were injected as mixture. The body weight of the mice was also monitored daily. Two weeks later, one mouse with the biggest tumor burden in each group was selected for dissection and observation. Various tissues including heart, liver, spleen, lung, kidney, and tumor were photographed and harvested for H&E analysis and virus DNA and iron content detection. The Kaplan-Meier method was used to analyze the mice survival over time. Mice were euthanized and recorded when the tumor burden reached 2000 mm^3^. In the second batch of animal experiment, the female *MMTV-PyMT* transgenic mice were randomly divided into three treatment groups (rAAV-NT+FeNPs, rAAV-T7+FeNPs-1, rAAV-T7+FeNPs-2; n = 8). At 11 weeks after birth (average tumor burden reached ~150 mm^3^), mice were intravenously administered every other day four times with rAAV-NT+FeNPs and rAAV-T7+FeNPs, respectively. Here, we set up an early treatment group (rAAV-T7+FeNPs-2), in which mice were injected with rAAV-T7+FeNPs four times every other day from 10 weeks after birth. The dose of viruses and FeNPs were 5×10^10^ vg/mouse and 3 mg/kg body weight, respectively. Virus and FeNPs were injected as mixture. The body weight, the number of tumor nodules and volume of tumor burden in mice were observed and measured daily. The Kaplan-Meier method was used to analyze the mice survival over time.

In addition, the surviving mice of pulmonary metastatic melanoma model, WEHI-3 xenografted model, and CT26 xenografted model whose tumors were eradicated by FAST were performed the re-challenge experiments at 250 days. The surviving mice (n = 5) of pulmonary metastatic melanoma model were intravenously injection 2×10^5^ B16F10 cells. The body weight of the mice was monitored daily. The mice were euthanized when the tumor size reached 2000 mm^3^ (for WEHI-3 and CT26 mice) or the body weight loss was greater than 20% of the predosing weight (for B16F10 mice). The Kaplan-Meier method was used to analyze the mice survival over time. The surviving mice of WEHI-3 xenografted model (n=6) were randomly divided into two groups, one group (n = 3) was subcutaneously transplantation with 1×10^7^ WEHI-3 cells into inner thighs, the other (n = 3) subcutaneously transplantation with 1×10^6^ CT26 cells into inner thighs. Similarly, the surviving mice of CT26 xenografted model (n = 6) were randomly divided into two groups, one group (n=3) was subcutaneously transplantation with 1×10^6^ CT26 into inner thighs, the other (n = 3) subcutaneously transplantation with 1×10^7^ WEHI-3 cells into inner thighs. The body weight and tumor size of the mice was monitored daily. The mice were euthanized when the tumor size reached 2000 mm^3^. The Kaplan-Meier method was used to analyze the mice survival over time.

### Hematoxylin and eosin (H&E) Staining

Tissues including heart, liver, spleen, lung, kidney and tumor were dissected, embedded in paraffin, sectioned, and stained with H&E using routine methods. Briefly, tissues were resected and fixed overnight in 4% paraformaldehyde solution (Sangon Biotech, China) at room temperature overnight. Subsequently, fixed specimens were embedded in paraffin, divided into 5 μm-thick sections, and then stained with hematoxylin staining solution (C0107, Beyotime) and eosin staining solution (C0109, Beyotime). The prepared slides were photographed by a microscope (IX51, Olympus).

### Detection of Immunogenic Cell Death Biomarkers

The HepG2, CT26, HL7702, and NIH-3T3 cells were seeded in the 48-well culture vessel (1×10^5^ cells) and incubated at 37°C in a CO_2_ incubator overnight. The cells were transfected 500 ng of plasmid (pDMP-NT or pDMP-T7) and incubated with 50 μg/mL of FeNPs for 24 h. As a positive control of ferroptosis, cells were exposed to erastin (5 μM) for 24 h. Then, the cells were received a standard immunofluorescence protocol to assess the surface-exposure of calreticulin (CRT). Briefly, the treated cells were incubated with primary rabbit anti-CRT antibody (abcam, ab92516) overnight at for 4 °C, followed by incubating with an Alexa Fluor 594-conjugated goat anti-rabbit IgG antibody (abcam, ab150080) for 1 h at room temperature. Then, the cells were further stained with DAPI (Sigma-Aldrich) for 20 min at room temperature before visualized under fluorescence microscope. To determine the release of high mobility group box-1 protein (HMGB1), 100 μL of supernatant from each treated well were collected and measured using an ELISA kit according to the manufacturer instruction. The optical density (OD) value was read at 450 nm on a Microplate reader.

### Statistical analysis

All data are presented as means values ± standard deviation (SD), and statistical analysis and graphs were performed through GraphPad Prism 8.0 software. Statistical differences between two groups were determined using two-tailed Student’s t-test. Comparisons of three or more groups were determined by one-way or two-way analysis of variance (ANOVA) with Tukey’s or Sidak’s multiple comparison test when appropriate. The Kaplan–Meier method was used to analyze the differences in animal survival and the P value was calculated by the log-rank test. Differences at p < 0.05 were considered statistically significant.

## References

1. D. Tang, R. Kang, T. V. Berghe, P. Vandenabeele, G. Kroemer, The molecular machinery of regulated cell death. Cell research 29, 347–364 (2019).

2. B. Hassannia, P. Vandenabeele, T. Vanden Berghe, Targeting Ferroptosis to Iron Out Cancer. Cancer Cell 35, 830–849 (2019).

3. M. H. Gao et al., Ferroptosis is an autophagic cell death process. Cell research 26, 1021–1032 (2016).

4. D. A. Stoyanovsky et al., Iron catalysis of lipid peroxidation in ferroptosis: Regulated enzymatic or random free radical reaction? Free Radical Bio Med 133, 153–161 (2019).

5. J. Li et al., Ferroptosis: past, present and future. Cell Death Dis 11, 88 (2020).

6. H. Z. Feng, B. R. Stockwell, Unsolved mysteries: How does lipid peroxidation cause ferroptosis? Plos Biol 16, e2006203 (2018).

7. S. J. Dixon et al., Ferroptosis: an iron-dependent form of nonapoptotic cell death. Cell 149, 1060–1072 (2012).

8. A. A. Salahudeen et al., An E3 ligase possessing an iron-responsive hemerythrin domain Is a regulator of iron homeostasis. Science 326, 722–726 (2009).

9. A. A. Vashisht et al., Control of iron homeostasis by an iron-Regulated ubiquitin ligase. Science 326, 718–721 (2009).

10. S. Y. Shen et al., Sensitive tumour detection and classification using plasma cell-free DNA methylomes. Nature 563, 579–583 (2018).

11. S. E. Kim et al., Ultrasmall nanoparticles induce ferroptosis in nutrient-deprived cancer cells and suppress tumour growth. Nature nanotechnology 11, 977–985 (2016).

12. L. Gao et al., Intrinsic peroxidase-like activity of ferromagnetic nanoparticles. Nature nanotechnology 2, 577–583 (2007).

13. G. Gao, J. Li, Y. Zhang, Y. Z. Chang, Cellular Iron Metabolism and Regulation. Advances in experimental medicine and biology 1173, 21–32 (2019).

14. V. Trujillo-Alonso et al., FDA-approved ferumoxytol displays anti-leukaemia efficacy against cells with low ferroportin levels. Nature nanotechnology 14, 616–622 (2019).

15. C. Zhang, F. Zhang, Iron homeostasis and tumorigenesis: molecular mechanisms and therapeutic opportunities. Protein Cell 6, 88–100 (2015).

16. D. H. Manz, N. L. Blanchette, B. T. Paul, F. M. Torti, S. V. Torti, Iron and cancer: recent insights. Ann. N. Y. Acad. Sci. 1368, 149–161 (2016).

17. Y. Liu, J. Wang, Effects of DMSA-coated Fe3O4 nanoparticles on the transcription of genes related to ion and osmosis homeostasis. Toxicol. Sci. 131, 521–536 (2013).

18. J. Gao, T. Luo, J. Wang, Gene interfered-ferroptosis therapy for cancers. Nat. Commun. 12, 5311 (2021).

19. G. Montosi et al., Autosomal-dominant hemochrom-atosis is associated with a mutation in the ferroportin (SLC11A3) gene. J. Clin. Invest. 108, 619–623 (2001).

20. M. Sabelli et al., Human macrophage ferroportin biology and the basis for the ferroportin disease. Hepatology 65, 1512–1525 (2017).

21. D. L. Zhang et al., Erythrocytic ferroportin reduces intracellular iron accumulation, hemolysis, and malaria risk. Science 359, 1520–1523 (2018).

22. J. Yang et al., An iron delivery pathway mediated by a lipocalin. Mol. Cell 10, 1045–1056 (2002).

23. S. Ziegler et al., Lipocalin 24p3 is regulated by the Wnt pathway independent of regulation by iron. Cancer Genet Cytogen 174, 16–23 (2007).

24. M. A. Badgley et al., Cysteine depletion induces pancreatic tumor ferroptosis in mice. Science 368, 85–89 (2020).

25. W. S. Yang et al., Regulation of ferroptotic cancer cell death by GPX4. Cell 156, 317–331 (2014).

26. M. Dodson, R. Castro-Portuguez, D. D. Zhang, NRF2 plays a critical role in mitigating lipid peroxidation and ferroptosis. Redox Biol 23, 101107 (2019).

27. K. Bersuker et al., The CoQ oxidoreductase FSP1 acts parallel to GPX4 to inhibit ferroptosis. Nature 575, 688–692 (2019).

28. Y. Tian et al., FTH1 Inhibits Ferroptosis Through Ferritinophagy in the 6-OHDA Model of Parkinson’s Disease. Neurotherapeutics 17, 1796–1812 (2020).

29. R. Sen, D. Baltimore, Multiple nuclear factors interact with the immunoglobulin enhancer sequences. Cell 46, 705–716 (1986).

30. Y. Ben-Neriah, M. Karin, Inflammation meets cancer, with NF-κB as the matchmaker. Nat. Immunol. 12, 715–723 (2011).

31. D. Wang et al., Control the intracellular NF-kappaB activity by a sensor consisting of miRNA and decoy. Int. J. Biochem. Cell Biol. 95, 43–52 (2018).

32. D. Wang, W. Dai, J. Wang, A cell-specific nuclear factor-κB-activating gene expression strategy for delivering cancer immunotherapy. Hum. Gene Ther. 30, 471–484 (2018).

33. W. Dai, J. Wu, D. Wang, J. Wang, Cancer gene therapy by NF-κB-activated cancer cell-specific expression of CRISPR/Cas9 targeting to telomere. Gene ther. 27, 266–280 (2020).

34. M. Li et al., State-of-the-art iron-based nanozymes for biocatalytic tumor therapy. Nanoscale Horizons 5, 202–217 (2020).

35. L. M. Bystrom, M. L. Guzman, S. Rivella, Iron and reactive oxygen species: friends or foes of cancer cells? Antioxid Redox Signal 20, 1917–1924 (2014).

36. H. Wang, C. Liu, Y. Zhao, G. Gao, Mitochondria regulation in ferroptosis. Eur J Cell Biol 99, 151058 (2020).

37. B. Zhou et al., Ferroptosis is a type of autophagy-dependent cell death. Semin Cancer Biol 66, 89–100 (2020).

38. P. H. Lizotte et al., In situ vaccination with cowpea mosaic virus nanoparticles suppresses metastatic cancer. Nat. Nanotechnol. 11, 295–303 (2016).

39. I. Efimova et al., Vaccination with early ferroptotic cancer cells induces efficient antitumor immunity. J Immunother Cancer 8, e001369 (2020).

40. Y. Yu et al., The ferroptosis inducer erastin enhances sensitivity of acute myeloid leukemia cells to chemotherapeutic agents. Mol Cell Oncol 2, e1054549 (2015).

41. Q. Wen, J. Liu, R. Kang, B. Zhou, D. Tang, The release and activity of HMGB1 in ferroptosis. Biochem Biophys Res Commun 510, 278–283 (2019).

42. B. Yu, B. Choi, W. Li, D. H. Kim, Magnetic field boosted ferroptosis-like cell death and responsive MRI using hybrid vesicles for cancer immunotherapy. Nat. Commun. 11, 3637 (2020).

43. F. Ye et al., HMGB1 regulates erastin-induced ferroptosis via RAS-JNK/p38 signaling in HL-60/NRAS(Q61L) cells. Am. J. Cancer Res. 9, 730–739 (2019).

