## Supplementary information for "Ferroptosis assassinates tumor (FAST)"

**Table S1. Target sequence of miRNA.**

| Name | miRNA target (5′-3′) | Name | miRNA target (5′-3′) |
|---|---|---|---|
| Human FSP1-1 | TCCACTTTATTTGTTTGTTTG | Human NRF2-1 | CCAGTGGATCTGCCAACTACT |
| Human FSP1-2 | CTTTATTTGTTTGTTTGTTTA | Human NRF2-2 | ATGCCCTCACCTGCTACTTTA |
| Mouse FSP1-1 | CCACGCTGGCCTGCATGCCAA | Mouse NRF2-1 | TACTCCCAGGTTGCCCACATT |
| Mouse FSP1-2 | GCTGGCCTGCATGCCAATGTT | Mouse NRF2-2 | GCAAGTTTGGCAGGAGCTATT |
| Human FTH1-1 | TTGTCTTTGAGGTCTTGGGAT | Human SLC7A11-1 | TGTTTGCTGTCTCCAGGTTAT |
| Human FTH1-2 | GGCTATCTTCCAGATTCCTTA | Human SLC7A11-2 | TGGCAGTTGCTGGGCTGATTT |
| Mouse FTH1-1 | GAAGCTGGCATGGCAGAATAT | Mouse SLC7A11-1 | CTGGGTGGAACTGCTCGTAAT |
| Mouse FTH1-2 | GGCAGAATATCTCTTTGACAA | Mouse SLC7A11-2 | AGACCTCTATAGTCTTCTAAA |
| Human GPX4-1 | CCGCCTTTGCCGCCTACTGAA | Human FPN | ATCATGTAAGCTGCAGGTAGA |
| Human GPX4-2 | CACCGTCTCTCCACAGTTCCT | Human LCN2 | AGTTCACGCTGGGCAACATTA |
| Mouse GPX4-1 | GGAGCCCATTCCTGAACCTTT | Mouse FPN | GCAAATCAGTGAGTCTGTATA |
| Mouse GPX4-2 | GGACGCCAAAGTCCTAGGAAA | Mouse LCN2 | AGGACTCAACTCAGAACTTGA |
| NT | GTCTCCACGCGCAGTACATTT | | |

**Table S2. Oligonucleotides sequences used to construct miRNA expression vector.**

| Name | Sequence (5′-3′) |
|---|---|
| Human miFSP1-F-1 | TGCTGCAAACAAACAAATAAAGTGGAGTTTTGGCCACTGACTGACTCCACTTTTTGTTTGTTTG |
| Human miFSP1-R-1 | CCTGCAAACAAACAAAAAGTGGAGTCAGTCAGTGGCCAAAACTCCACTTTATTTGTTTGTTTGC |
| Human miFSP1-F-2 | TGCTGTAAACAAACAAACAAATAAAGGTTTTGGCCACTGACTGACCTTTATTTTTGTTTGTTTA |
| Human miFSP1-R-2 | CCTGTAAACAAACAAAAATAAAGGTCAGTCAGTGGCCAAAACCTTTATTTGTTTGTTTGTTTAC |
| Mouse miFSP1-F-1 | TGCTGTTGGCATGCAGGCCAGCGTGGGTTTTGGCCACTGACTGACCCACGCTGCTGCATGCCAA |
| Mouse miFSP1-R-1 | CCTGTTGGCATGCAGCAGCGTGGGTCAGTCAGTGGCCAAAACCCACGCTGGCCTGCATGCCAAC |
| Mouse miFSP1-F-2 | TGCTGAACATTGGCATGCAGGCCAGCGTTTTGGCCACTGACTGACGCTGGCCTATGCCAATGTT |
| Mouse miFSP1-R-2 | CCTGAACATTGGCATAGGCCAGCGTCAGTCAGTGGCCAAAACGCTGGCCTGCATGCCAATGTTC |
| Human miFTH1-F-1 | TGCTGATCCCAAGACCTCAAAGACAAGTTTTGGCCACTGACTGACTTGTCTTTGGTCTTGGGAT |
| Human miFTH1-R-1 | CCTGATCCCAAGACCAAAGACAAGTCAGTCAGTGGCCAAAACTTGTCTTTGAGGTCTTGGGATC |
| Human miFTH1-F-2 | TGCTGTAAGGAATCTGGAAGATAGCCGTTTTGGCCACTGACTGACGGCTATCTCAGATTCCTTA |
| Human miFTH1-R-2 | CCTGTAAGGAATCTGAGATAGCCGTCAGTCAGTGGCCAAAACGGCTATCTTCCAGATTCCTTAC |
| Mouse miFTH1-F-1 | TGCTGATATTCTGCCATGCCAGCTTCGTTTTGGCCACTGACTGACGAAGCTGGTGGCAGAATAT |
| Mouse miFTH1-R-1 | CCTGATATTCTGCCACCAGCTTCGTCAGTCAGTGGCCAAAACGAAGCTGGCATGGCAGAATATC |
| Mouse miFTH1-F-2 | TGCTGTTGTCAAAGAGATATTCTGCCGTTTTGGCCACTGACTGACGGCAGAATCTCTTTGACAA |
| Mouse miFTH1-R-2 | CCTGTTGTCAAAGAGATTCTGCCGTCAGTCAGTGGCCAAAACGGCAGAATATCTCTTTGACAAC |
| Human miGPX4-F-1 | TGCTGTTCAGTAGGCGGCAAAGGCGGGTTTTGGCCACTGACTGACCCGCCTTTCGCCTACTGAA |
| Human miGPX4-R-1 | CCTGTTCAGTAGGCGAAAGGCGGGTCAGTCAGTGGCCAAAACCCGCCTTTGCCGCCTACTGAAC |
| Human miGPX4-F-2 | TGCTGAGGAACTGTGGAGAGACGGTGGTTTTGGCCACTGACTGACCACCGTCTCCACAGTTCCT |
| Human miGPX4-R-2 | CCTGAGGAACTGTGGAGACGGTGGTCAGTCAGTGGCCAAAACCACCGTCTCTCCACAGTTCCTC |
| Mouse miGPX4-F-1 | TGCTGAAAGGTTCAGGAATGGGCTCCGTTTTGGCCACTGACTGACGGAGCCCACCTGAACCTTT |
| Mouse miGPX4-R-1 | CCTGAAAGGTTCAGGTGGGCTCCGTCAGTCAGTGGCCAAAACGGAGCCCATTCCTGAACCTTTC |
| Mouse miGPX4-F-2 | TGCTGTTTCCTAGGACTTTGGCGTCCGTTTTGGCCACTGACTGACGGACGCCAGTCCTAGGAAA |
| Mouse miGPX4-R-2 | CCTGTTTCCTAGGACTGGCGTCCGTCAGTCAGTGGCCAAAACGGACGCCAAAGTCCTAGGAAAC |
| Human miNRF2-F-1 | TGCTGAGTAGTTGGCAGATCCACTGGGTTTTGGCCACTGACTGACCCAGTGGATGCCAACTACT |
| Human miNRF2-R-1 | CCTGAGTAGTTGGCATCCACTGGGTCAGTCAGTGGCCAAAACCCAGTGGATCTGCCAACTACTC |
| Human miNRF2-F-2 | TGCTGTAAAGTAGCAGGTGAGGGCATGTTTTGGCCACTGACTGACATGCCCTCCTGCTACTTTA |
| Human miNRF2-R-2 | CCTGTAAAGTAGCAGGAGGGCATGTCAGTCAGTGGCCAAAACATGCCCTCACCTGCTACTTTA |
| Mouse miNRF2-F-1 | TGCTGAATGTGGGCAACCTGGGAGTAGTTTTGGCCACTGACTGACTACTCCCATTGCCCACATT |
| Mouse miNRF2-R-1 | CCTGAATGTGGGCAATGGGAGTAGTCAGTCAGTGGCCAAAACTACTCCCAGGTTGCCCACATTC |
| Mouse miNRF2-F-2 | TGCTGAATAGCTCCTGCCAAACTTGCGTTTTGGCCACTGACTGACGCAAGTTTCAGGAGCTATT |
| Mouse miNRF2-R-2 | CCTGAATAGCTCCTGAAACTTGCGTCAGTCAGTGGCCAAAACGCAAGTTTGGCAGGAGCTATTC |
| Human miSLC7A11-F-1 | TGCTGATAACCTGGAGACAGCAAACAGTTTTGGCCACTGACTGACTGTTTGCTCTCCAGGTTAT |
| Human miSLC7A11-R-1 | CCTGATAACCTGGAGAGCAAACAGTCAGTCAGTGGCCAAAACTGTTTGCTGTCTCCAGGTTATC |
| Human miSLC7A11-F-2 | TGCTGAAATCAGCCCAGCAACTGCCAGTTTTGGCCACTGACTGACTGGCAGTTTGGGCTGATTT |
| Human miSLC7A11-R-2 | CCTGAAATCAGCCCAAACTGCCAGTCAGTCAGTGGCCAAAACTGGCAGTTGCTGGGCTGATTTC |
| Mouse miSLC7A11-F-1 | TGCTGATTACGAGCAGTTCCACCCAGGTTTTGGCCACTGACTGACCTGGGTGGCTGCTCGTAAT |
| Mouse miSLC7A11-R-1 | CCTGATTACGAGCAGCCACCCAGGTCAGTCAGTGGCCAAAACCTGGGTGGAACTGCTCGTAATC |
| Mouse miSLC7A11-F-2 | TGCTGTTTAGAAGACTATAGAGGTCTGTTTTGGCCACTGACTGACAGACCTCTAGTCTTCTAAA |

| | |
|---|---|
| Mouse miSLC7A11-R-2 | CCTGTTTAGAAGACTAGAGGTCTGTCAGTCAGTGGCCAAAACAGACCTCTATAGTCTTCTAAAC |
| Human miFPN-F | TGCTGTCTACCTGCAGCTTACATGATGTTTTGGCCACTGACTGACATCATGTACTGCAGGTAGA |
| Human miFPN-R | CCTGTCTACCTGCAGTACATGATGTCAGTCAGTGGCCAAAACATCATGTAAGCTGCAGGTAGAC |
| Mouse miFPN-F | TGCTGTATACAGACTCACTGATTTGCGTTTTGGCCACTGACTGACGCAAATCAGAGTCTGTATA |
| Mouse miFPN-R | CCTGTATACAGACTCTGATTTGCGTCAGTCAGTGGCCAAAACGCAAATCAGTGAGTCTGTATAC |
| Human miLCN2-F | TGCTGTAATGTTGCCCAGCGTGAACTGTTTTGGCCACTGACTGACAGTTCACGGGGCAACATTA |
| Human miLCN2-R | CCTGTAATGTTGCCCCGTGAACTGTCAGTCAGTGGCCAAAACAGTTCACGCTGGGCAACATTAC |
| Mouse miLCN2-F | TGCTGTCAAGTTCTGAGTTGAGTCCTGTTTTGGCCACTGACTGACAGGACTCATCAGAACTTGA |
| Mouse miLCN2-R | CCTGTCAAGTTCTGATGAGTCCTGTCAGTCAGTGGCCAAAACAGGACTCAACTCAGAACTTGAC |
| miNT-F | TGCTGAAATGTACTGCGCGTGGAGACGTTTTGGCCACTGACTGACGTCTCCACGCAGTACATTT |
| miNT-R | CCTGAAATGTACTGCGTGGAGACGTCAGTCAGTGGCCAAAACGTCTCCACGCGCAGTACATTTC |

**Table S3. Primer sequences used for qPCR.**

| Name | Primer sequence (5′-3′) | Name | Primer sequence (5′-3′) |
|---|---|---|---|
| AAV-F | TGCATGACCAGGCTCAGCTA | AAV-R | GACAGGGAAGGGAGCAGTG |
| Mouse RELA-F | TGCGATTCCGCTATAAATGCG | Mouse RELA-R | ACAAGTTCATGTGGATGAGGC |
| Human GAPDH-F | ATTTGGTCGTATTGGGCG | Human GAPDH-R | CTCGCTCCTGGAAGATGG |
| Mouse GAPDH-F | TCACCACCATGGAGAAGGC | Mouse GAPDH-R | GCTAAGCAGTTGGTGGTGCA |
| Human FSP1-F | GTGAGCGGGTGAGCAATCT | Human FSP1-R | CTTGATGCCGGTGCAGAGAA |
| Human FTH1-F | CCCCCATTTGTGTGACTTCAT | Human FTH1-R | GCCCGAGGCTTAGCTTTCATT |
| Human GPX4-F | GAGGCAAGACCGAAGTAAACTAC | Human GPX4-R | CCGAACTGGTTACACGGGAA |
| Human NRF2-F | TCAGCGACGGAAAGAGTATGA | Human NRF2-R | CCACTGGTTTCTGACTGGATGT |
| Human SLC7A11-F | GCGTGGGCATGTCTCTGAC | Human SLC7A11-R | GCTGGTAATGGACCAAAGACTTC |
| Human FPN-F | CACAACCGCCAGAGAGGATG | Human FPN-R | CACATCCGATCTCCCCAAGT |
| Human LCN2-F | CCCGCAAAAGATGTATGCCA | Human LCN2-R | CTCACCACTCGGACGAGGTA |
| Mouse FSP1-F | CTGCCTACCGCAGTGCATT | Mouse FSP1-R | ACGCCATCATTTCTGCCCA |
| Mouse FTH1-F | CAAGTGCGCCAGAACTACCA | Mouse FTH1-R | GCCACATCATCTCGGTCAAAA |
| Mouse GPX4-F | GATGGAGCCCATTCCTGAACC | Mouse GPX4-R | CCCTGTACTTATCCAGGCAGA |
| Mouse NRF2-F | TCTTGGAGTAAGTCGAGAAGTGT | Mouse NRF2-R | GTTGAAACTGAGCGAAAAAGGC |
| Mouse SLC7A11-F | GGCACCGTCATCGGATCAG | Mouse SLC7A11-R | CTCCACAGGCAGACCAGAAAA |
| Mouse FPN-F | TGGAACTCTATGGAAACAGCCT | Mouse FPN-R | TGGCATTCTTATCCACCCAGT |
| Mouse LCN2-F | TGGCCCTGAGTGTCATGTG | Mouse LCN2-R | CTCTTGTAGCTCATAGATGGTGC |
| Mouse Ki67-F | ATCATTGACCGCTCCTTTAGGT | Mouse Ki67-R | GCTCGCCTTGATGGTTCCT |
| Mouse TIM-3-F | ACTCTACCTACATCTGGGACACT | Mouse TIM-3-R | TCTCCTTTGTTGAGATCGCCC |
| Mouse CD34-F | TTCCTGATGAACCGTCGCAG | Mouse CD34-R | TGGTAAGCAGGGTTGTGAGG |
| Mouse CD38-F | TTTAGCCAGGTGTCTGGGGA | Mouse CD38-R | AAGTGCTTCGTGGTAGGCTC |
| Mouse CD133-F | CCTTGTGGTTCTTACGTTTGTTG | Mouse CD133-R | CGTTGACGACATTCTCAAGCTG |
| Mouse CD44-F | TCGATTTGAATGTAACCTGCCG | Mouse CD44-R | CAGTCCGGGAGATACTGTAGC |
| Mouse ALDH1-F | GGAATACCGTGGTTGTCAAGCC | Mouse ALDH1-R | CCAGGGACAATGTTTACCACGC |
| Mouse Tyrp1-F | ACTTGATGGGATCCAGAAGC | Mouse Tyrp1-R | CTGATTGGTCCACCCTCAGT |

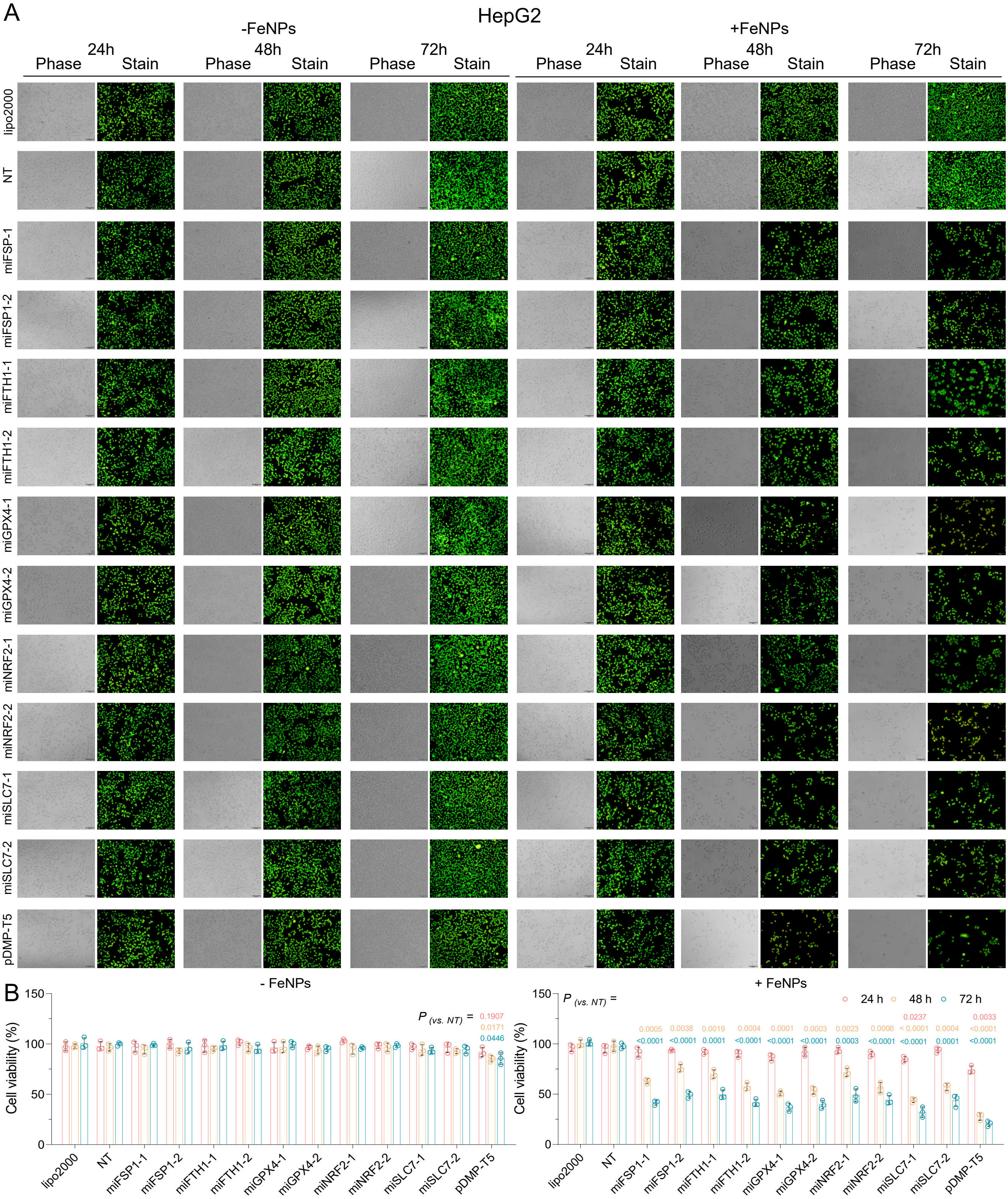


**Figure S1. Treatment of HepG2 with pDMP-miR vectors and FeNPs.** Cells were transfected by various plasmids overnight, then incubated with or without 50 µg/mL FeNPs for 24, 48, 72 h. (A) Representative images of AO&EB-stained cells. (B) The cell viability detected by the CCK-8 assay. All values are mean ± s.d. (n = 3 wells). Red, orange, and blue respectively represents the statistical significance obtained by comparing the data of all other groups with pDMP-NT at 24 h, 48 h, and 72 h. miFSP1, pDMP-miFSP1; miFTH1, pDMP-miFTH1; miNRF2, pDMP-miNRF2; miGPX4, pDMP-miGPX4; miSLC7, pDMP-miSLC7A11. -1, -miR1; -2, miR2 (two miRs were designed for each target gene).

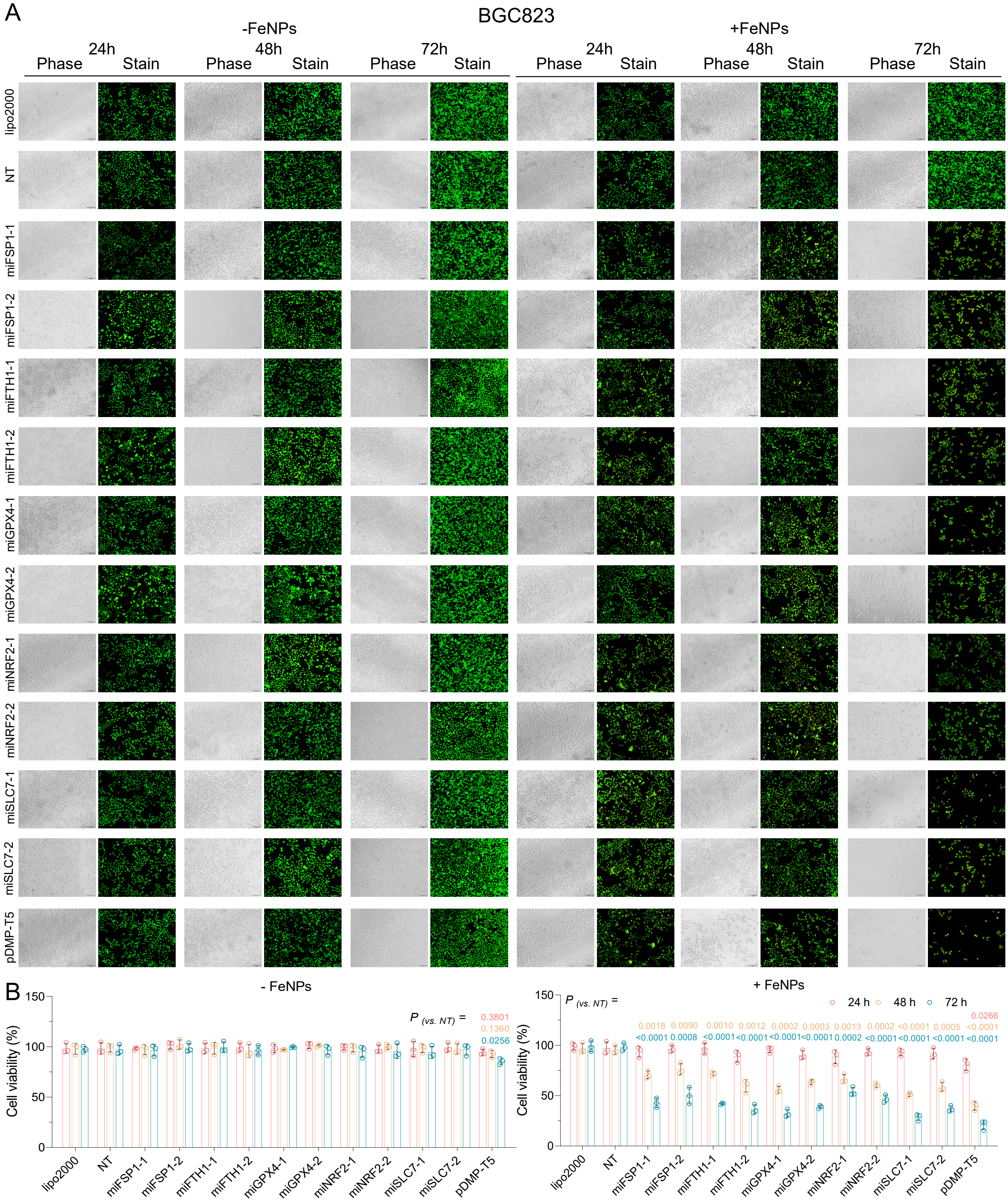


**Figure S2.** Treatment of BGC823 with pDMP-miR vectors and FeNPs. Cells were transfected by various plasmids overnight, then incubated with or without 50 μg/mL FeNPs for 24, 48, 72 h. (A) Representative images of AO&EB-stained cells. (B) The cell viability detected by the CCK-8 assay. All values are mean ± s.d. (n = 3 wells). Red, orange, and blue respectively represents the statistical significance obtained by comparing the data of all other groups with pDMP-NT at 24 h, 48 h, and 72 h. miFSP1, pDMP-miFSP1; miFTH1, pDMP-miFTH1; miNRF2, pDMP-miNRF2; miGPX4, pDMP-miGPX4; miSLC7, pDMP-miSLC7A11. -1, -miR1; -2, miR2 (two miRs were designed for each target gene).

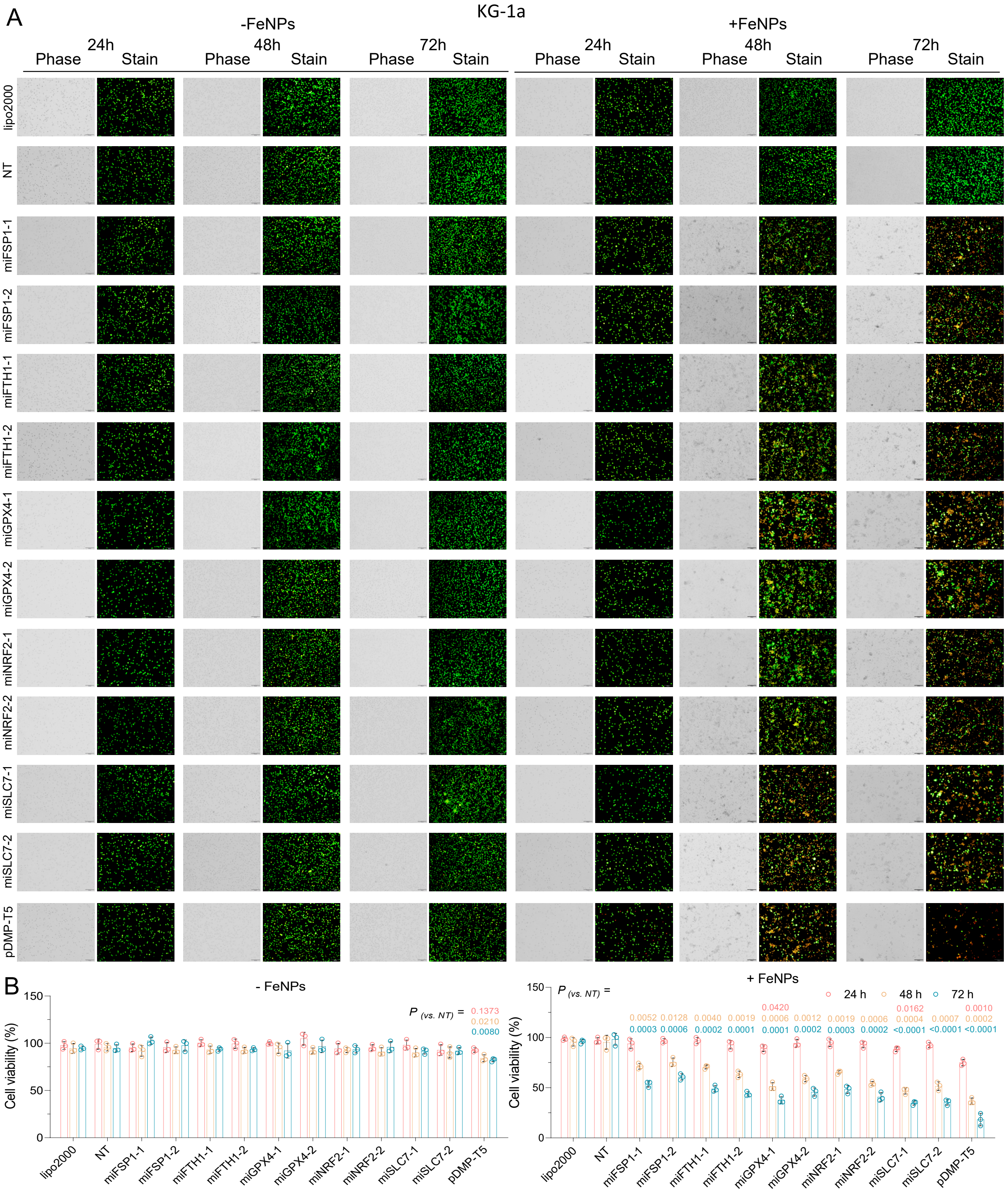


**Figure S3.** Treatment of KG-1a with pDMP-miR vectors and FeNPs. Cells were transfected by various plasmids overnight, then incubated with or without 50 μg/mL FeNPs for 24, 48, 72 h. (A) Representative images of AO&EB-stained cells. (B) The cell viability detected by the CCK8 assay. All values are mean ± s.d. (n = 3 wells). Red, orange, and blue respectively represents the statistical significance obtained by comparing the data of all other groups with pDMP-NT at 24 h, 48 h, and 72 h. miFSP1, pDMP-miFSP1; miFTH1, pDMP-miFTH1; miNRF2, pDMP-miNRF2; miGPX4, pDMP-miGPX4; miSLC7, pDMP-miSLC7A11. -1, -miR1; -2, miR2 (two miRs were designed for each target gene).

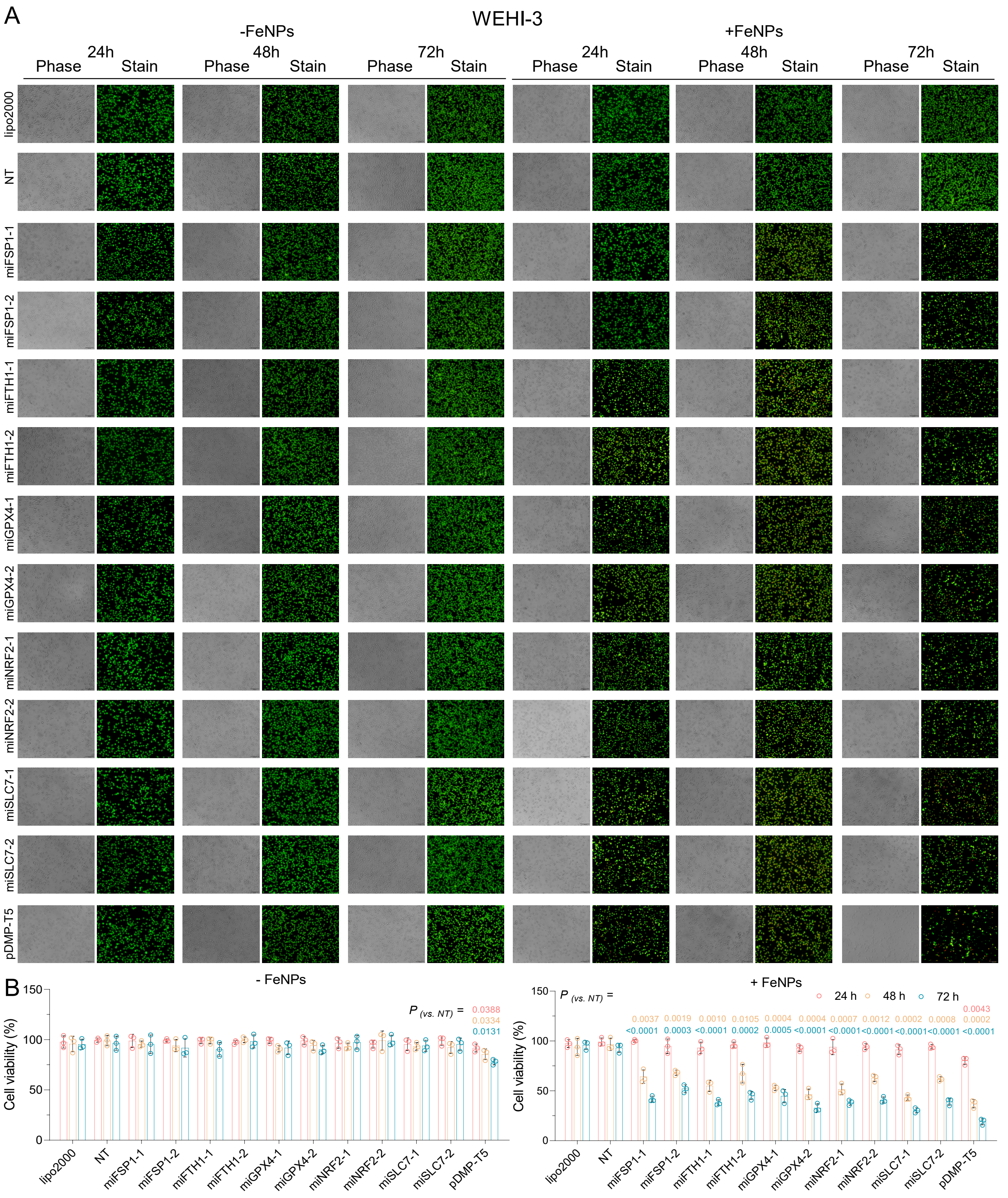


**Figure S4.** Treatment of WEHI-3 with pDMP-miR vectors and FeNPs. Cells were transfected by various plasmids overnight, then incubated with or without 50 μg/mL FeNPs for 24, 48, 72 h. (A) Representative images of AO&EB-stained cells. (B) The cell viability detected by the CCK-8 assay. All values are mean ± s.d. (n = 3 wells). Red, orange, and blue respectively represents the statistical significance obtained by comparing the data of all other groups with pDMP-NT at 24 h, 48 h, and 72 h. miFSP1, pDMP-miFSP1; miFTH1, pDMP-miFTH1; miNRF2, pDMP-miNRF2; miGPX4, pDMP-miGPX4; miSLC7, pDMP-miSLC7A11. -1, -miR1; -2, miR2 (two miRs were designed for each target gene).

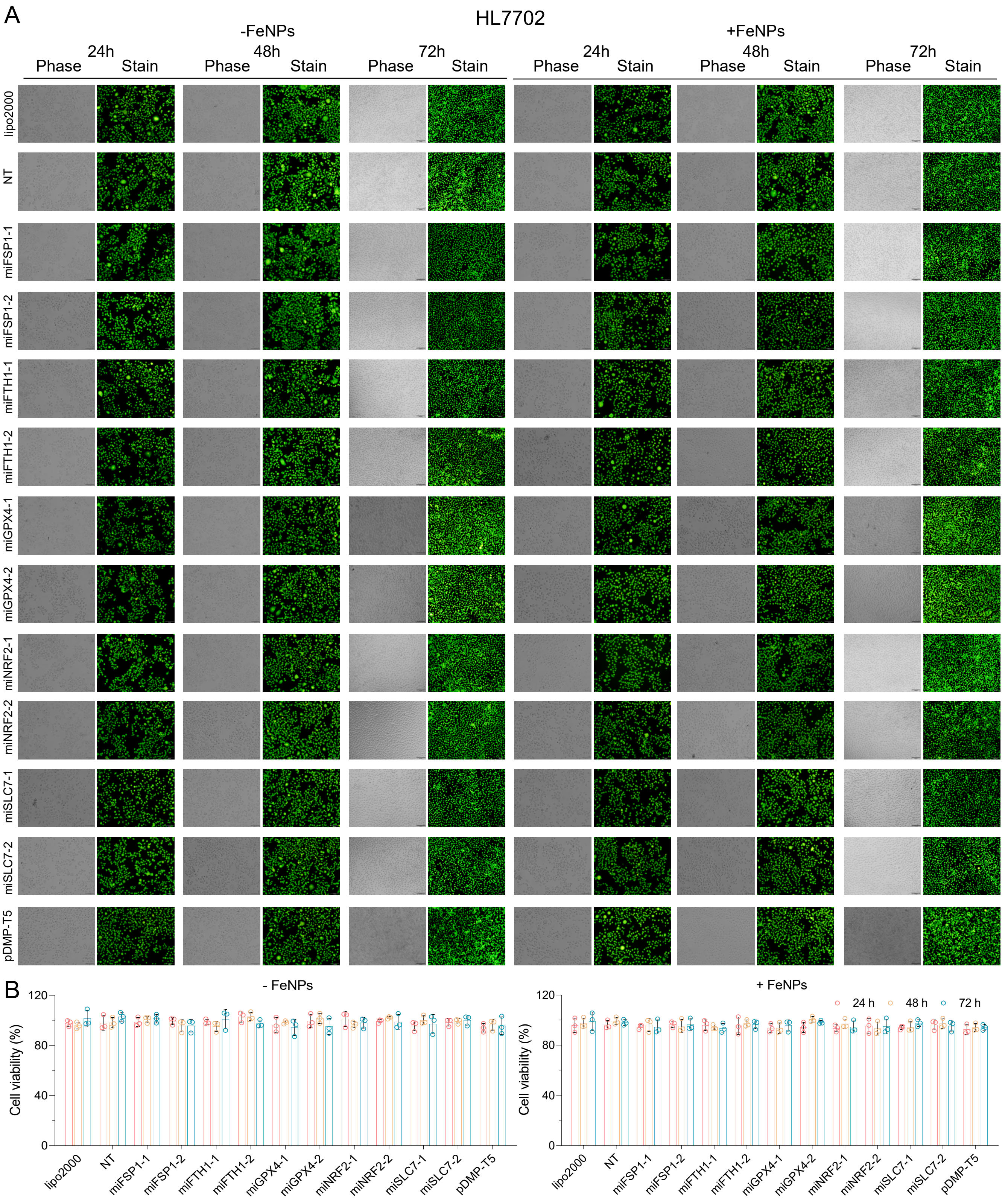


**Figure S5.** Treatment of HL7702 with pDMP-miR vectors and FeNPs. Cells were transfected by various plasmids overnight, then incubated with or without 50 μg/mL FeNPs for 24, 48, 72 h. (A) Representative images of AO&EB-stained cells. (B) The cell viability detected by the CCK-8 assay. All values are mean ± s.d. (n = wells). Red, orange, and blue respectively represents the statistical significance obtained by comparing the data of all other groups with pDMP-NT at 24 h, 48 h, and 72 h. miFSP1, pDMP-miFSP1; miFTH1, pDMP-miFTH1; miNRF2, pDMP-miNRF2; miGPX4, pDMP-miGPX4; miSLC7, pDMP-miSLC7A11. -1, -miR1; -2, miR2 (two miRs were designed for each target gene).

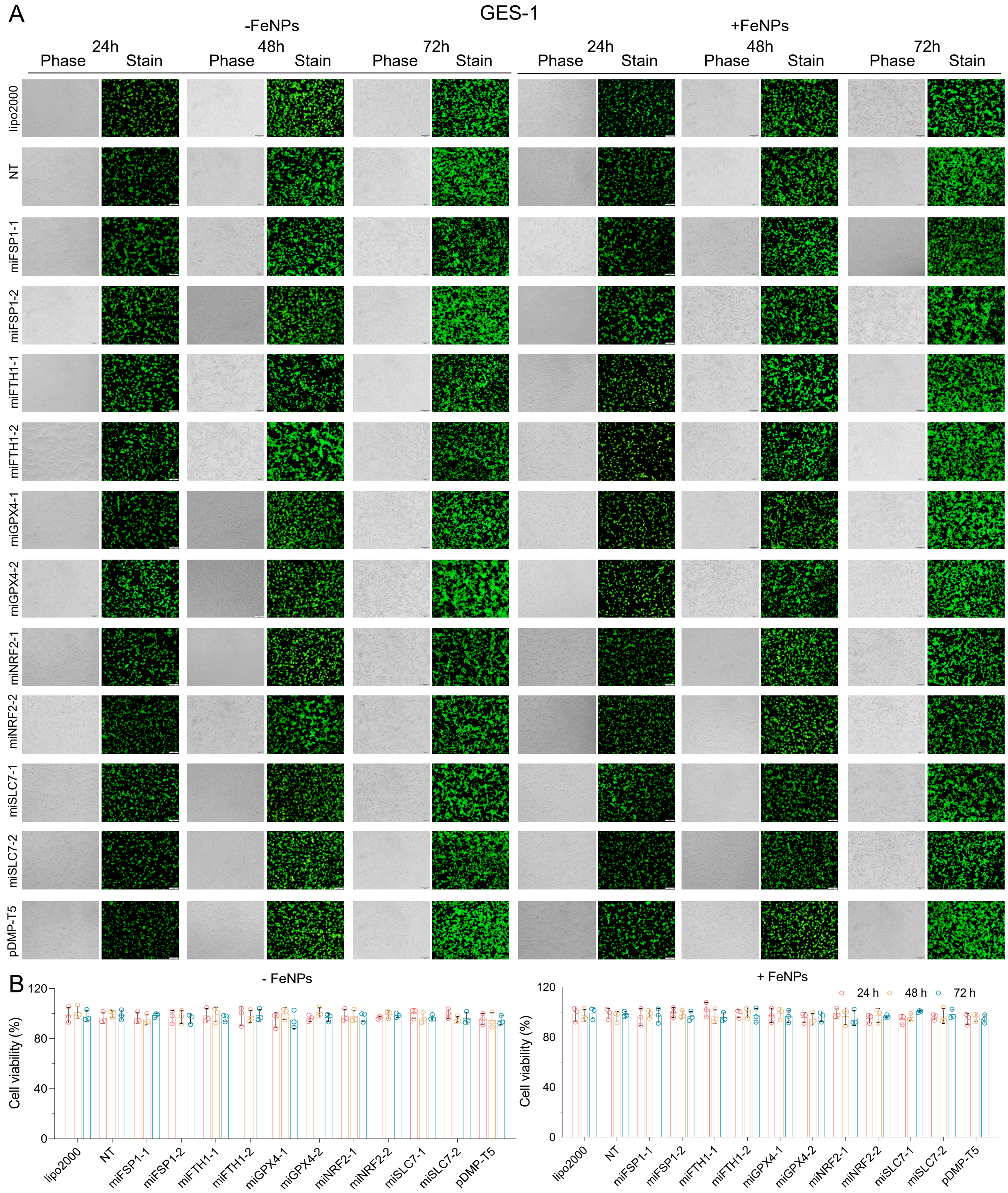


**Figure S6.** Treatment of GES-1 with pDMP-miR vectors and FeNPs. Cells were transfected by various plasmids overnight, then incubated with or without 50 μg/mL FeNPs for 24, 48, 72 h. (A) Representative images of AO&EB-stained cells. (B) The cell viability detected by the CCK-8 assay. All values are mean ± s.d. (n = 3 wells). Red, orange, and blue respectively represents the statistical significance obtained by comparing the data of all other groups with pDMP-NT at 24 h, 48 h, and 72 h. miFSP1, pDMP-miFSP1; miFTH1, pDMP-miFTH1; miNRF2, pDMP-miNRF2; miGPX4, pDMP-miGPX4; miSLC7, pDMP-miSLC7A11. -1, -miR1; -2, miR2 (two miRs were designed for each target gene).

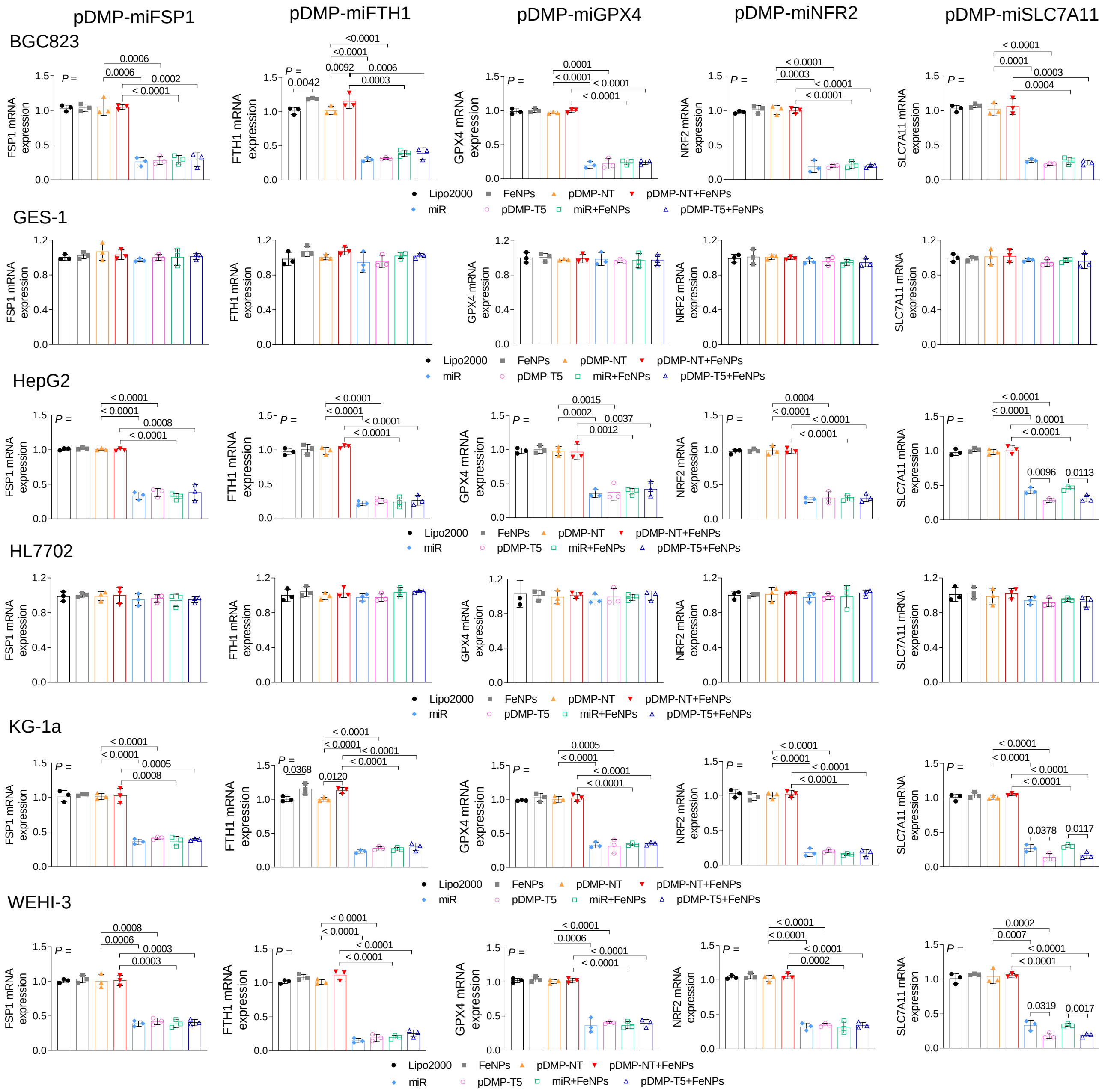

**Figure S7.** Gene expression in the pDMP-miR vector-treated cells. The mRNA levels of FSP1, FTH1, NRF2, GPX4, and SLC7A11 were analyzed by RT-qPCR in six cell lines (BGC823, GES-1, HepG2, HL7702, KG-1a, and WEHI-3). All values are mean ± s.d. (n = 3 wells). All figures use a same set of symbols. All significant difference is shown with P values. MiR, pDMP-miFSP1, pDMP-miFTH1, pDMP-miNRF2, pDMP-miGPX4, or pDMP-miSLC7A11.

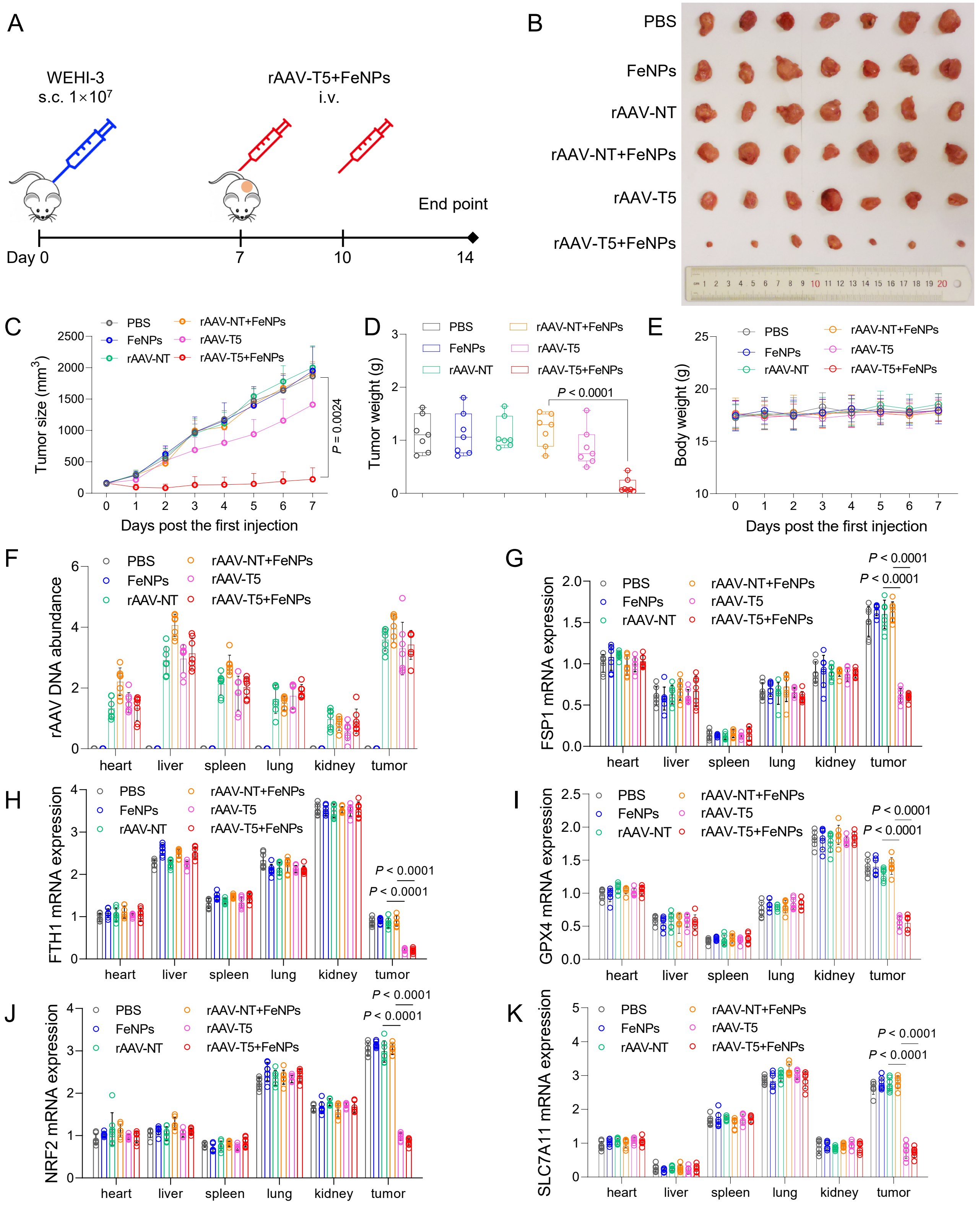


**Figure S8.** The in vivo antitumor effects of rAAV-T5 and FeNPs in the WEHI-3 xenograft mice. A–E, tumor growth detection. (A) Schematics of animal treatment. s.c., subcutaneously injection; i.v., intravenous injection. (B) Tumor imaging. (C) Tumor growth curve. (D) Tumor weight. (E) Average body weight. Data are presented as mean ± s.d. (n = 7 mice). (F) Abundance of virus DNA in tissues. (G–K) represented FSP1, FTH1, GPX4, NRF2, SLC7A11 mRNA expression in tissues, respectively (n = 7 mice).

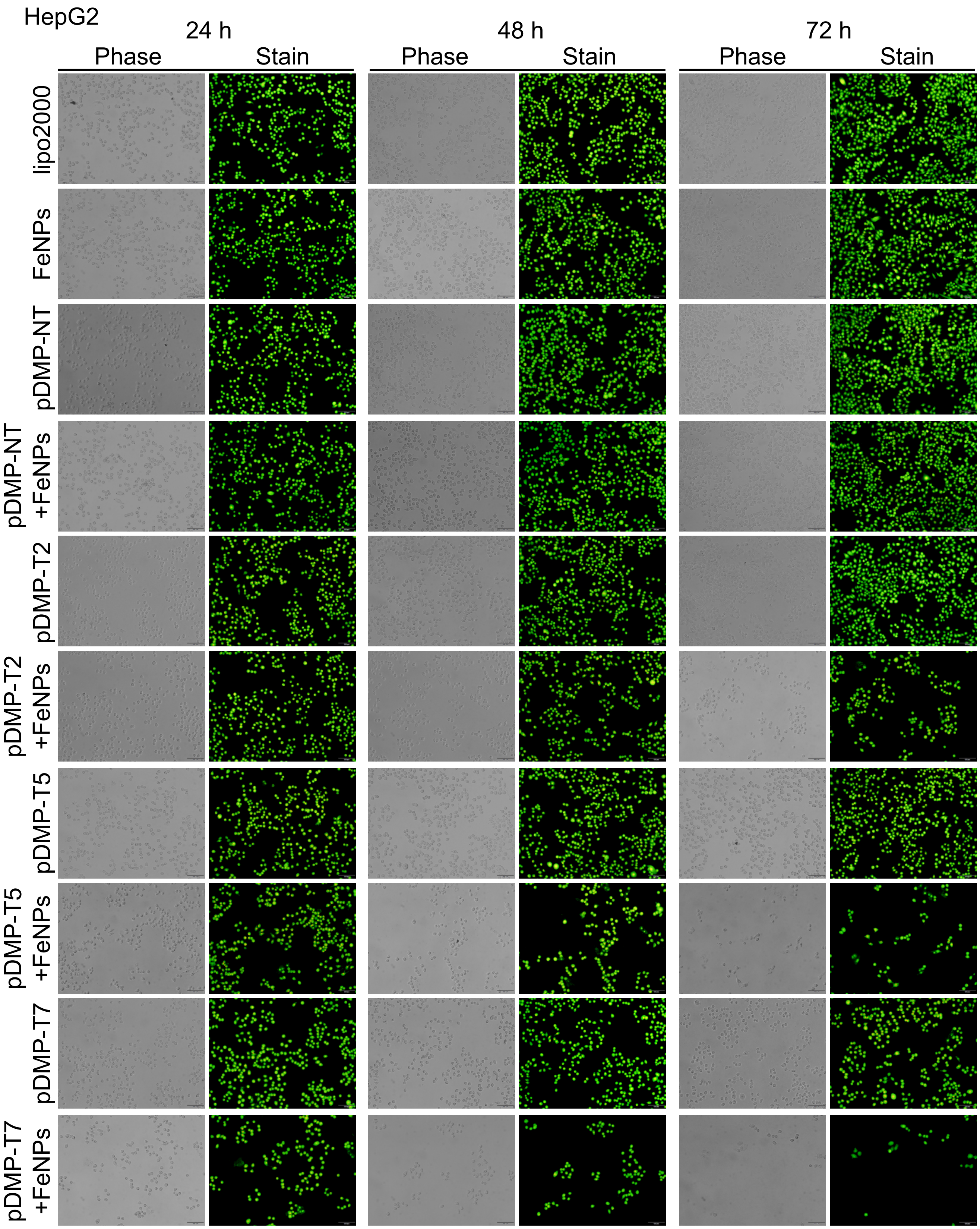


**Figure S9.** The effects of pDMP-T2/T5/T7 on the viability of HepG2. Cells were transfected with pDMP-T2/T5/T7 overnight. Cells were then cultured with or without 50 μg/mL FeNPs for 24 h, 48 h and 72 h, respectively. Cells were stained with AO&EB and imaged.

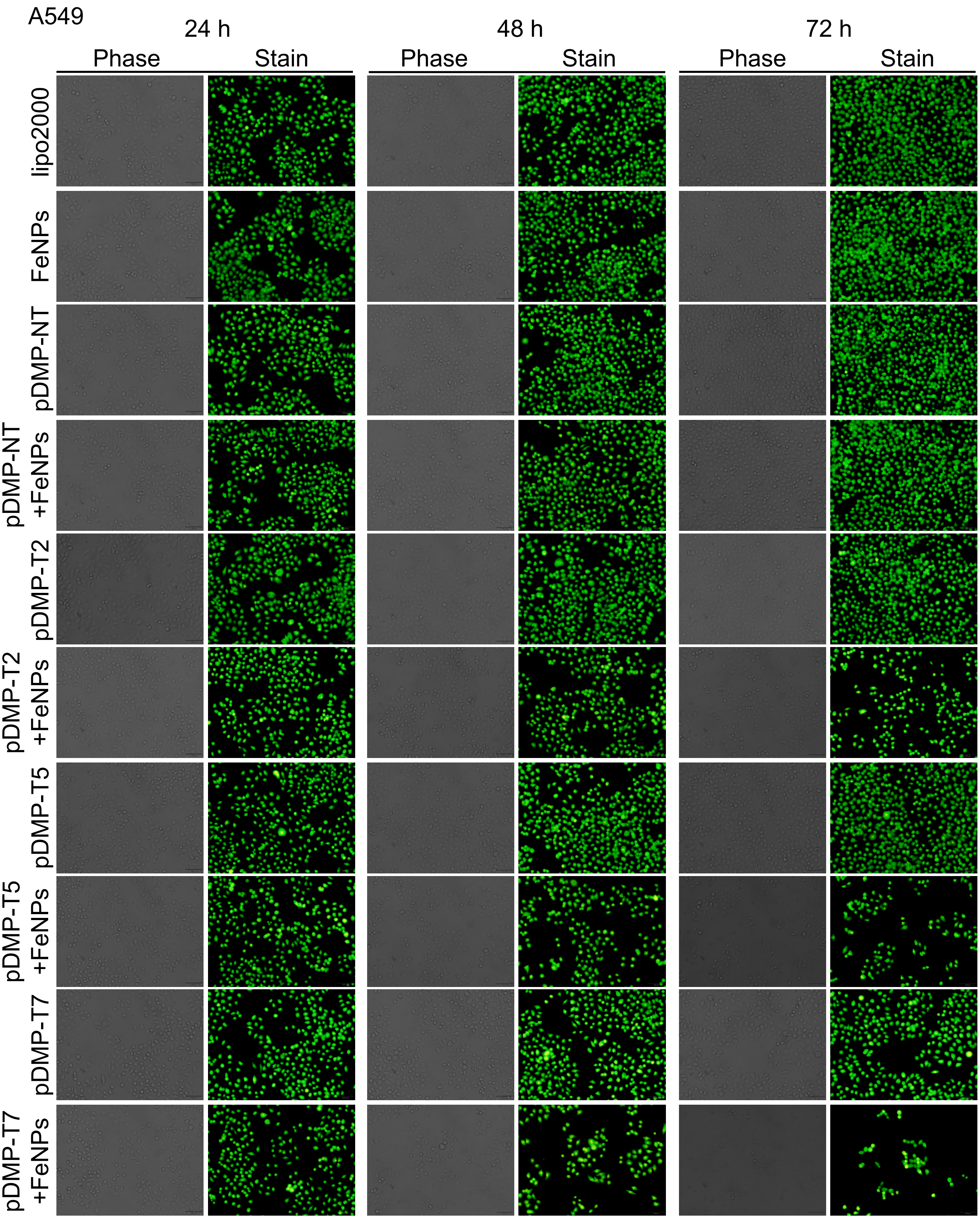


**Figure S10.** The effects of pDMP-T2/T5/T7 on the viability of A549. Cells were transfected with pDMP-T2/T5/T7 overnight. Cells were then cultured with or without 50 μg/mL FeNPs for 24 h, 48 h and 72 h, respectively. Cells were stained with AO&EB and imaged.

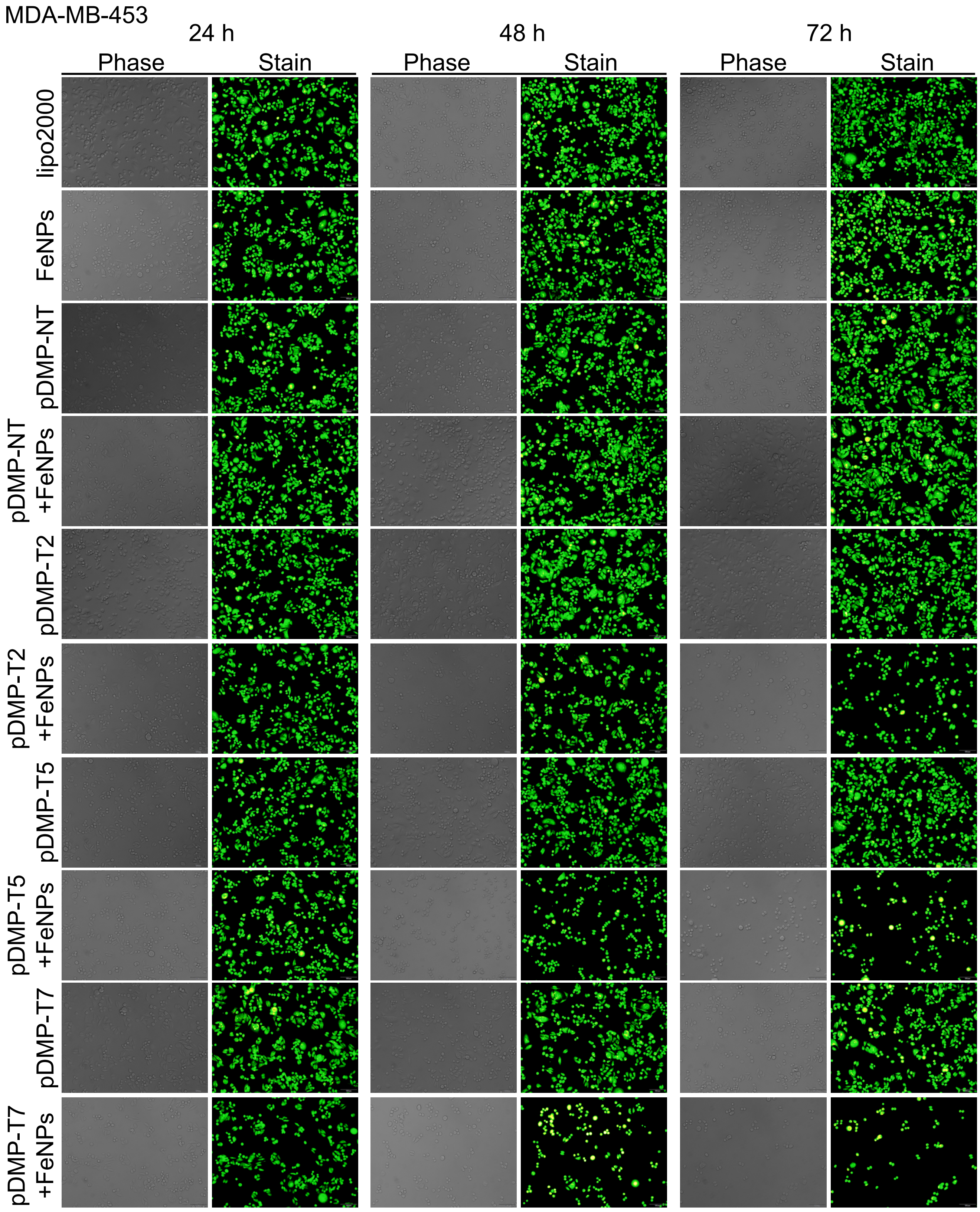


**Figure S11.** The effects of pDMP-T2/T5/T7 on the viability of MDA-MB-453. Cells were transfected with pDMP-T2/T5/T7 overnight. Cells were then cultured with or without 50 μg/mL FeNPs for 24 h, 48 h and 72 h, respectively. Cells were stained with AO&EB and imaged.

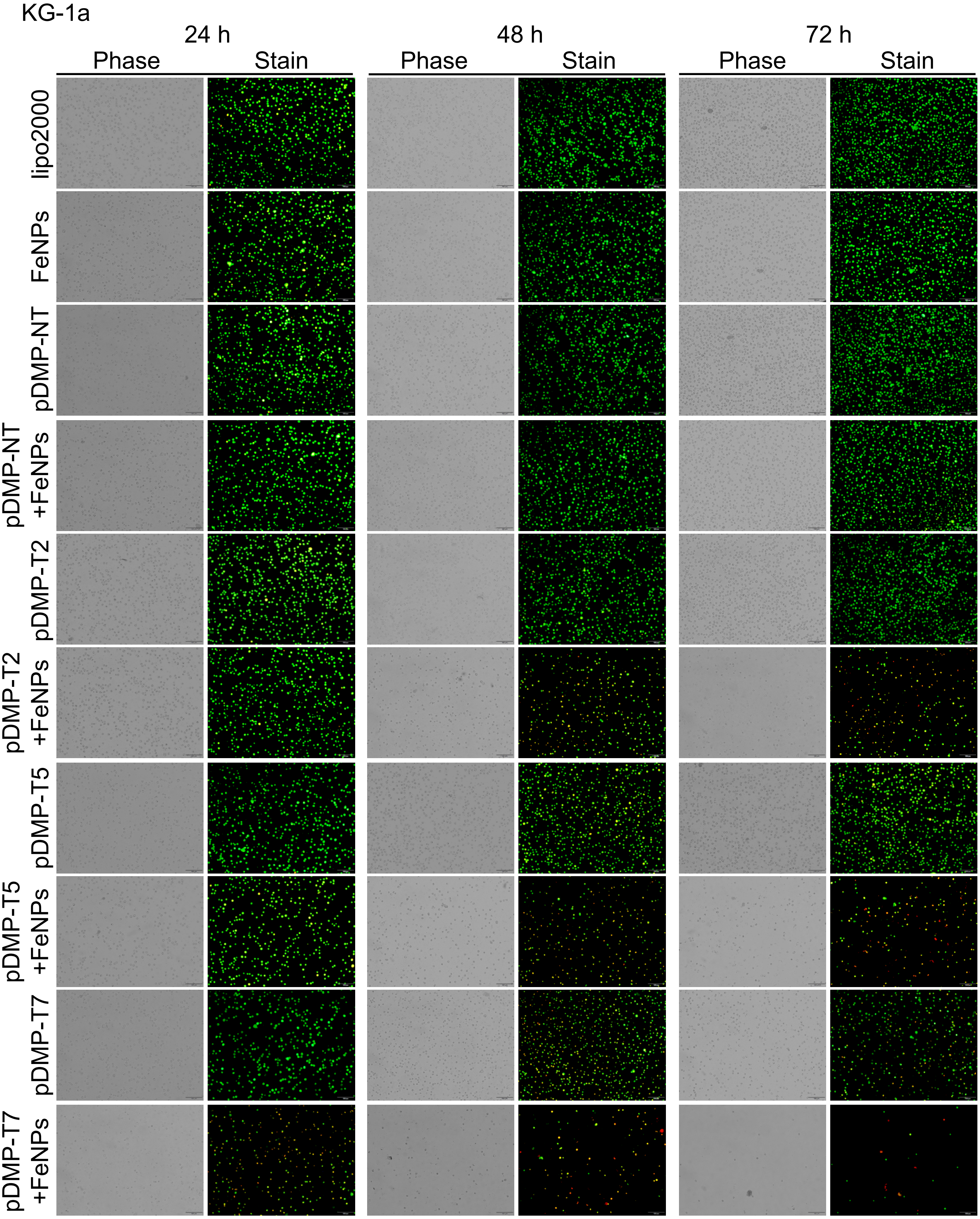


**Figure S12.** The effects of pDMP-T2/T5/T7 on the viability of KG-1a. Cells were transfected with pDMP-T2/T5/T7 overnight. Cells were then cultured with or without 50 μg/mL FeNPs for 24 h, 48 h and 72 h, respectively. Cells were stained with AO&EB and imaged.

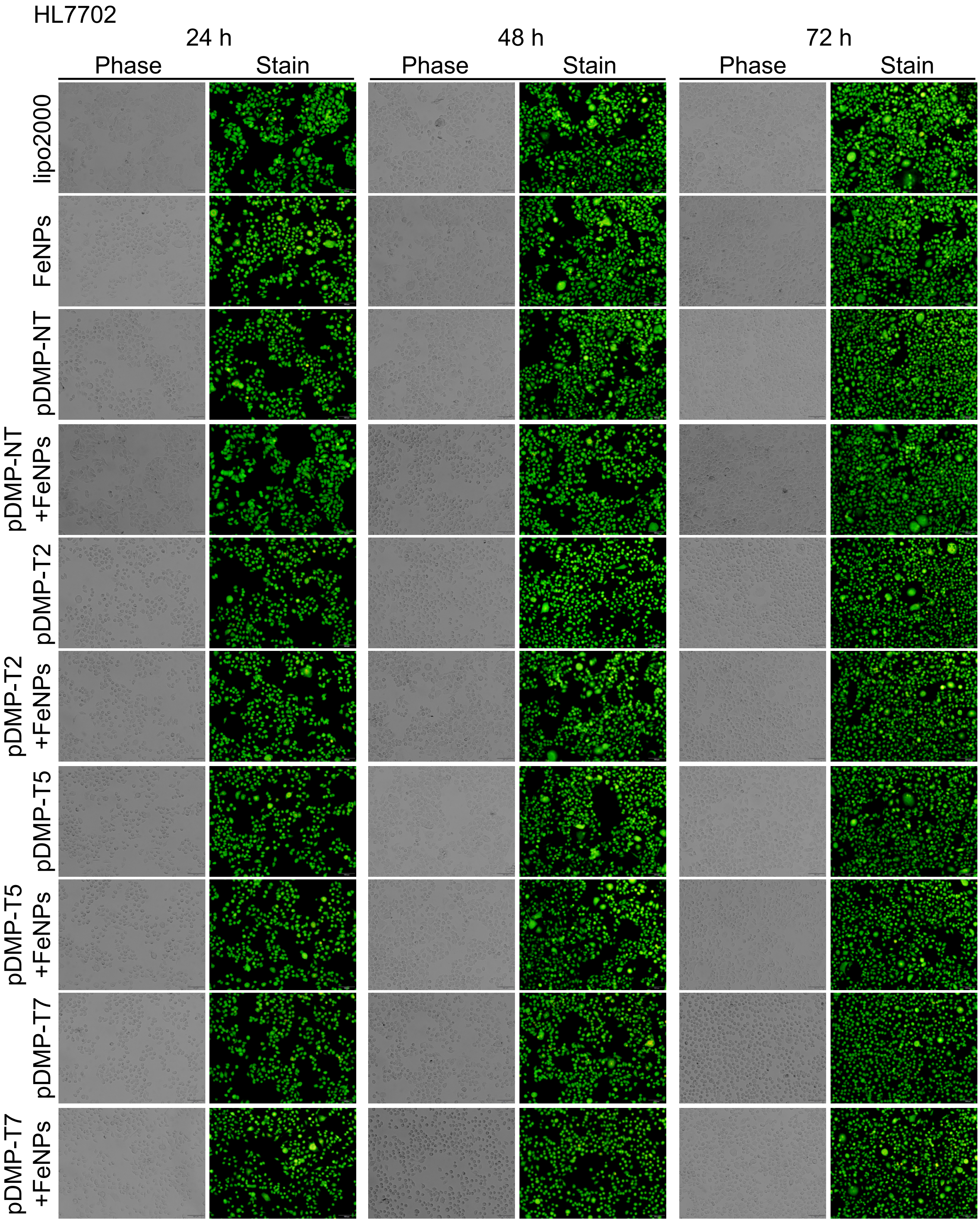


**Figure S13.** The effects of pDMP-T2/T5/T7 on the viability of HL7702. Cells were transfected with pDMP-T2/T5/T7 overnight. Cells were then cultured with or without 50 μg/mL FeNPs for 24 h, 48 h and 72 h, respectively. Cells were stained with AO&EB and imaged.

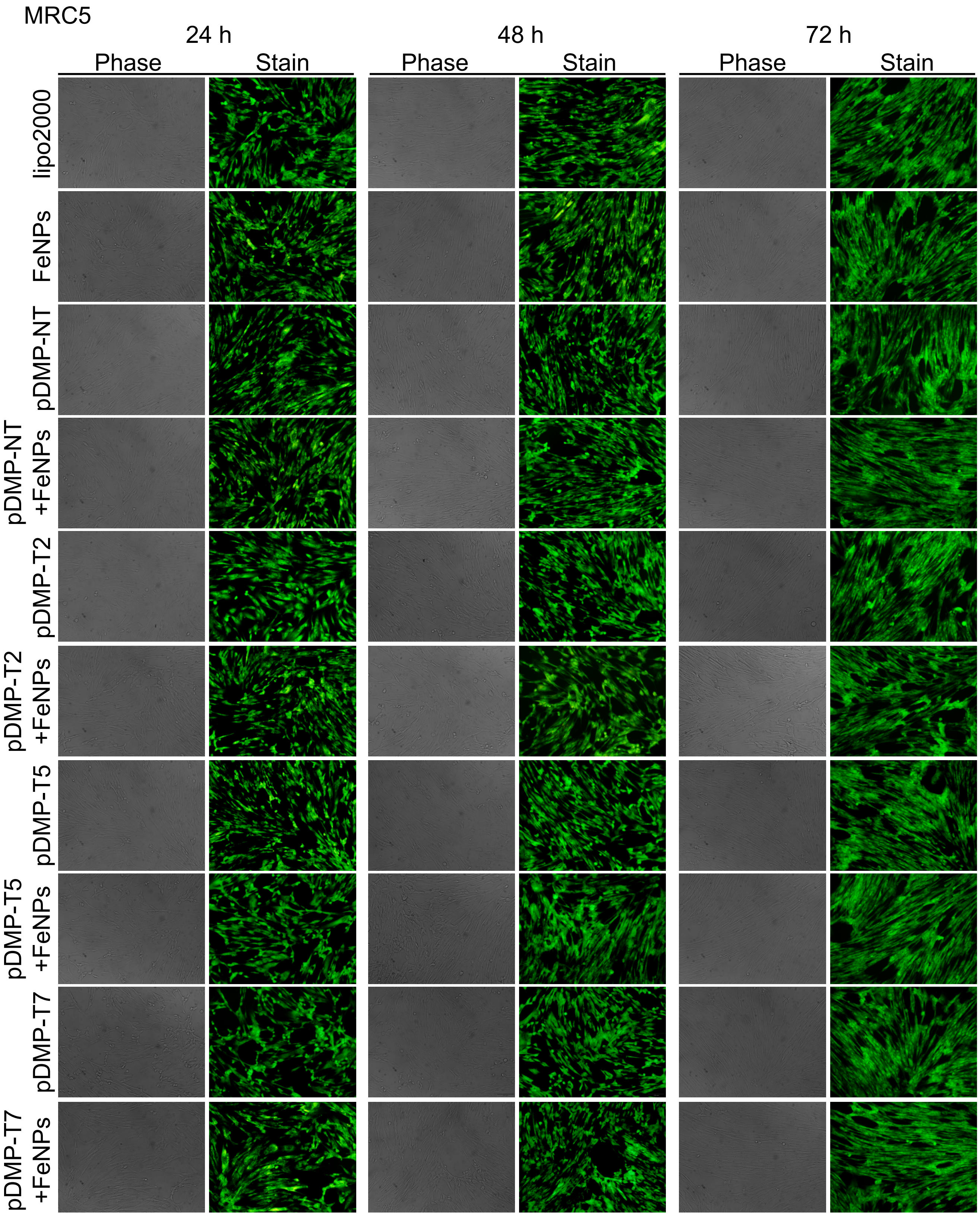


**Figure S14.** The effects of pDMP-T2/T5/T7 on the viability of MRC5. Cells were transfected with pDMP-T2/T5/T7 overnight. Cells were then cultured with or without 50 μg/mL FeNPs for 24 h, 48 h and 72 h, respectively. Cells were stained with AO&EB and imaged.

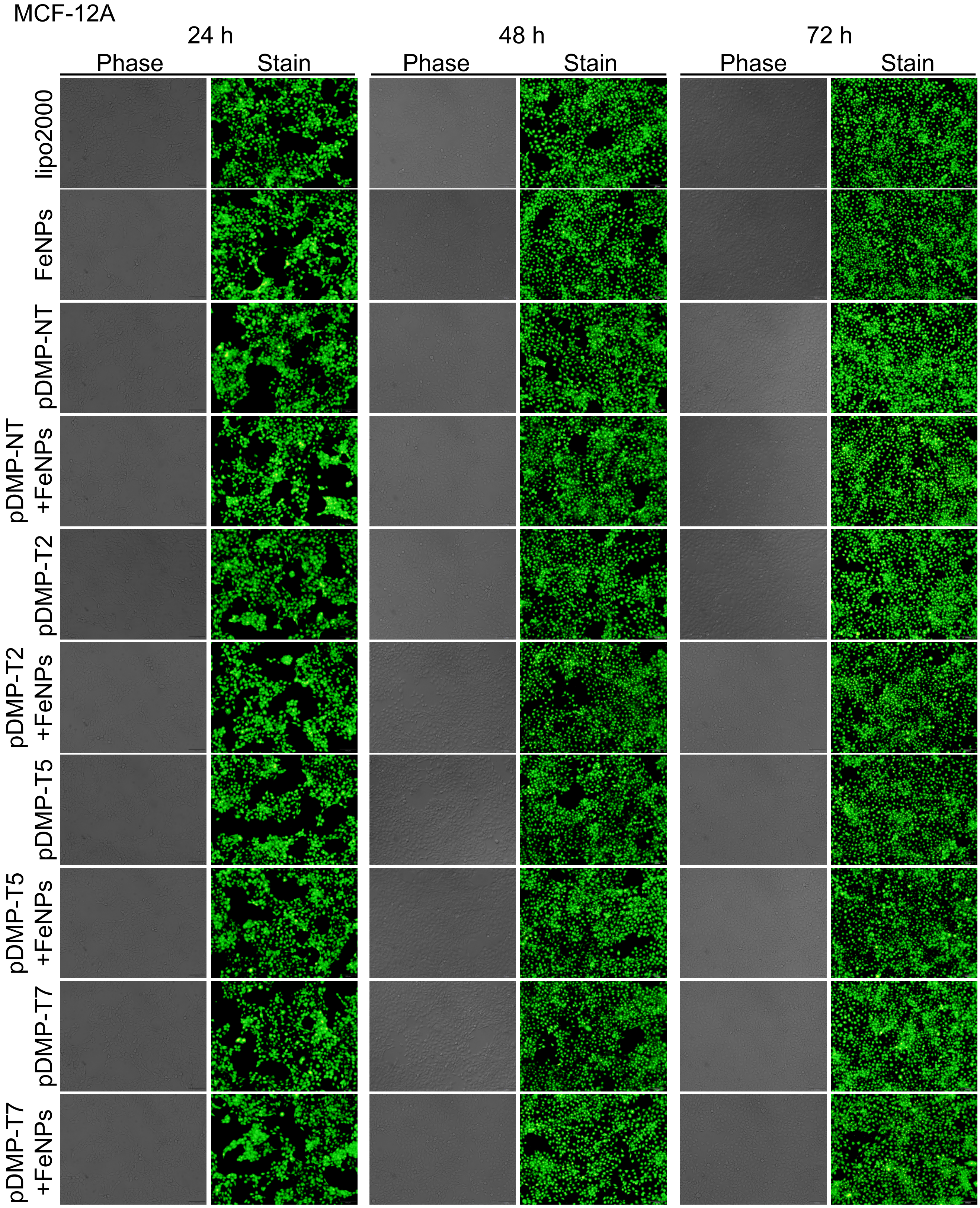


**Figure S15.** The effects of pDMP-T2/T5/T7 on the viability of MCF-12A. Cells were transfected with pDMP-T2/T5/T7 overnight. Cells were then cultured with or without 50 μg/mL FeNPs for 24 h, 48 h and 72 h, respectively. Cells were stained with AO&EB and imaged.

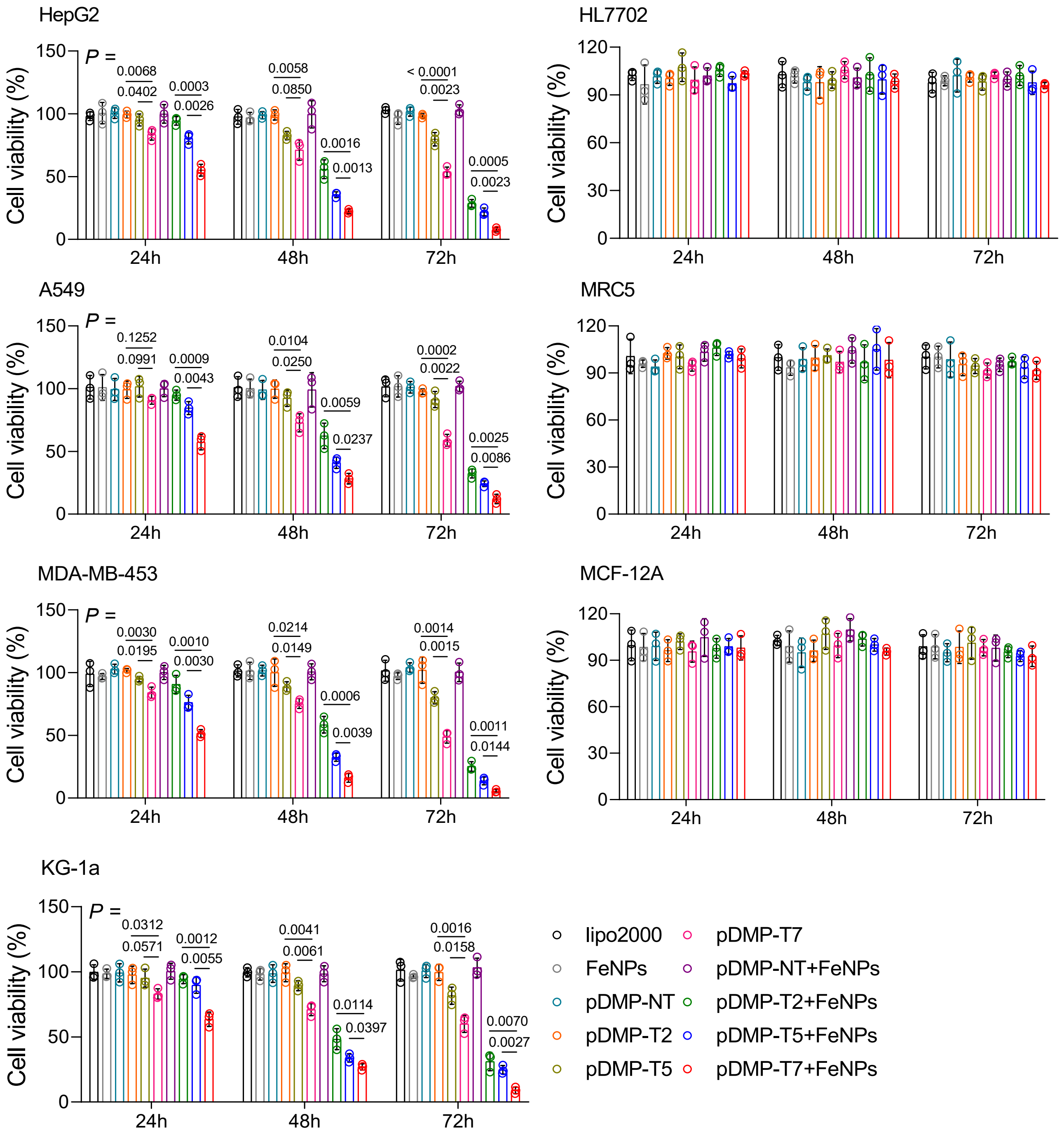


**Figure S16.** Cell viability of HepG2, HL7702, KG-1a, A549, MRC5, MDA-MB-453, and MCF-12A cells detected by the CCK-8 assay. Cells were transfected with pDMP-T2/T5/T7 overnight. Cells were then cultured with or without 50 μg/mL FeNPs for 24 h, 48 h and 72 h, respectively. All values are mean ± s.d. (n = 3 wells). All figures use a same set of symbols.

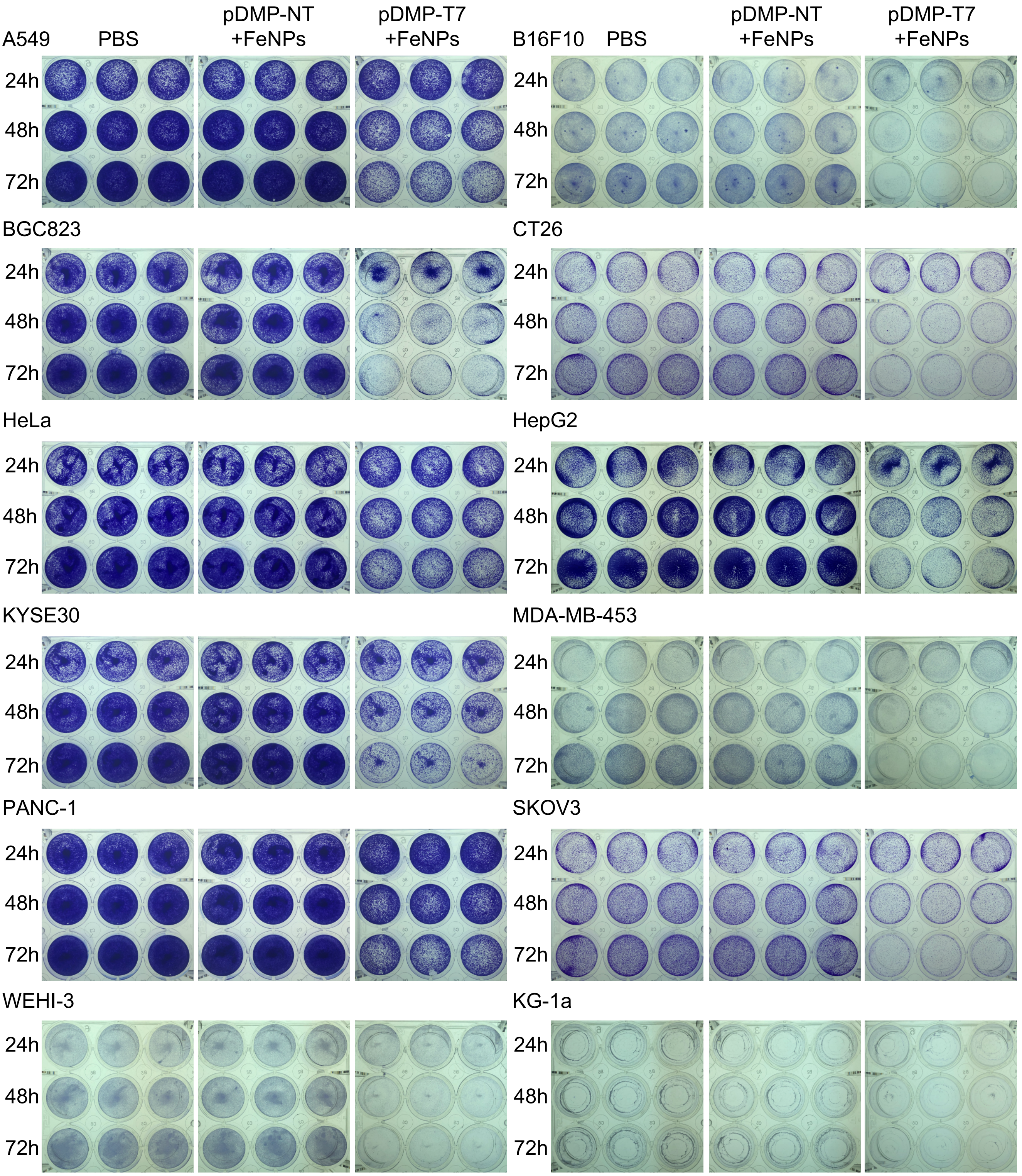


**Figure S17.** Effect of FAST treatment on viability of cancer cells detected by crystal violet assay. Cells were transfected with pDMP-T7/NT overnight. Cells were then cultured with 50 μg/mL FeNPs for 24 h, 48 h and 72 h, respectively. PBS, cells just transfected by Lipofectamine and treated with phosphate buffered saline (PBS). Cells were stained with crystal violet at the final concentration of 0.02% (w/v) for 5 min at room temperature. Each treatment was conducted in triplicates.

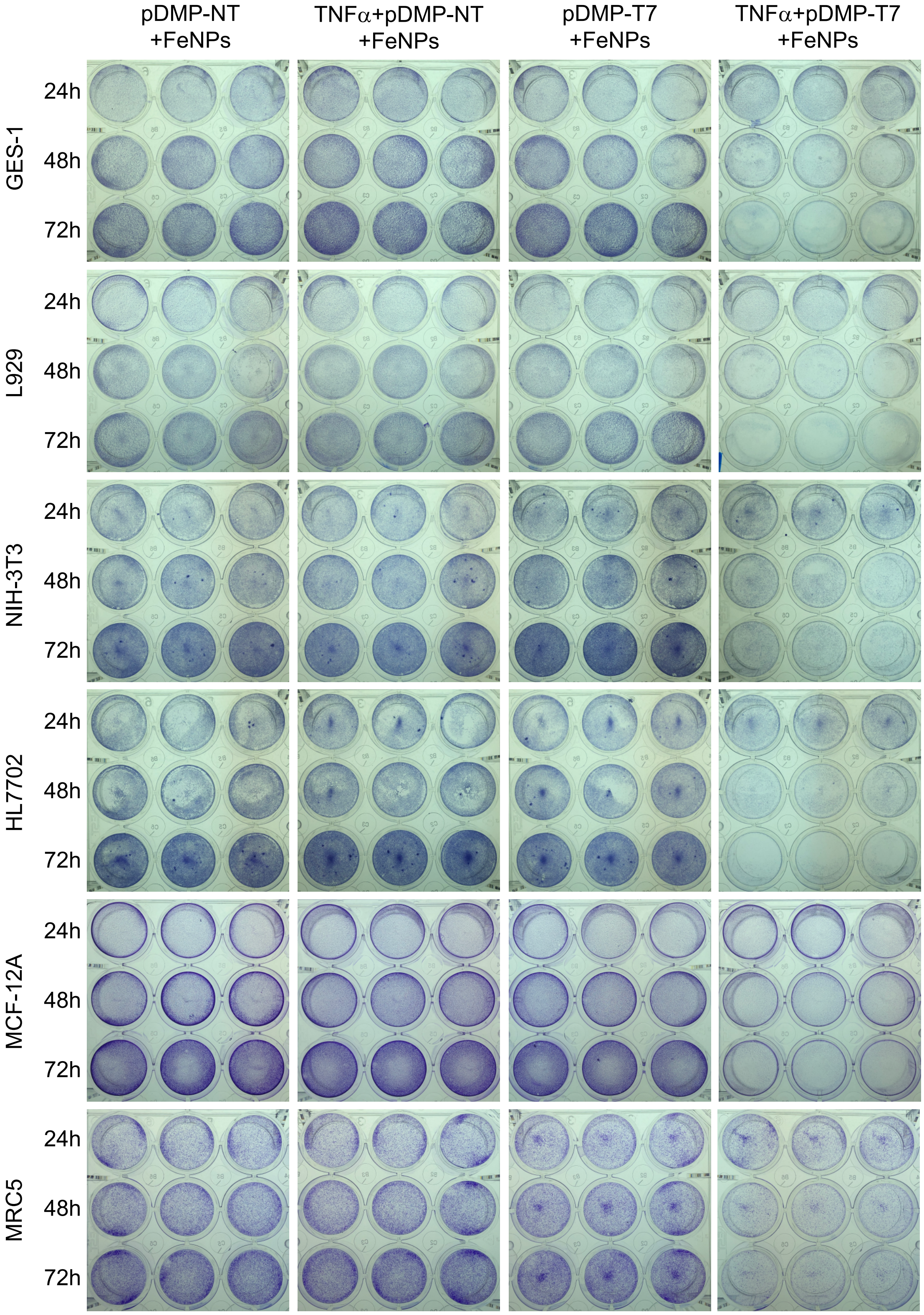


**Figure S18.** Effect of FAST treatment on viability of normal cells detected by crystal violet assay. Cells were transfected with pDMP-T7/NT overnight. Cells were then cultured with 50 μg/mL FeNPs for 24 h, 48 h and 72 h, respectively. If needed, cells were induced with TNFα at a final concentration of 10 ng/mL for 1 h before transfection. Cells were stained with crystal violet at the final concentration of 0.02% (w/v) for 5 min at room temperature. Each treatment was conducted in triplicates..

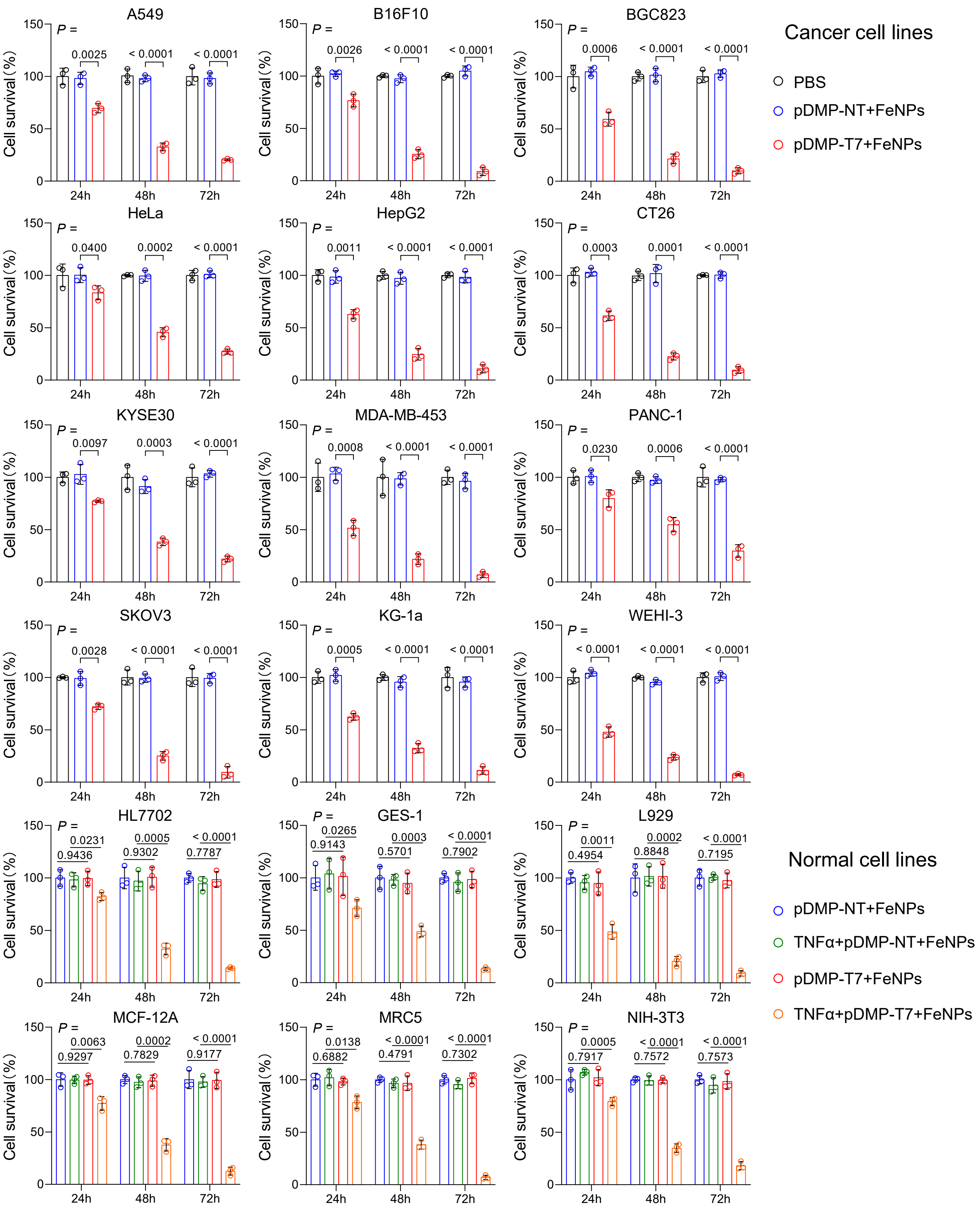


**Figure S19.** Effect of FAST treatment on viability of cancer and normal cells detected by crystal violet assay. The crystal violet-stained cells in Supplementary Figure 17 and 18 were eluted with 0.1 M sodium citrate in 50% (v/v) ethanol. The absorbance of elution at 585 nm was recorded. All values are mean ± s.d. (n = 3 wells).

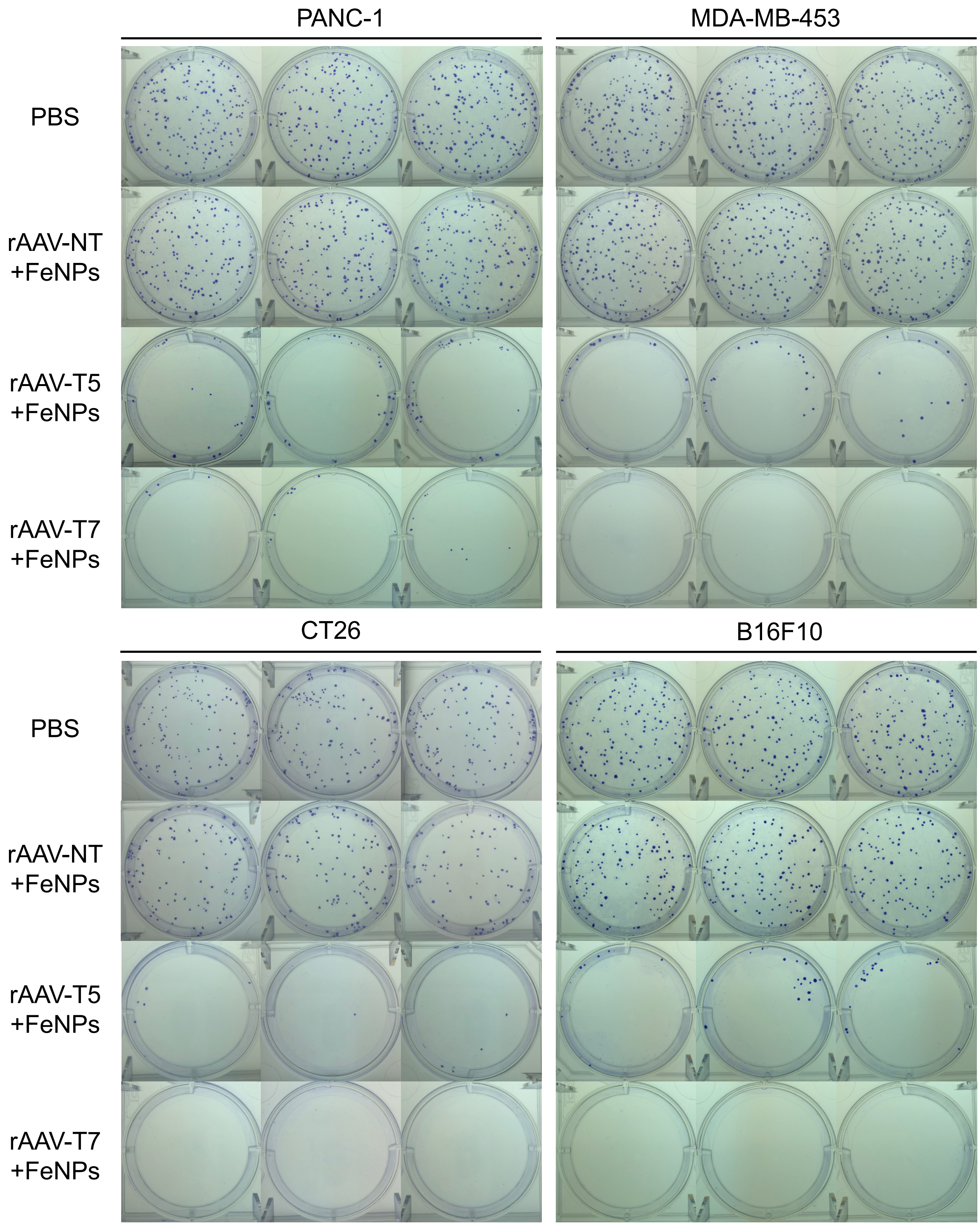


**Figure S20.** Effect of FAST treatment on clone formation of cancer cells. Cells were infected with rAAV-NT/T5/T7 at the dose of $1\times10^5$ vg per cell for 24 h and then incubated with 50 μg/mL FeNPs for another 48 h. PBS, cells just treated with phosphate buffered saline (PBS). Two hundred of treated cells were seeded into 6-well plate and cultured for 2 weeks. Cells were stained with crystal violet at the final concentration of 0.02% (w/v) for 5 min at room temperature. The stained cells were imaged. Each treatment was conducted in triplicates.

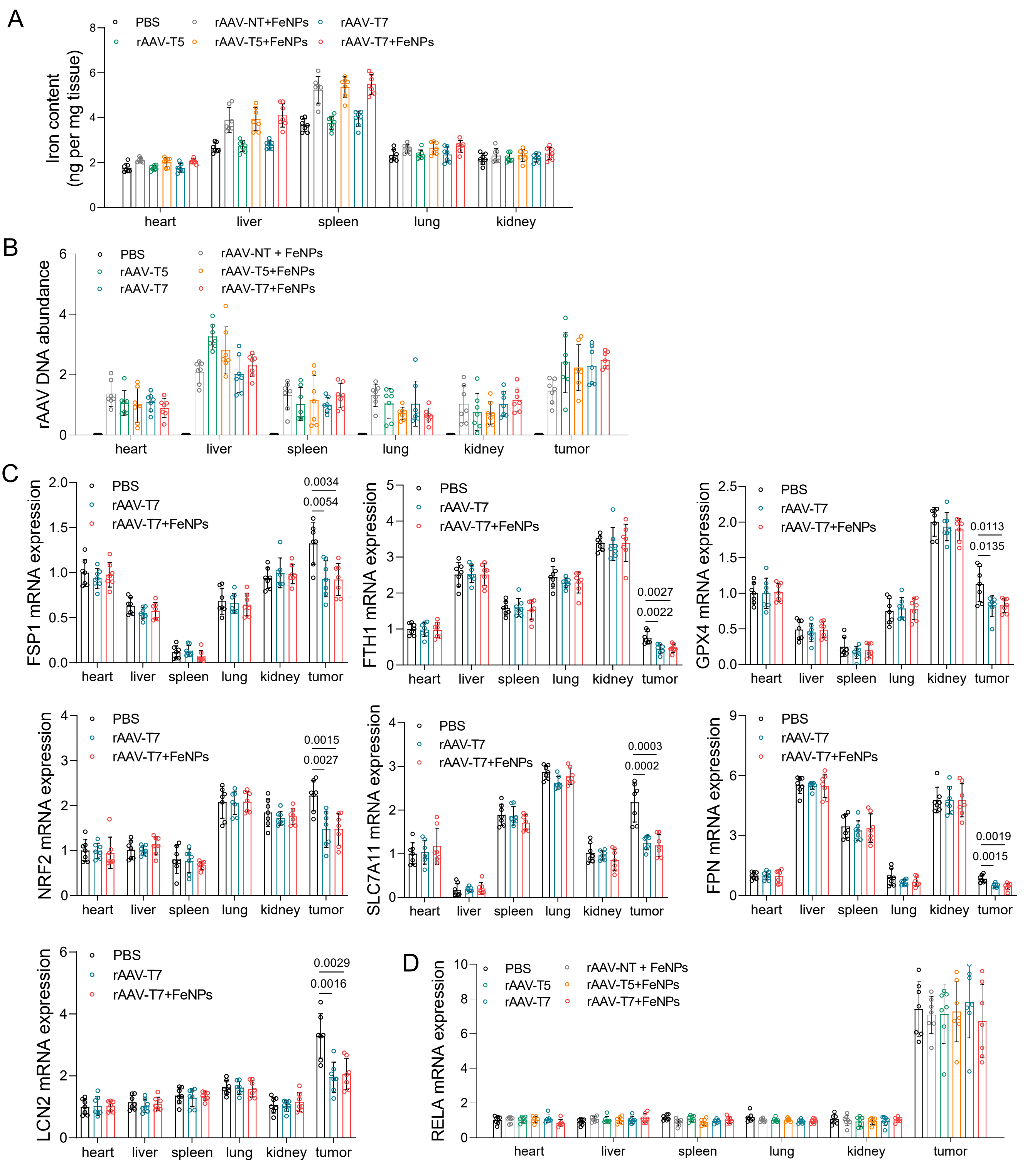


**Figure S21.** Effect of FAST treatment on the iron content and target gene expression in the WEHI-3 xenograft mice. (A) Iron content in tissues. (B) rAAV DNA abundance in tissues. (C) FSP1, FTH1, GPX4, NRF2, SLC7A11, FPN, LCN2 mRNA expression in tissues. (D) RELA mRNA expression in tissues. All data are presented as mean ± s.d. (n = 7 mice).

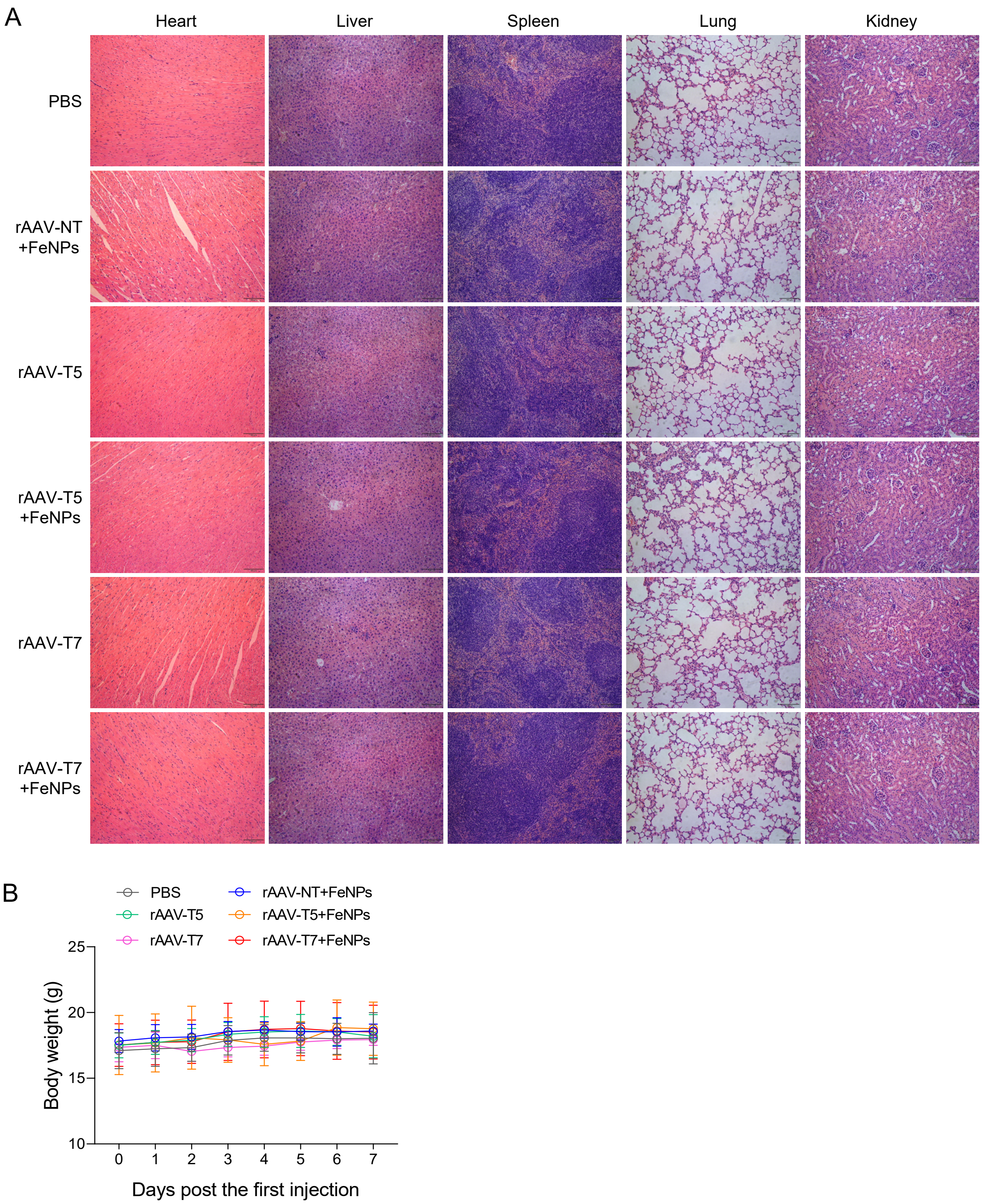


**Figure S22.** Effect of FAST treatment on the tissue structure and body weight of the WEHI-3 xenograft mice. (A) Representative images of H&E-stained sections of major organs (heart, liver, spleen, lung, and kidney). (B) Average body weight. Data are presented as mean ± s.d. (n = 7 mice).

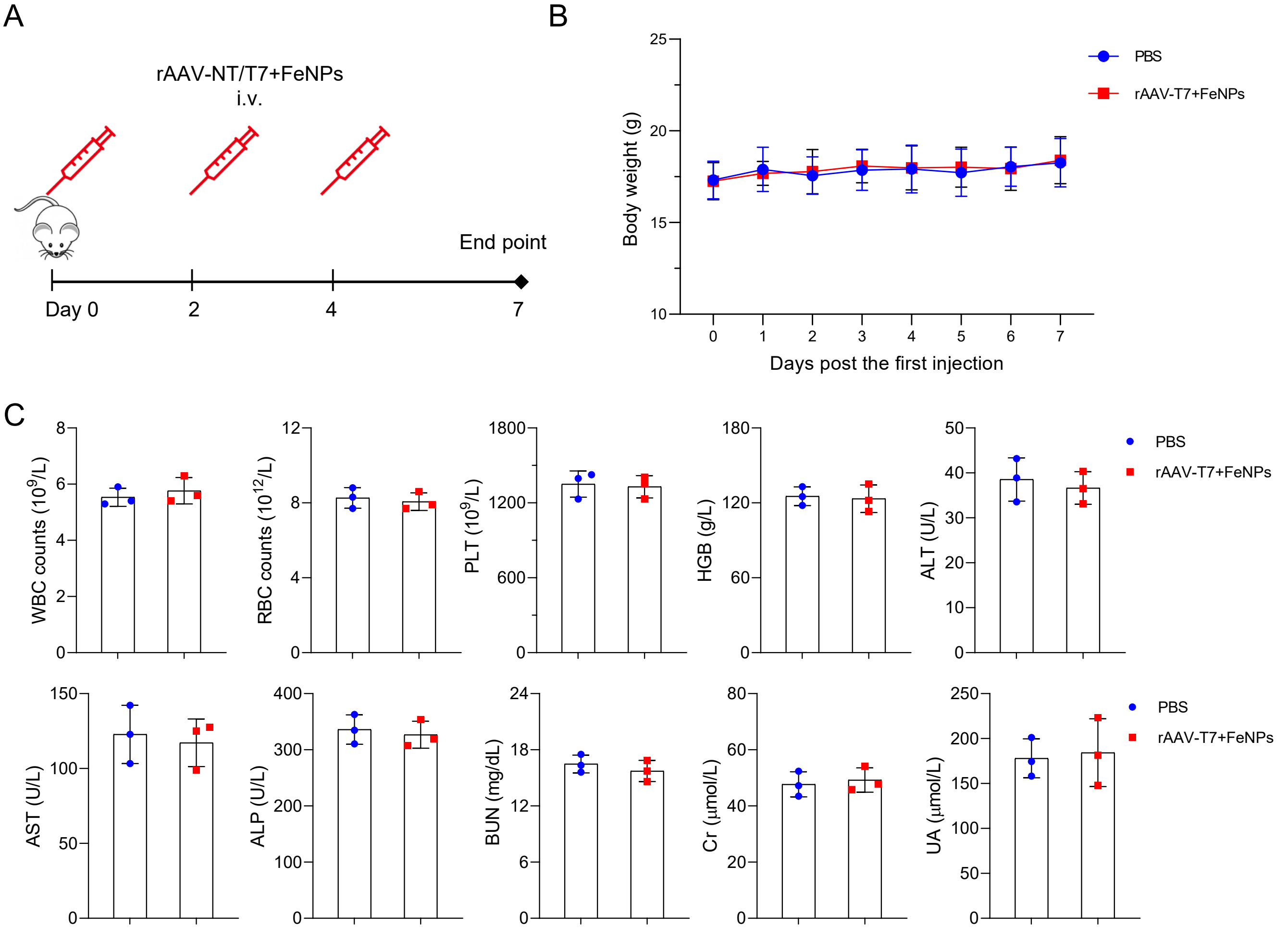


**Figure S23.** Safety evaluations of the FAST. (A) Schematic illustration for the FAST treatment.(B) Average body weight of all mice in PBS and FAST treatment groups (n = 5 mice). (C) Routine blood test and serum biochemical indices detection (n = 3 mice). WBC, white blood cell; RBC, red blood cell; PLT, platelet, HGB, hemoglobin; ALT, Alanine aminotransferase; AST, Aspartate aminotransferase; ALP, alkaline phosphatase; BUN, blood urea nitrogen; Cr, creatinine; UA, uric acid.

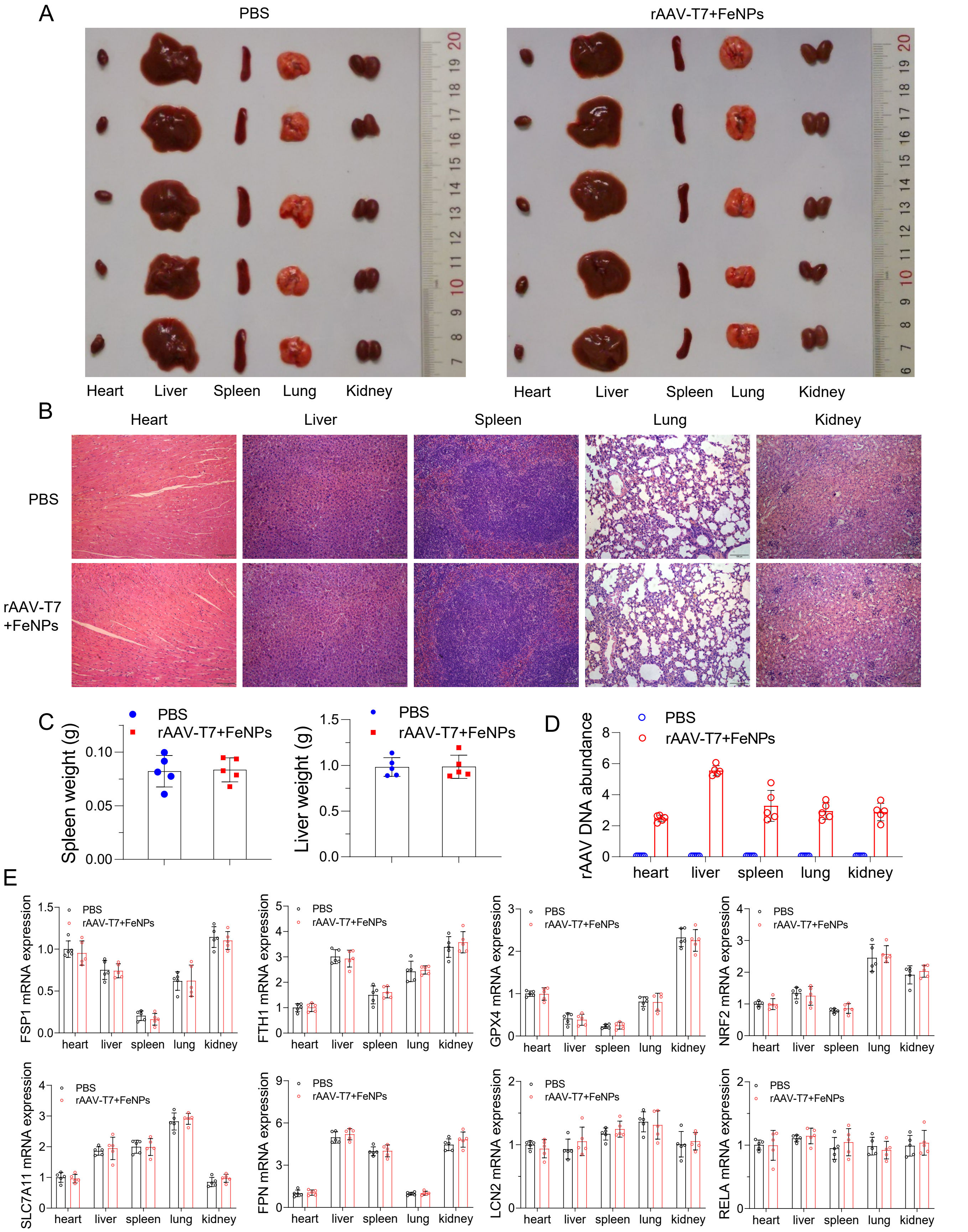


**Figure S24.** Safety evaluations of the FAST. The systematic toxicity of FAST to major organs of BALB/c mice was assessed. (A) Photos of dissected heart, liver, spleen, lung, and kidney of the PBS- and FAST-treated mice. (B) Representative images of H&E-stained tissue sections of heart, liver, spleen, lung, and kidney of the PBS- and FAST-treated mice. (C) Spleen and liver weight (n = 5 mice). (D) Abundance of virus DNA in tissues (n = 5 mice). (E) FSP1, FTH1, GPX4, NRF2, SLC7A11, FPN, LCN2, and RELA mRNA expression in tissues (n = 5 mice).

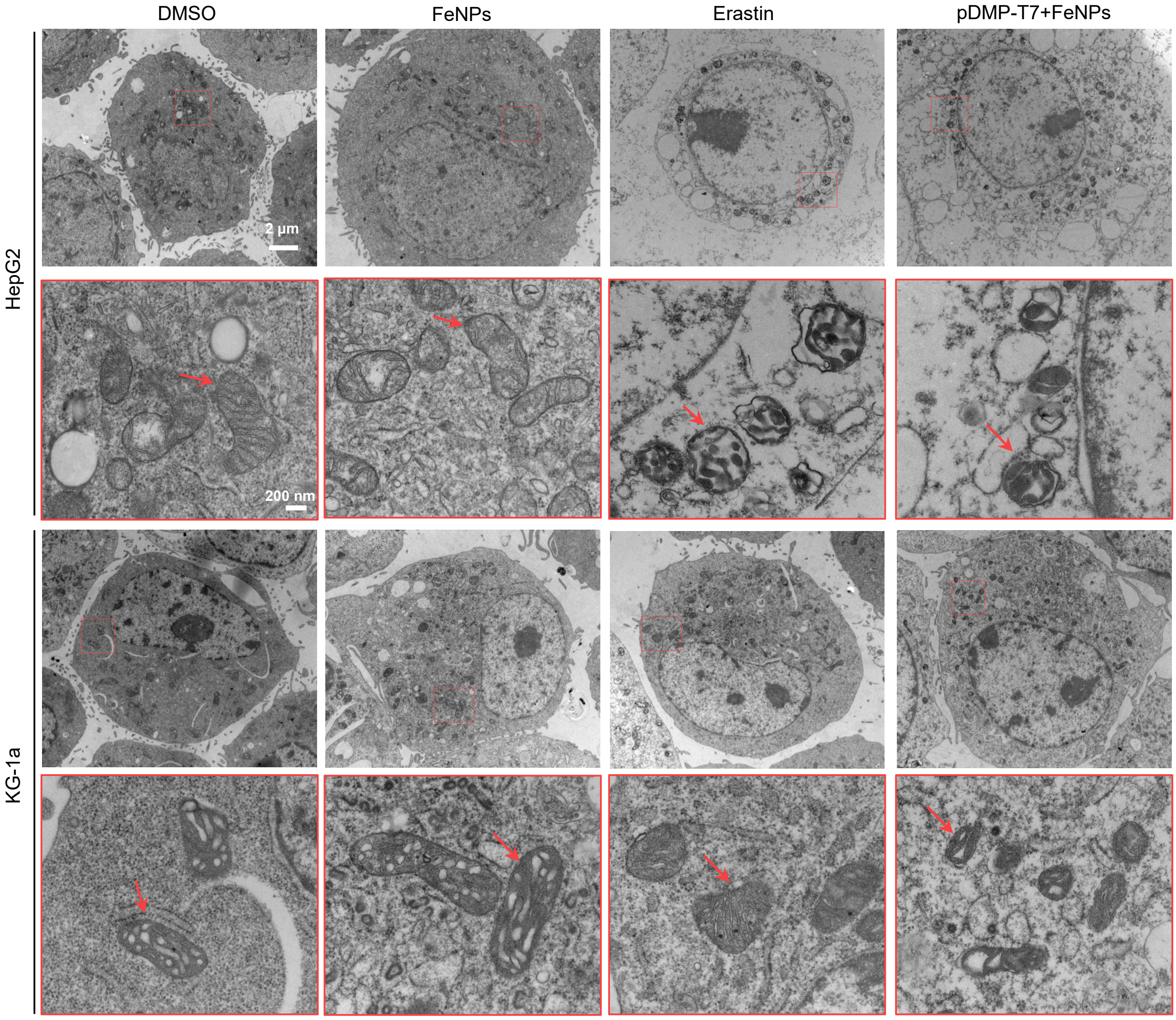


**Figure S25.** TEM images of HepG2 and KG-1a cells after the treatment of DMSO (48 h, 0.1%), FeNPs (48 h, 50 μg/mL), Erastin (8 h, 10 μM), and pDMP-T7+FeNPs (FAST; plasmid transfection overnight and then incubated with 50 μg/mL FeNPs for 48 h), respectively. The below outlined images represent the amplified areas (in red box) in up images to show mitochondria (red arrow).

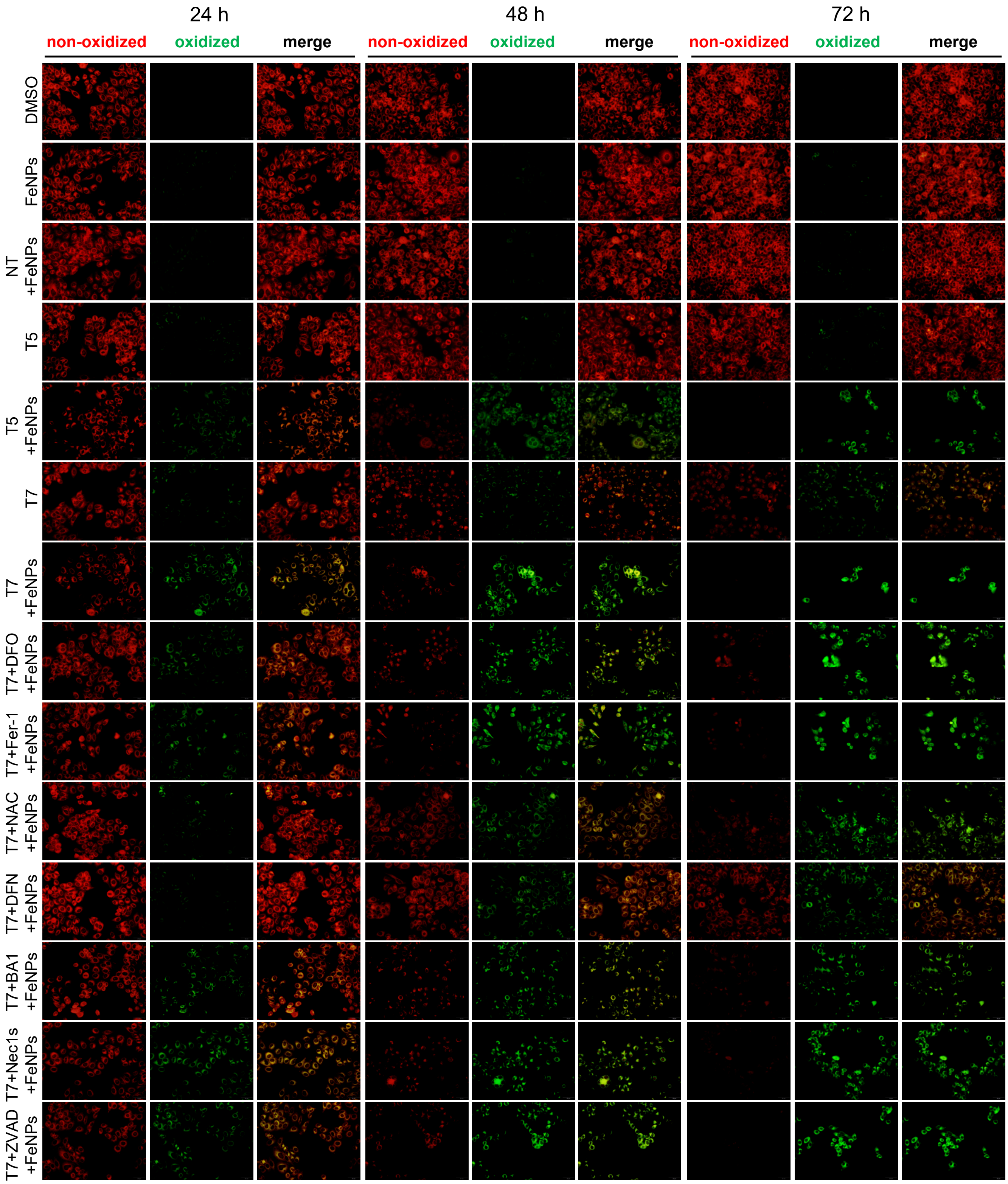


**Figure S26.** Detection of lipid ROS. Representative images of HepG2 cells stained with C11-BODIPY. Cells were transfected with various plasmids (pDMP-NT/T5/T7) overnight. The transfected cells were incubated with 50 μg/mL FeNPs for 24 h, 48 h, 72 h. For groups treated with inhibitors, the transfected cells were co-incubated with 50 μg/mL FeNPs and indicated inhibitors for 24 h, 48 h, 72 h. Cells were then stained with C11-BODIPY and imaged by fluorescence microscope. Fer-1 (1 μM); DFO (100 μM); NAC (1 mM); DFN (a mixture of 1 μM Fer-1, 100 μM DFO and 1 mM NAC); BA1 (1 nM); Nec1s (10 μM); ZVAD (50 μM). Fer-1, ferrostatin-1; DFO, deferoxamine; NAC, N-acetylcysteine; ZVAD, ZVAD-FMK; Nec1s, Necrostatin-1s; BA1, Bafilomycin A1; NT, pDMP-NT; T5, pDMP-T5; T7, pDMP-T7.

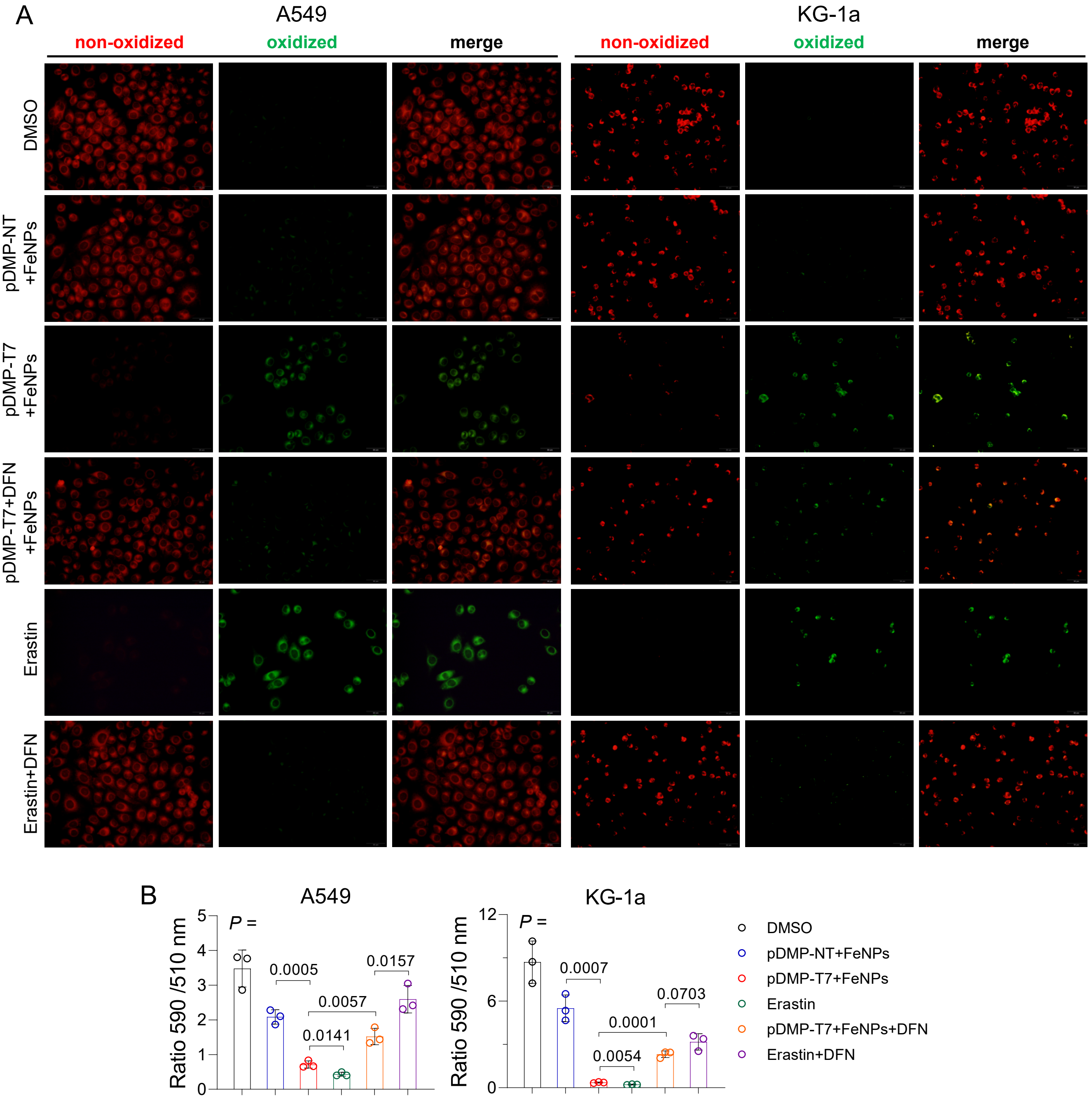


**Figure S27.** Detection of lipid ROS. (A) Representative images of A549 and KG-1a cells stained with C11-BODIPY. Cells were transfected with pDMP-NT/T7 overnight and then incubated with 50 μg/mL FeNPs for 72 h. Cells were exposed to erastin (10 μM) for 8 h as a positive control of ferroptosis. For groups treated with inhibitors, the transfected cells were co-incubated with 50 μg/mL FeNPs and DFN for 72 h, and the erastin-treated cell were co-incubated with erastin (10 μM) and DFN for 8 h. DFN, a mixture of 1 μM Fer-1, 100 μM DFO and 1 mM NAC. (B) Quantified results of lipid peroxidation. The lipid peroxidation in cells were determined by quantitating the fluorescence intensities analyzed by ImageJ software and calculating the ratio of intensity in 590 to 510 channels (n = 3 images). Fer-1, ferrostatin-1; DFO, deferoxamine; NAC, N-acetylcysteine.

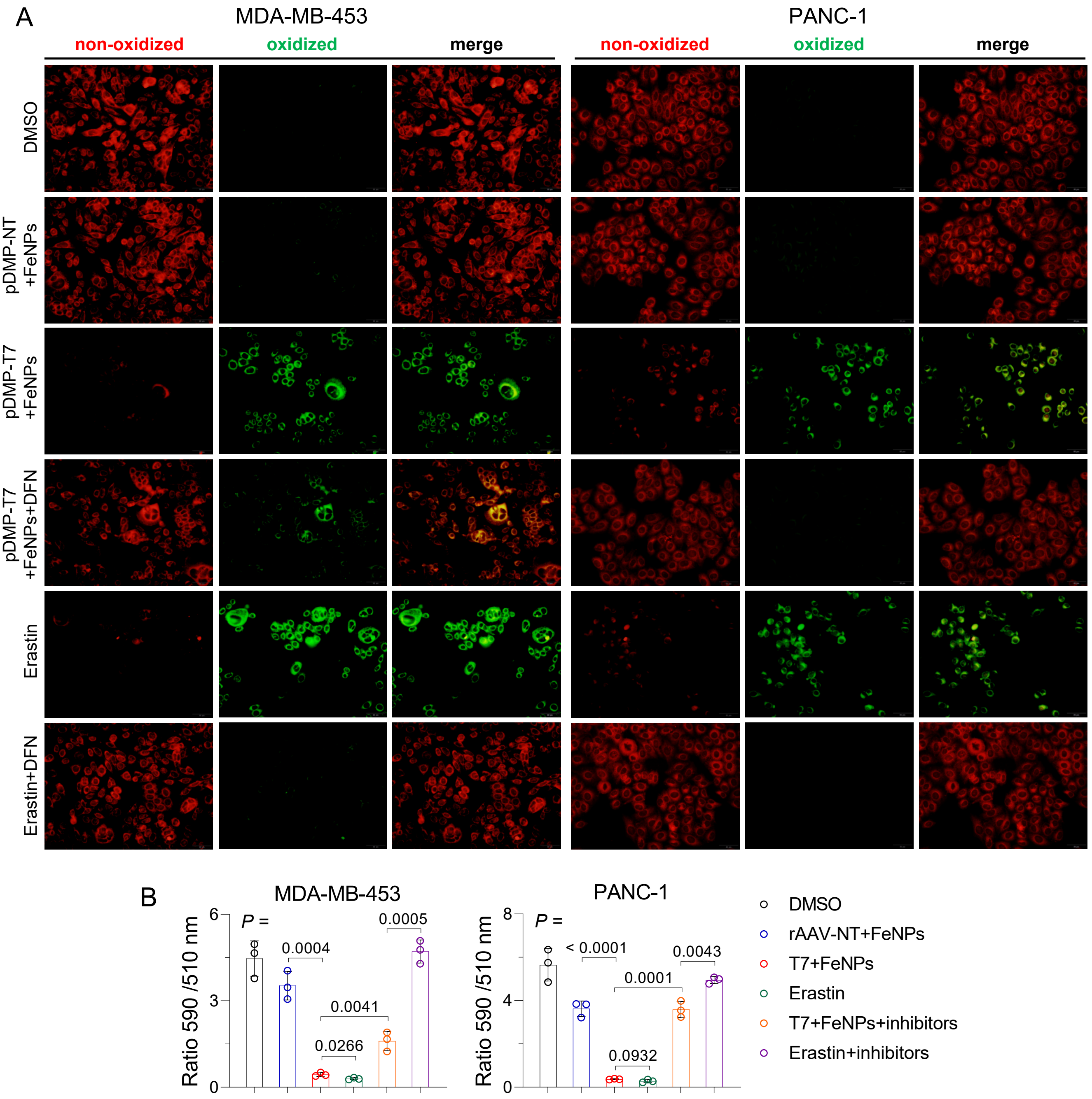


**Figure S28.** Detection of lipid ROS. (A) Representative images of MDA-MB-453 and PANC-1 cells stained with C11-BODIPY. Cells were transfected with pDMP-NT/T7 overnight and then incubated with 50 μg/mL FeNPs for 72 h. Cells were exposed to erastin (10 μM) for 8 h as a positive control of ferroptosis. For groups treated with inhibitors, the transfected cells were co-incubated with 50 μg/mL FeNPs and DFN for 72 h, and the erastin-treated cell were co-incubated with erastin (10 μM) and DFN for 8 h. DFN, a mixture of 1 μM Fer-1, 100 μM DFO and 1 mM NAC. (B) Quantified results of lipid peroxidation. The lipid peroxidation in cells were determined by quantitating the fluorescence intensities analyzed by ImageJ software and calculating the ratio of intensity in 590 to 510 channels (n = 3 images). Fer-1, ferrostatin-1; DFO, deferoxamine; NAC, N-acetylcysteine.

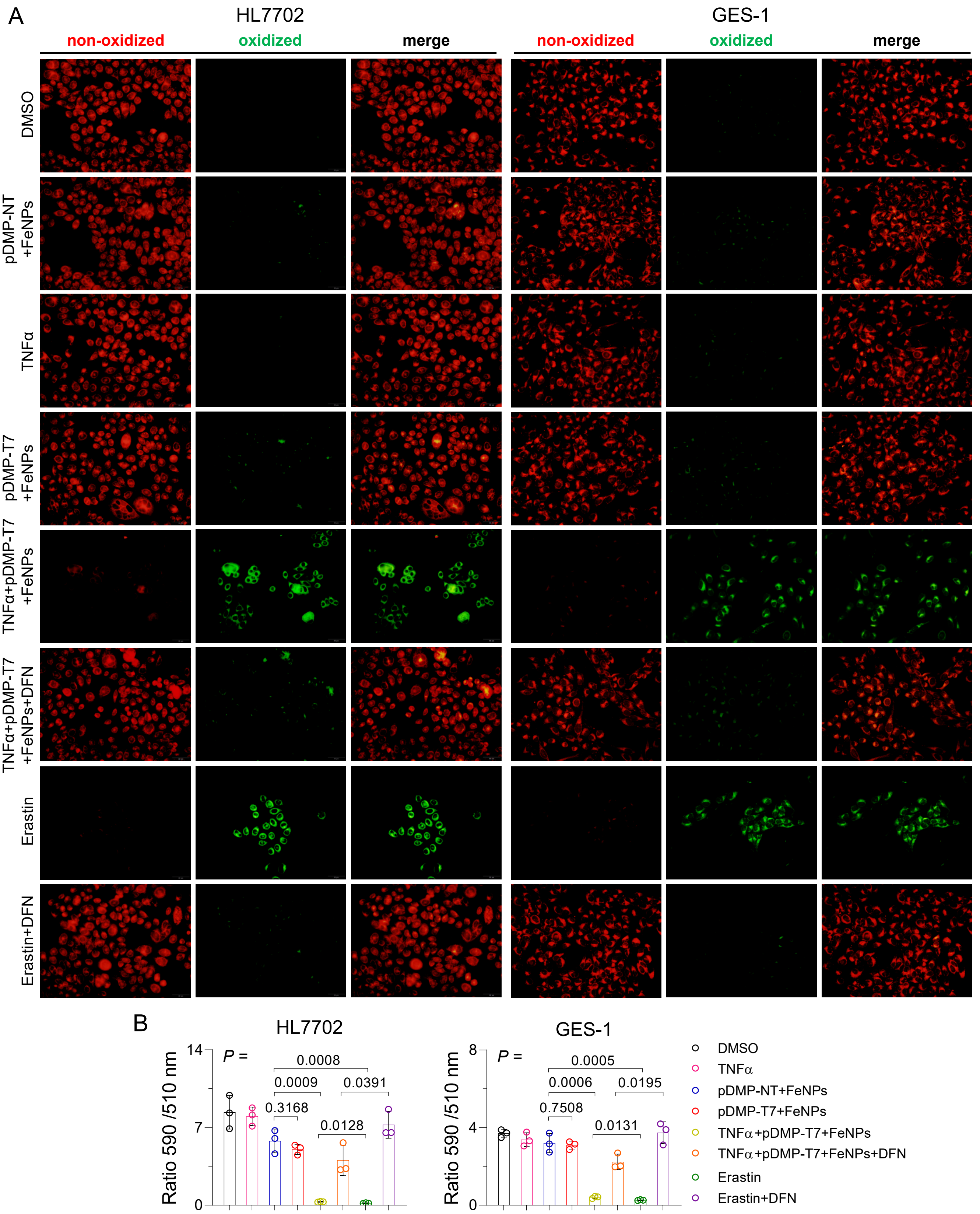


**Figure S29.** Detection of lipid ROS. Representative images of HL7702 and GES-1 cells stained with C11-BODIPY. Cells were induced with or without TNFα at a final concentration of 10 ng/mL for 1 h before transfection. Cells were transfected with pDMP-NT/T7 overnight and then co-incubated with FeNPs and DFN for 72 h. Cells were also incubated with erastin (10 μM) or co-incubated with erastin (10 μM) and DFN for 8 h as controls. DFN, a mixture of 1 μM Fer-1, 100 μM DFO and 1 mM NAC. (B) Quantified results of lipid peroxidation. The lipid peroxidation in cells were determined by quantitating the fluorescence intensities analyzed by ImageJ software and calculating the ratio of intensity in 590 to 510 channels (n = 3 images). Fer-1, ferrostatin-1; DFO, deferoxamine; NAC, N-acetylcysteine.

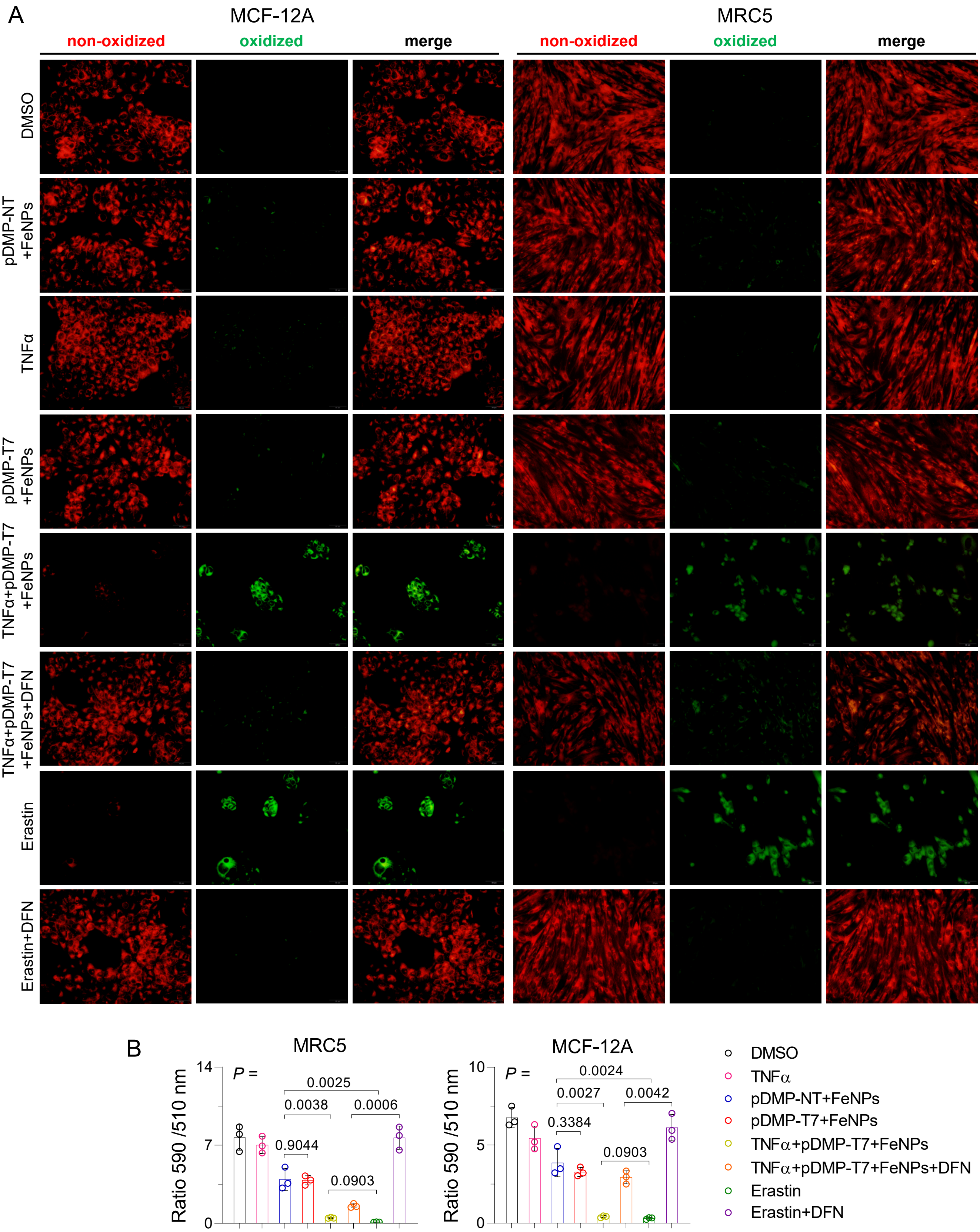


**Figure S30.** Detection of lipid ROS. (A) Representative images of MCF-12A and MRC5 cells stained with C11-BODIPY. Cells were induced with or without TNFα at a final concentration of 10 ng/mL for 1 h before transfection. Cells were transfected with pDMP-NT/T7 overnight and then co-incubated with 50 µg/mL FeNPs and DFN for 72 h. Cells were also incubated with erastin (10 µM) or co-incubated with erastin (10 µM) and DFN for 8 h as controls. DFN, a mixture of 1 µM Fer-1, 100 µM DFO and 1 mM NAC. (B) Quantified results of lipid peroxidation. The lipid peroxidation in cells were determined by quantitating the fluorescence intensities analyzed by ImageJ software and calculating the ratio of intensity in 590 to 510 channels (n = 3 images). Fer-1, ferrostatin-1; DFO, deferoxamine; NAC, N-acetylcysteine.

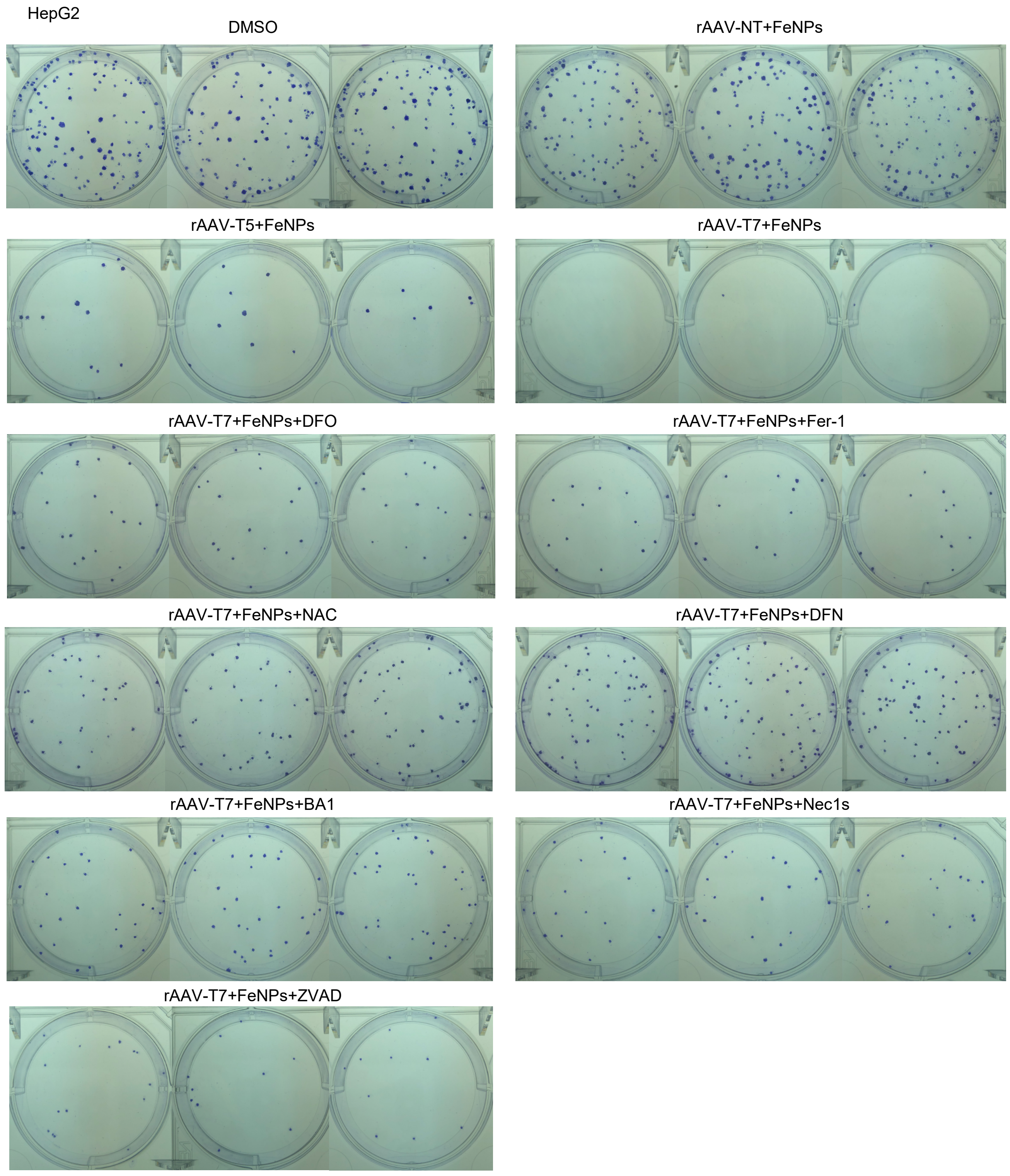


**Figure S31.** Clone formation assays of HepG2. Cells were infected with rAAV-NT/T5/T7 at the dose of $1\times10^5$ vg per cell for 24 h and then incubated with 50 μg/mL FeNPs for another 48 h. For groups treated with inhibitors, the transfected cells were co-incubated with 50 μg/mL FeNPs and indicated inhibitors for 48 h. Two hundred of treated cells were seeded into 6-well plate and cultured for 2 weeks. At this time, colonies were clearly visible (> 50 cells). Cells were stained with crystal violet at the final concentration of 0.02% (w/v) for 5 min at room temperature. The stained cells were imaged. Each treatment was conducted in triplicates. Fer-1 (1 μM); DFO (100 μM); NAC (1 mM); DFN (a mixture of 1 μM Fer-1, 100 μM DFO and 1 mM NAC); BA1 (1 nM); Nec1s (10 μM); ZVAD (50 μM). Fer-1, ferrostatin-1; DFO, deferoxamine; NAC, N-acetylcysteine; ZVAD, ZVAD-FMK; Nec1s, Necrostatin-1s; BA1, Bafilomycin A1.

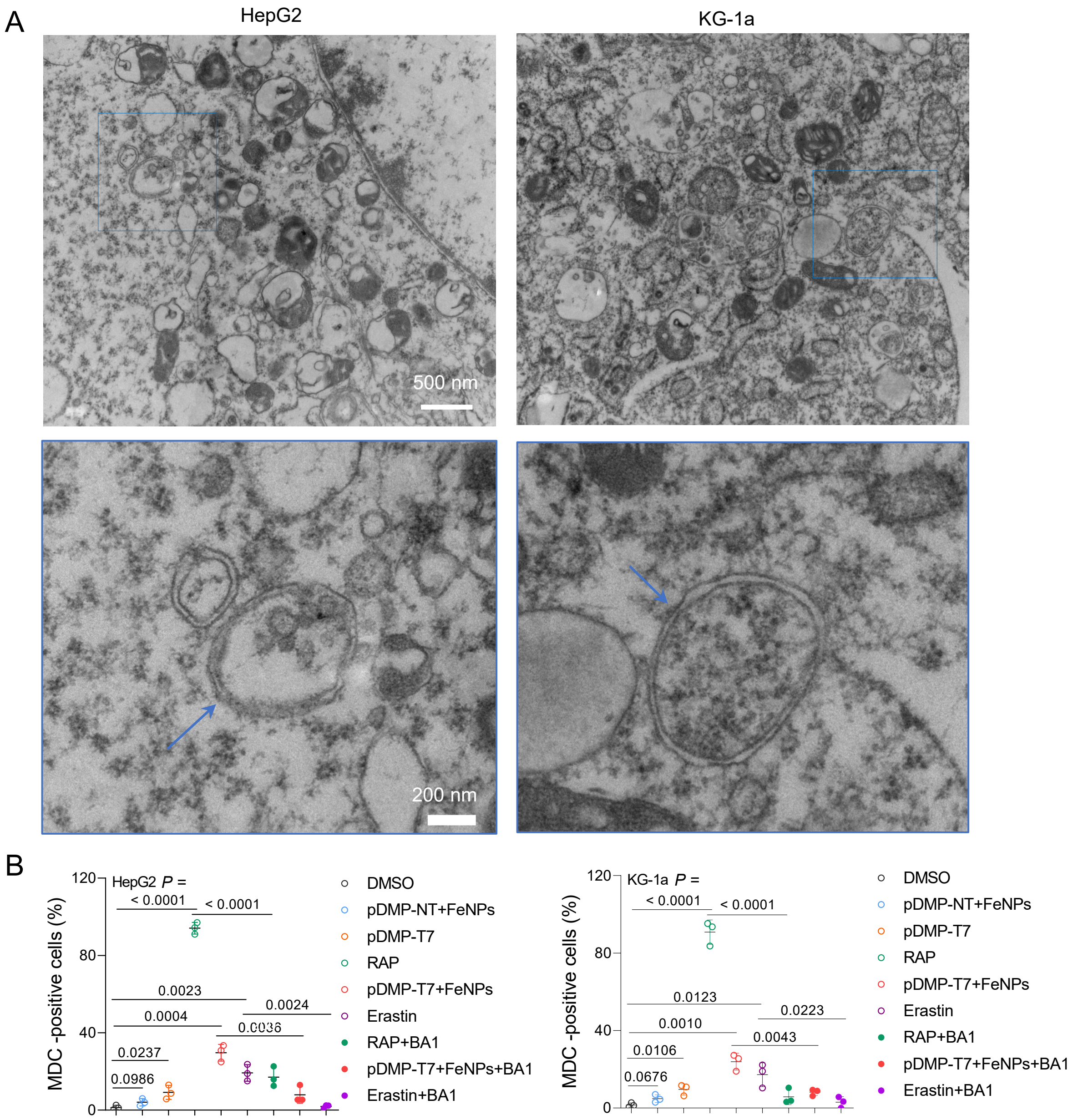


**Figure S32.** (A) Visualization of morphology of HepG2 and KG-1a cells post FAST treatment (48 h) using TEM. The below outlined images represent the amplified areas (in blue box) in up images to show autophagosome (blue arrow). (B) Quantification of Dansylcadaverine (MDC) positive cells by fluorescence microscopy. Three random fields representing 100 cells were counted. Data are shown as mean ± s.d (n = 3 biological replicates).

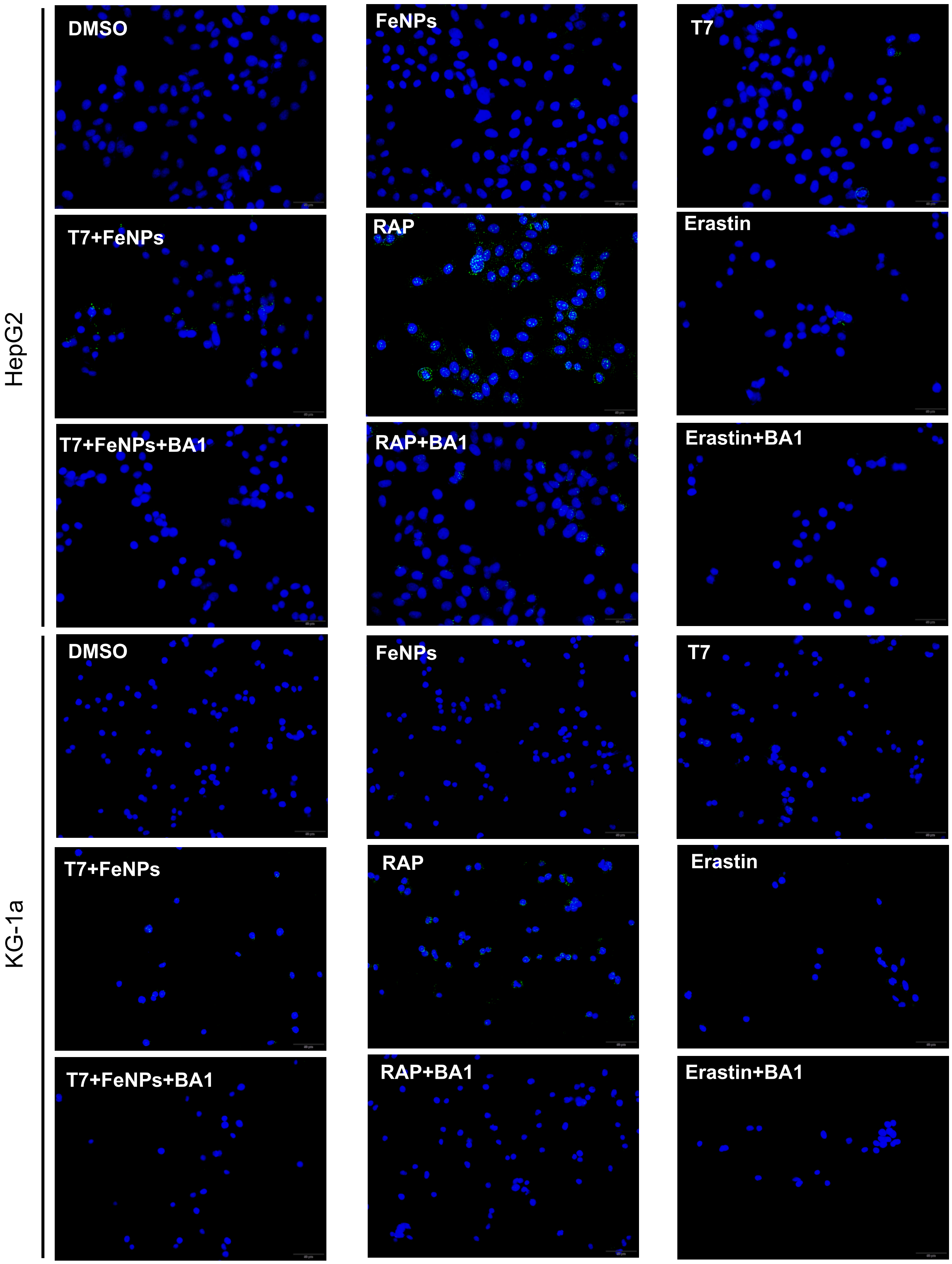


**Figure S33.** Analysis of autophagy in HepG2 and KG-1a post FAST treatment by fluorescence microscope. Cells were transfected with pDMP-T7 (T7) overnight and then incubated with or without FeNPs for 72 h. Cells treated with Rapamycin (RAP) at a final concentration of 500 nM for 12 h was used as a positive control of induced cell autophagy. Cells co-treated with Bafilomycin A1 (BA1) at a final concentration of 1 nM was used to reverse autophagy because BA1 is a typical autophagy inhibitor, in which cells were co-incubated with BA1 and FeNPs for 72 h, BA1 and RAP for 12 h, and BA1 and Erastin for 8 h, respectively. After treatment, cells were stained with DAPI and MDC dye. Blue, nucleus; green dots, autophagosome accumulation.

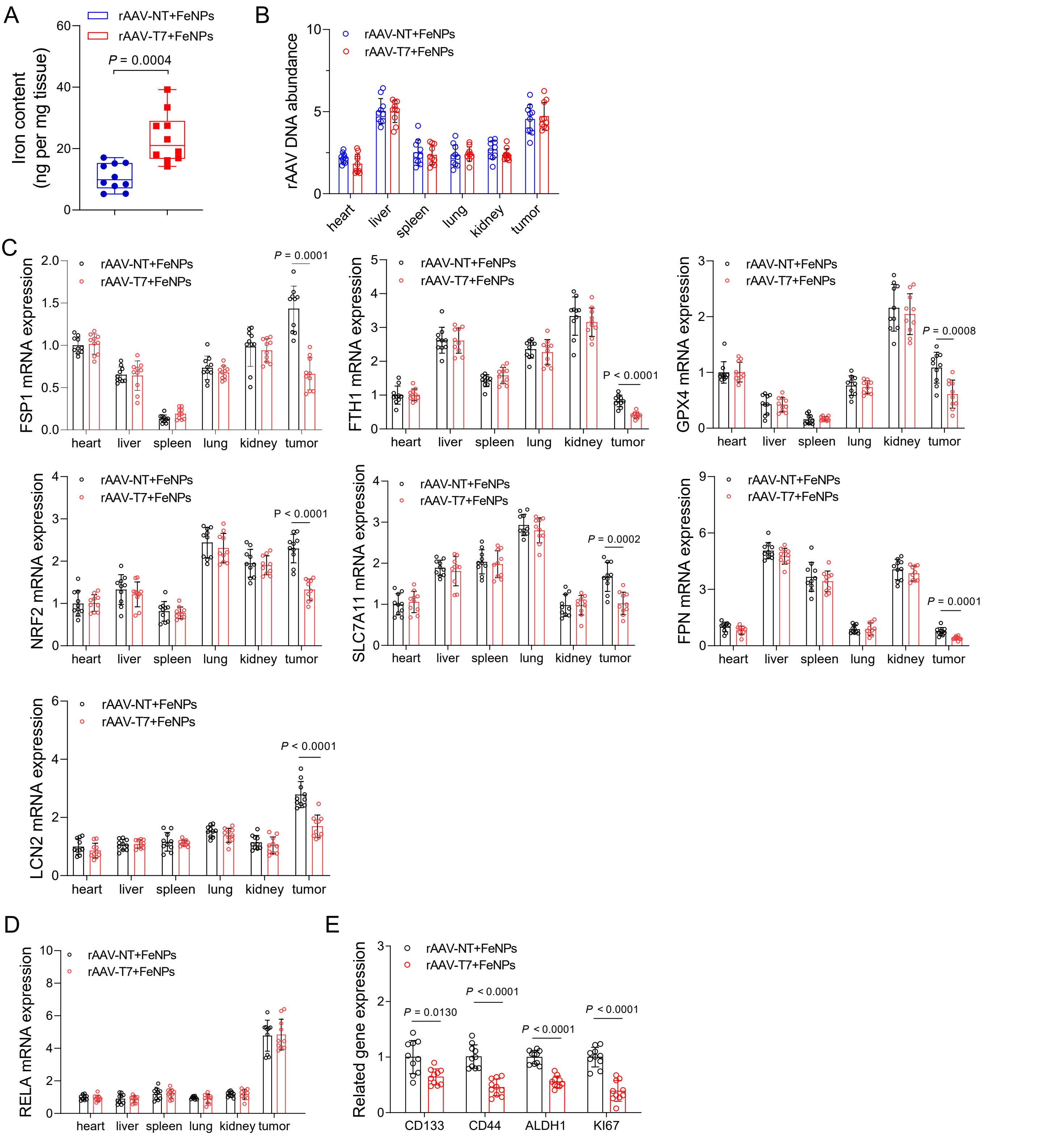


**Figure S34.** Effect of FAST on expression of targeted gene in CT-26 xenograft mice. (A) Iron content in tumor. (B) Abundance of virus DNA in tissues. (C) Expression of FSP1, FTH1, GPX4, NRF2, SLC7A11, FPN, LCN2 mRNA in tissues. (D) Expression of RELA mRNA in tissues. (E) Expression of genes as stemness- and proliferation-related markers in tumors. All data are presented as mean ± s.d. (n = 10 mice).

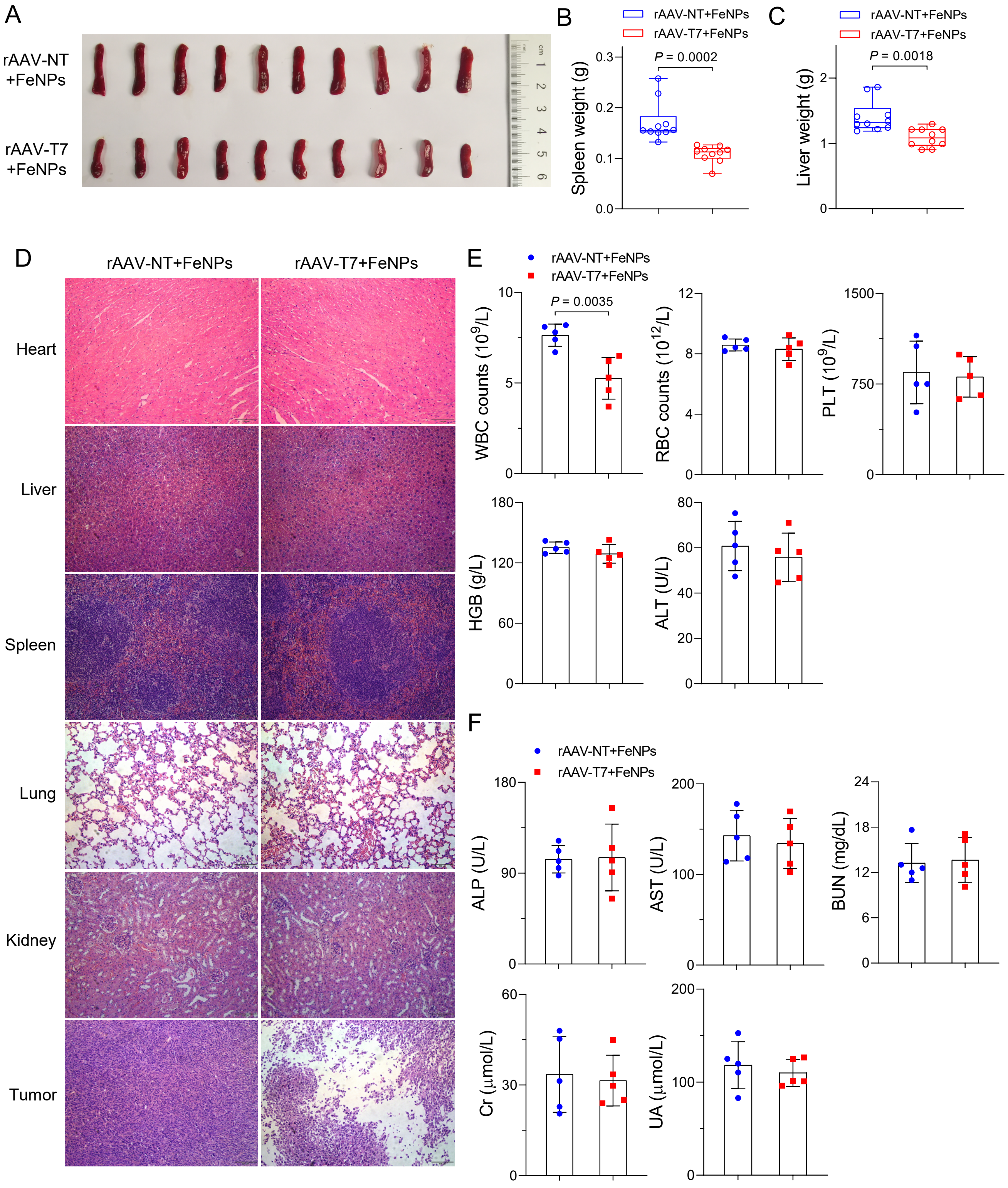


**Figure S35.** The in vivo antitumor effects of FAST in the CT-26 xenograft mice. (A) photographs of spleen. (B) Spleen weight (n = 10 mice). (C) Liver weight (n = 10 mice). (D) H&E-stained sections of major organs. (E) Routine blood test (WBC, RBC, PLT, HGB, ALT, n = 5 mice). (F) Serum biochemical indices (AST, ALP, BUN, Cr, UA, n = 5 mice). Data are presented as mean ± s.d. WBC, white blood; RBC, red blood cell; PLT, platelet, HGB, hemoglobin; ALT, Alanine aminotransferase; AST, Aspartate aminotransferase; ALP, alkaline phosphatase; BUN, blood urea nitrogen; Cr, creatinine; UA, uric acid.

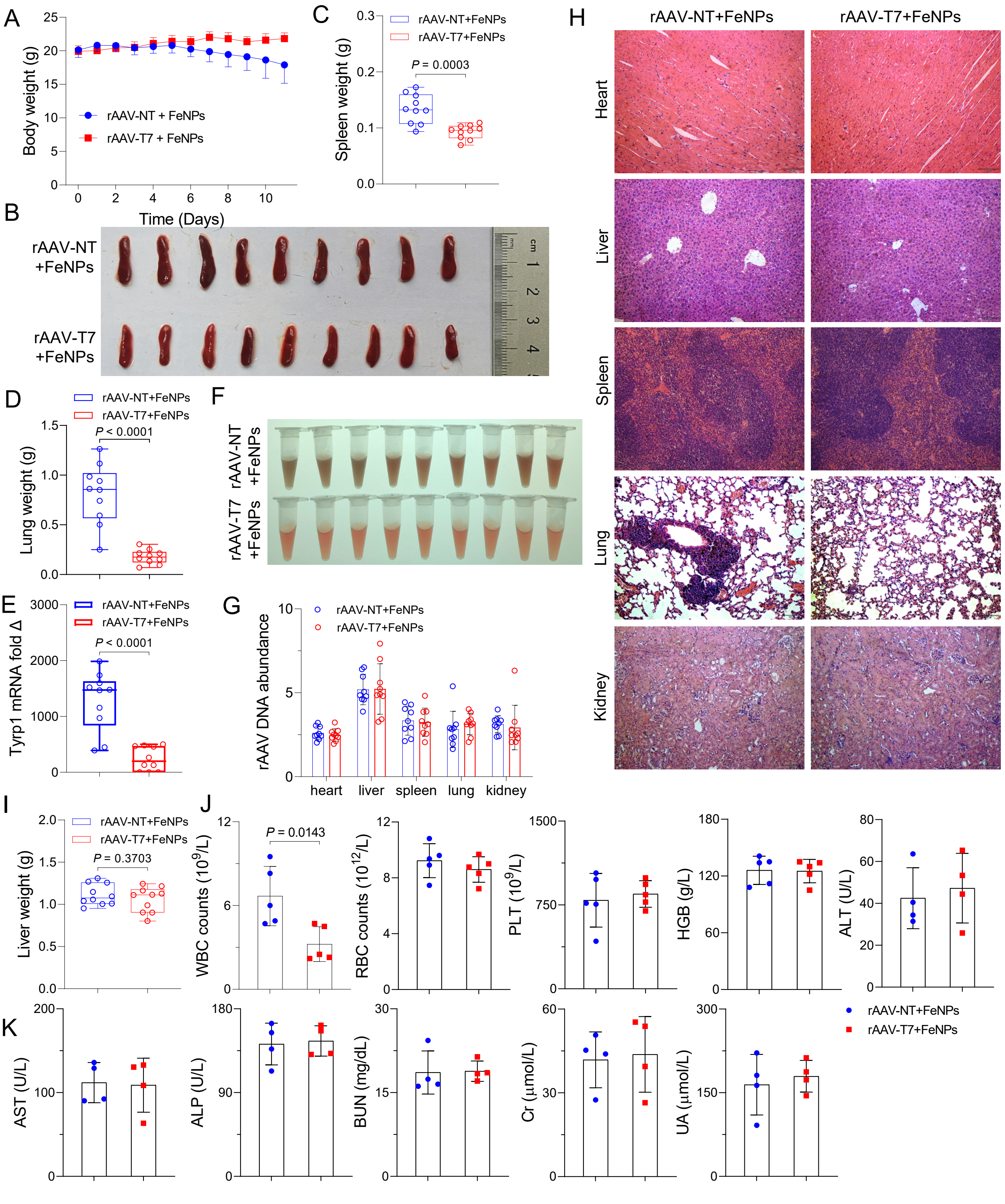


**Figure S36.** The in vivo antitumor effects of FAST in the pulmonary metastatic melanoma model. (A) Body weight (n = 9 mice). (B) Spleen photos. (C) The weight of spleen (n = 9 mice). (D) Lung weight (n = 9 mice). (E) Expression of melanocyte-specific Tyrp1 mRNA in lung detected by RT-qPCR (n = 9 mice). (F) Supernatant of ground lung extract. The solution turns black to indicate enrichment of melanin (n = 9 mice). (G) Abundance of virus DNA in tissues (n = 9 mice). (H) H&E-stained sections of major organs. (I) Liver weight (n = 9 mice). (J) Routine blood test (WBC, RBC, PLT, HGB, ALT, n = 5 mice). (K) Serum biochemical indices (AST, ALP, BUN, Cr, UA, n = 4 mice). Data are presented as mean ± s.d. WBC, white blood cell; RBC, red blood cell; PLT, platelet, HGB, hemoglobin; ALT, Alanine aminotransferase; AST, Aspartate aminotransferase; ALP, alkaline phosphatase; BUN, blood urea nitrogen; Cr, creatinine; UA, uric acid.

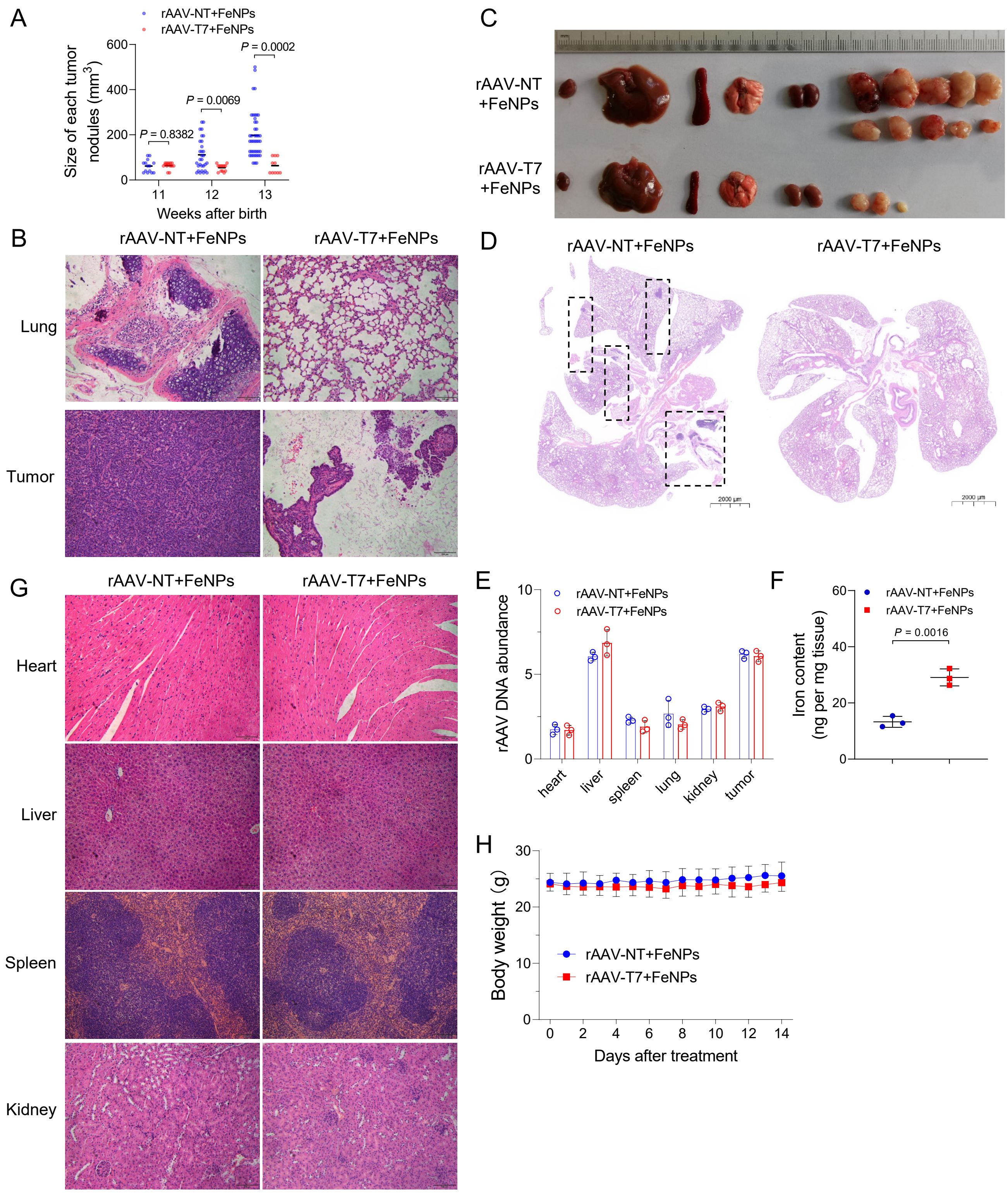


**Figure S37.** The in vivo antitumor effects of FAST in spontaneous breast cancer model. (A) Comparison of volume of each tumor nodule (n = 5 mice). (B) H&E-stained sections of lung and tumor. (C) Photos of dissected heart, liver, spleen, lung, kidney, and tumors. (D) Representative H&E-stained lung section. Dotted-line blank box indicates the metastatic foci. (E) Abundance of virus DNA in tissues (n = 3 mice). (F) Iron content in tumor (n = 3 mice). (G) H&E-stained sections of other organs. (H) Body weight (n = 5 mice).

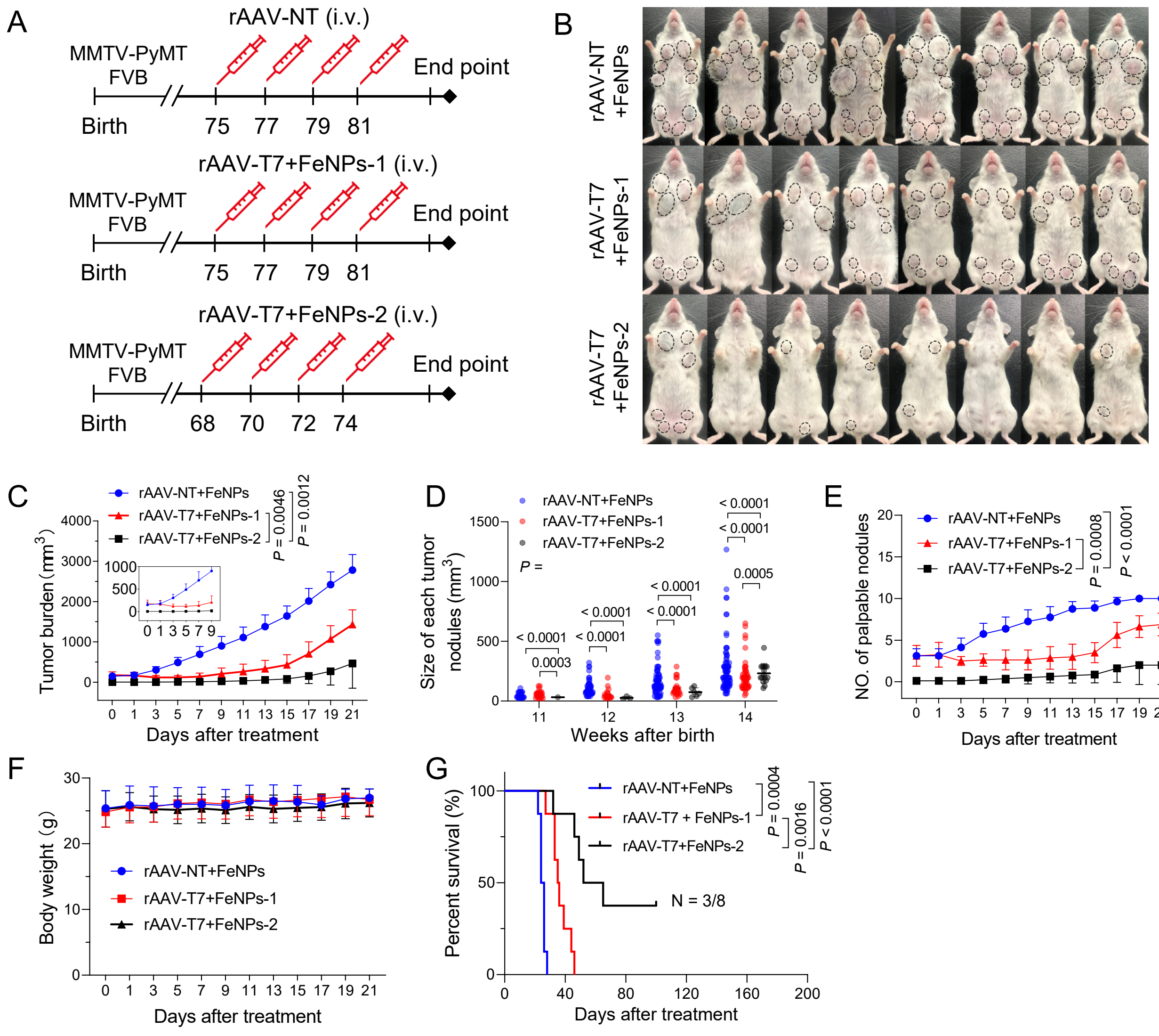


**Figure S38.** The in vivo antitumor effects of FAST in spontaneous breast cancer model. (A) Schematics of animal treatment. (B) Representative image showing gross appearance of tumors. Dotted-line circles demarcate palpable mammary tumor nodules. (C) Comparison of total tumor burden. Tumor burden was calculated as the sum volume of all tumor nodule of a mouse. (D) Comparison of volume of each tumor nodule. (E) Comparison of the number of palpable tumor nodules. (F) Body weight. (F) Kaplan-Meier survival curve. Data are presented as mean ± s.d (n = 8 mice).

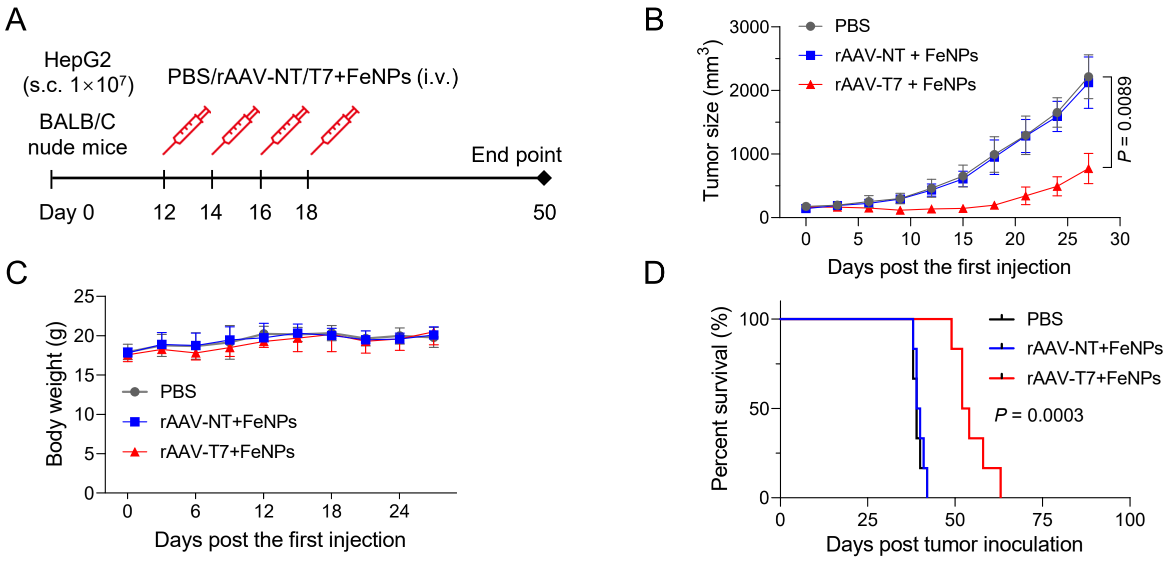


**Figure S39.** The in vivo antitumor effects of FAST in the liver cancer model of mice. The model was constructed by subcutaneously injecting the HepG2 cells. (B) Tumor growth curve. (C) Average body weight. (D) Kaplan-Meier survival curve. Data are presented as mean ± s.d. (n = 6 mice).

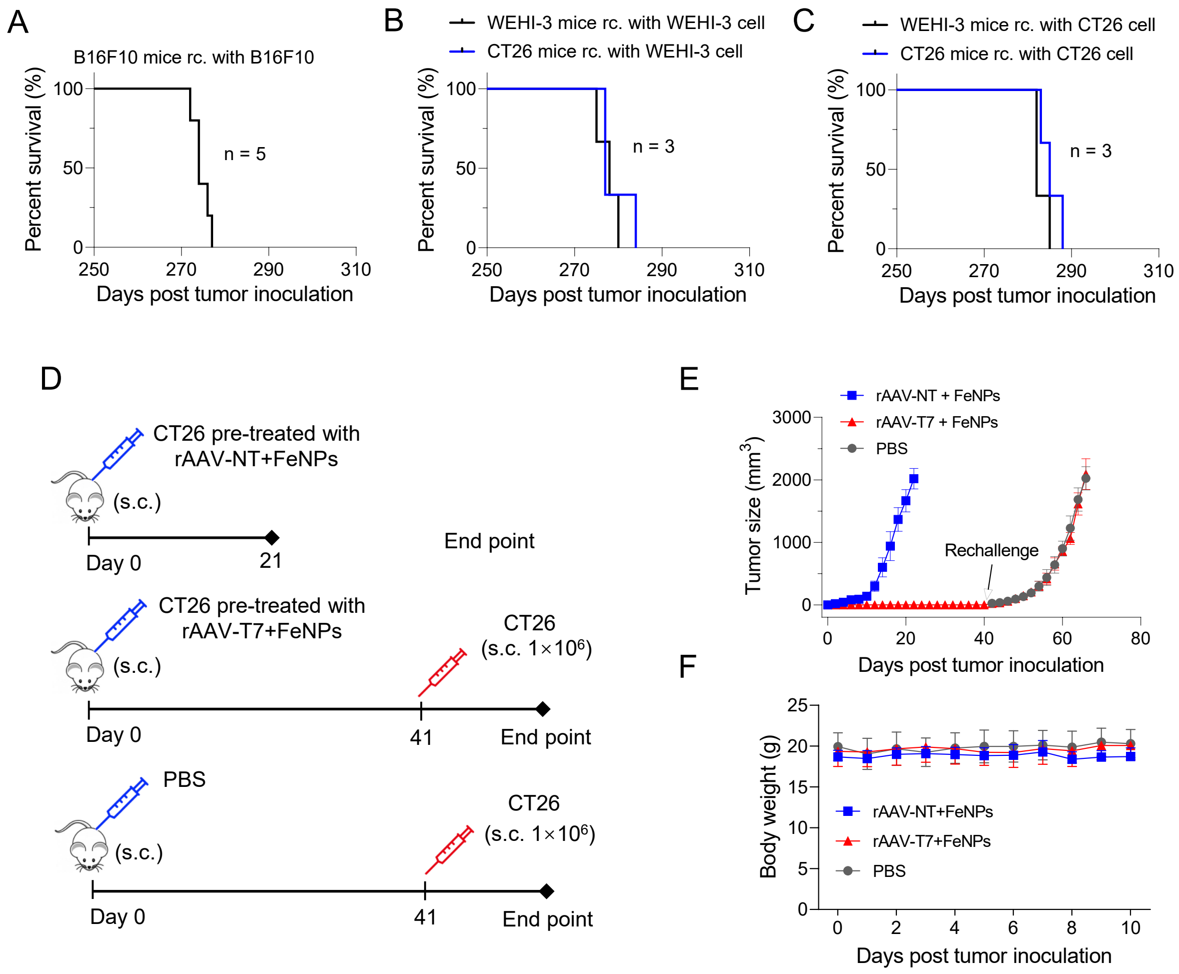


**Figure S40.** The in vivo tumorigenicity assays of FAST. (A–C) The re-challenge experiments of survived mice of three tumor models (Fig. 3H, Fig. 4F and Fig. 4L). (A) Kaplan-Meier survival curve of survived WEHI-3 xenograft mice (pulmonary metastatic melanoma mice) re-challenged with B16F10 ($n = 5$ mice). (B) Kaplan-Meier survival curve of survived WEHI-3 and CT-26 xenograft mice re-challenged with WEHI-3 ($n = 3$ mice). (C) Kaplan-Meier survival curve of survived CT-26 and WEHI-3 xenograft mice re-challenged with CT-26 ($n = 3$ mice). Data are presented as mean ± s.d. rc., re-challenged. (D–E) The re-challenge experiment with CT-26 cells. The colon cancer mice model was established with CT-26 cells. (D) Schematics of animal treatment. (E) Tumor growth curve. (F) Average body weight. Each treatment was used to three mice.

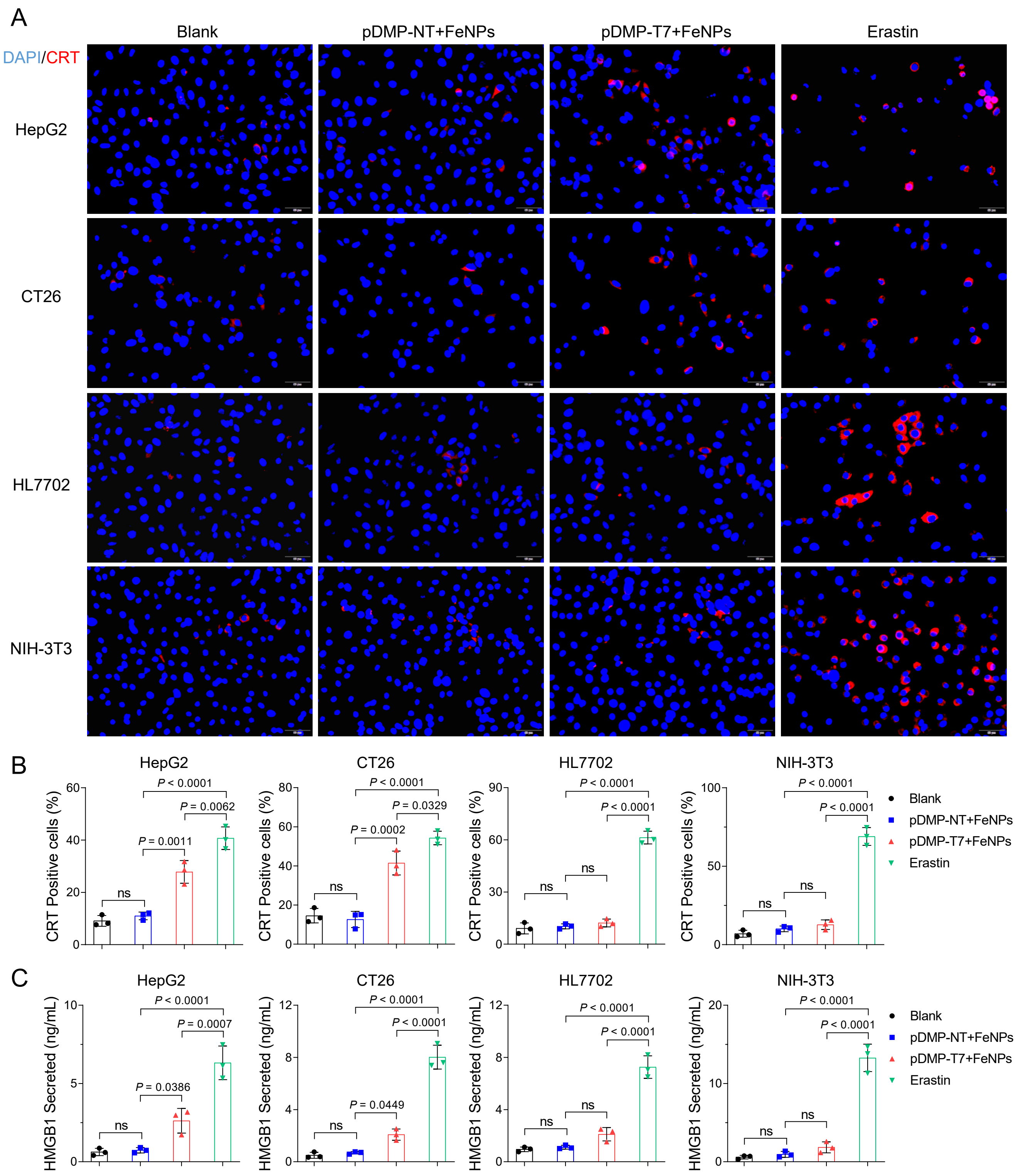


**Figure S41.** In vitro immunogenic cell death by FAST. (A) Fluorescence microscopy images of Calreticulin (CRT) expression on HepG2, CT26, HL7702, and NIH-3T3 cell lines. Cells were transfected with pDMP-NT/T7 overnight and then incubated with FeNPs for 24 h. Cells were exposed to erastin (5 μM) for 24 h as a positive control of ferroptosis. Cell nuclei was stained with DAPI (blue) and CRT was stained with Alexa-594-conjugated anti-CRT antibody (red). Scale bar: 50 μm. (B) The CRT positive cells were counted and analyzed by Image J software (n = 3 images). (C) High mobility group box-1 protein (HMGB1) released from HepG2, CT-26, HL7702, and NIH-3T3 cell lines detected by ELISA assay (n = 3 wells).

**Plasmids and the functional sequences**

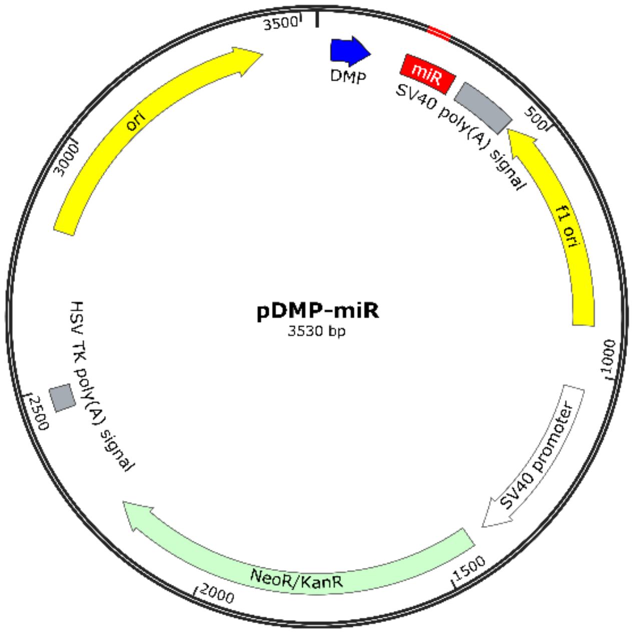


pDMP-miR

DMP+miR+SV40 poly(A) signal

GGGAATTTCCGGGGACTTTCCGGGAATTTCCGGGGACTTTCCGGGAATTTCCTAGAGGGTATATAA
TGGAAGCTCGACTTCCAGGCTAGCGAATTCGCTAAGCACTTCGTGGCCGTCGATCGTTTAAAGGGA
GGTAGTGAGTCGACCAGTGGATCCTGGAGGCTTGCTGAAGGCTGTATGCTGGAGACGCAGTGAGC
CGAGATCGCGCCACCGCGTCTCGCAGGACACAAGGCCTGTTACTAGCACTCACATGGAACAAATG
GCCCAGATCTGGCCGCACTCGAGATAACTTGTTTATTGCAGCTTATAATGGTTACAAATAAAGCAA
TAGCATCACAAATTTCACAAATAAAGCATTTTTTTCACTGCATTCTAGTTGTGGTTTGTCCAAACTC
ATCAATGTATCTTA

The sequence in the box in above skeleton vector can be replaced by the following sequences for constructing pDMP-miRNAs targeting genes of interest:

NT:

TGCTGAAATGTACTGCGCGTGGAGACGTTTTGGCCACTGACTGACGTCTCCACGCAGTACATTT

Human FSP1-1:

TGCTGCAAACAAACAAATAAAGTGGAGTTTTGGCCACTGACTGACTCCACTTTTTGTTTGTTTG

Human FSP1-2:

TGCTGTAAACAAACAAACAAATAAAGGTTTTGGCCACTGACTGACCTTTATTTTTGTTTGTTTA

Mouse FSP1-1:

TGCTGTTGGCATGCAGGCCAGCGTGGGTTTTGGCCACTGACTGACCCACGCTGCTGCATGCCAA

Mouse FSP1-2:

TGCTGAACATTGGCATGCAGGCCAGCGTTTTGGCCACTGACTGACGCTGGCCTATGCCAATGTT

Human FTH1-1:

TGCTGATCCCAAGACCTCAAAGACAAGTTTTGGCCACTGACTGACTTGTCTTTGGTCTTGGGAT

Human FTH1-2:

TGCTGTAAGGAATCTGGAAGATAGCCGTTTTGGCCACTGACTGACGGCTATCTCAGATTCCTTA

Mouse FTH1-1:
TGCTGATATTCTGCCATGCCAGCTTCGTTTTGGCCACTGACTGACGAAGCTGGTGGCAGAATAT
Mouse FTH1-2:
TGCTGTTGTCAAAGAGATATTCTGCCGTTTTGGCCACTGACTGACGGCAGAATCTCTTTGACAA
Human GPX4-1:
TGCTGTTCAGTAGGCGGCAAAGGCGGGTTTTGGCCACTGACTGACCCGCCTTTCGCCTACTGAA
Human GPX4-2:
TGCTGAGGAACTGTGGAGAGACGGTGGTTTTGGCCACTGACTGACCACCGTCTCCACAGTTCCT
Mouse GPX4-1:
TGCTGAAAGGTTCAGGAATGGGCTCCGTTTTGGCCACTGACTGACGGAGCCCACCTGAACCTTT
Mouse GPX4-2:
TGCTGTTTCCTAGGACTTTGGCGTCCGTTTTGGCCACTGACTGACGGACGCCAGTCCTAGGAAA
Human NRF2-1:
TGCTGAGTAGTTGGCAGATCCACTGGGTTTTGGCCACTGACTGACCCAGTGGATGCCAACTACT
Human NRF2-2:
TGCTGTAAAGTAGCAGGTGAGGGCATGTTTTGGCCACTGACTGACATGCCCTCCTGCTACTTTA
Mouse NRF2-1:
TGCTGAATGTGGGCAACCTGGGAGTAGTTTTGGCCACTGACTGACTACTCCCATTGCCCACATT
Mouse NRF2-2:
TGCTGAATAGCTCCTGCCAAACTTGCGTTTTGGCCACTGACTGACGCAAGTTTCAGGAGCTATT
Human SLC7A11-1:
TGCTGATAACCTGGAGACAGCAAACAGTTTTGGCCACTGACTGACTGTTTGCTCTCCAGGTTAT
Human SLC7A11-2:
TGCTGAAATCAGCCCAGCAACTGCCAGTTTTGGCCACTGACTGACTGGCAGTTTGGGCTGATTT
Mouse SLC7A11-1:
TGCTGATTACGAGCAGTTCCACCCAGGTTTTGGCCACTGACTGACCTGGGTGGCTGCTCGTAAT
Mouse SLC7A11-2:
TGCTGTTTAGAAGACTATAGAGGTCTGTTTTGGCCACTGACTGACAGACCTCTAGTCTTCTAAA
Human FPN:
TGCTGTCTACCTGCAGCTTACATGATGTTTTGGCCACTGACTGACATCATGTACTGCAGGTAGA
Mouse FPN:
TGCTGTATACAGACTCACTGATTTGCGTTTTGGCCACTGACTGACGCAAATCAGAGTCTGTATA
Human LCN2:
TGCTGTAATGTTGCCCAGCGTGAACTGTTTTGGCCACTGACTGACAGTTCACGGGGCAACATTA
Mouse LCN2:
TGCTGTCAAGTTCTGAGTTGAGTCCTGTTTTGGCCACTGACTGACAGGACTCATCAGAACTTGA

**The map of miRNA co-expression plasmids:**

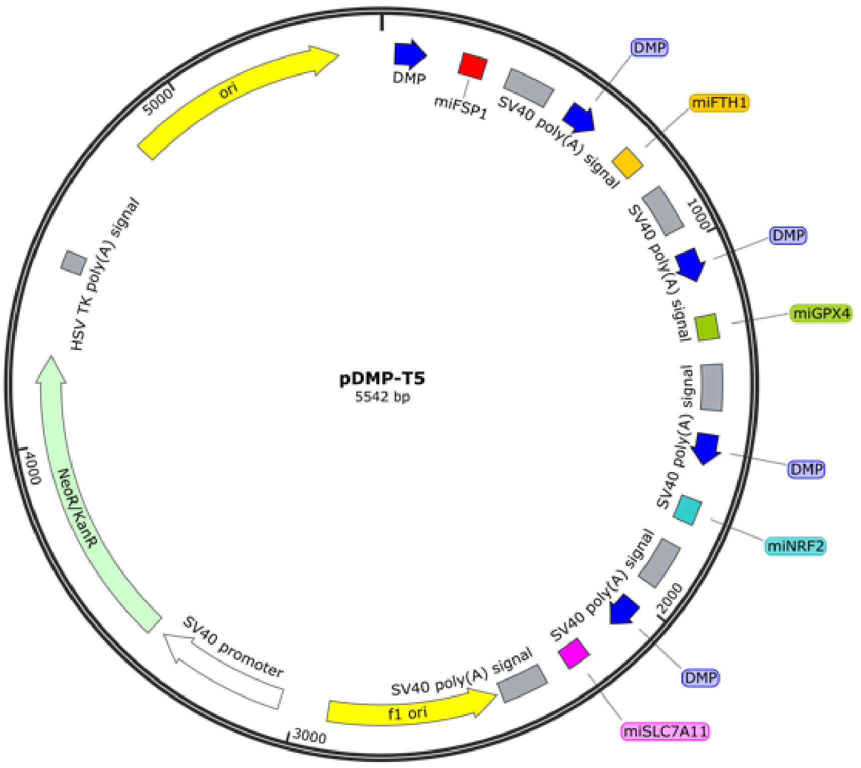


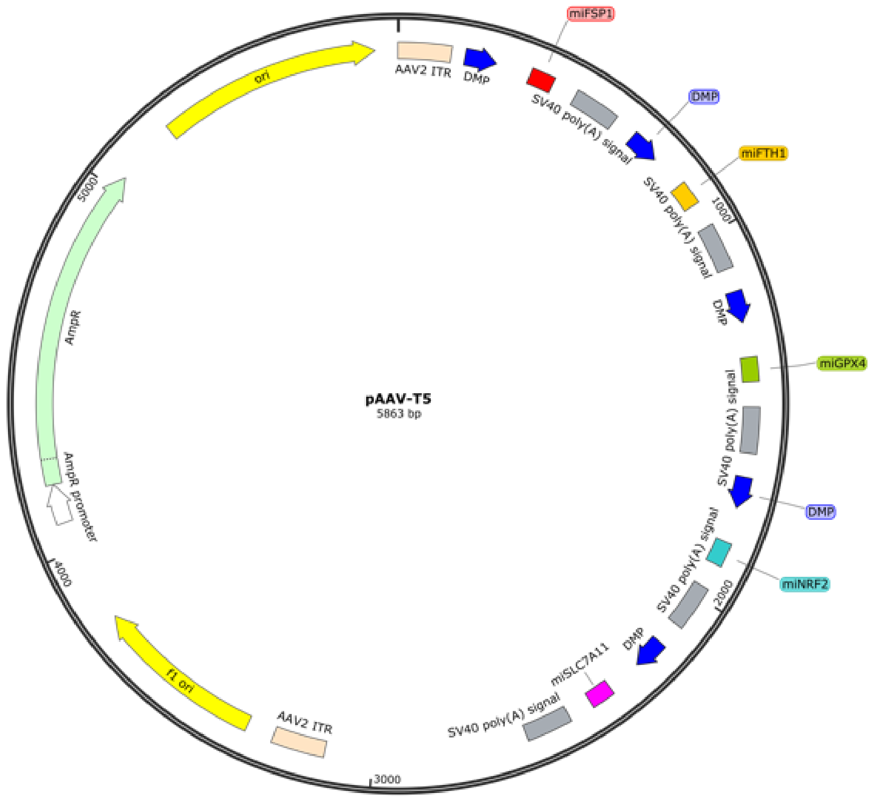

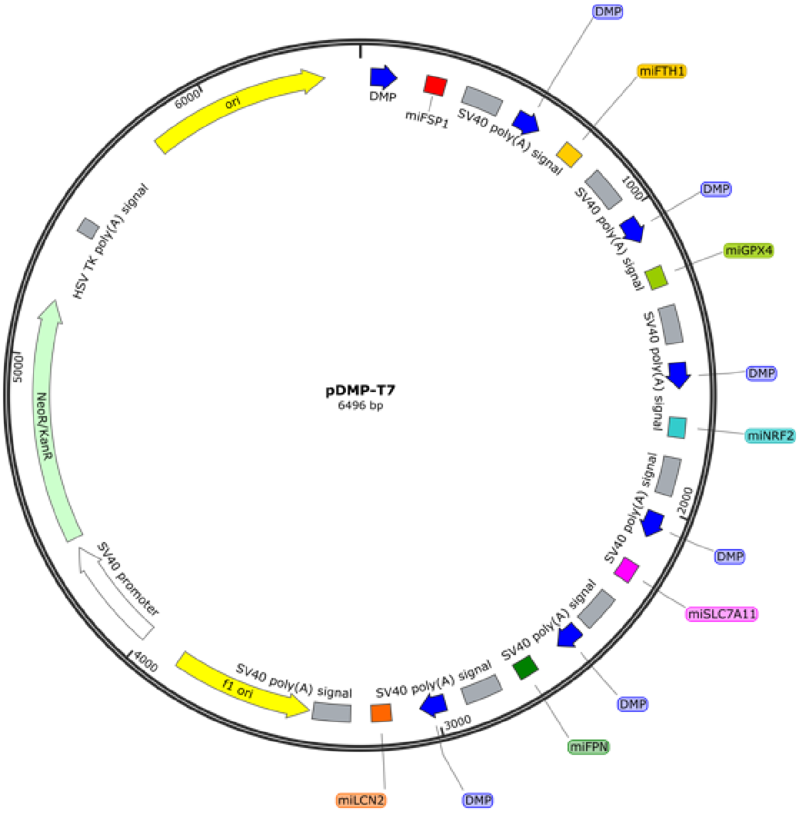
pDMP-T7
6496 bp
DMP
miFSP1
SV40 poly(A) signal
miFTH1
miGPX4
miNRF2
miSLC7A11
miFPN
miLCN2
f1 ori
SV40 promoter
NeoR/KanR
HSV TK poly(A) signal
ori
1000
2000
3000
4000
5000
6000

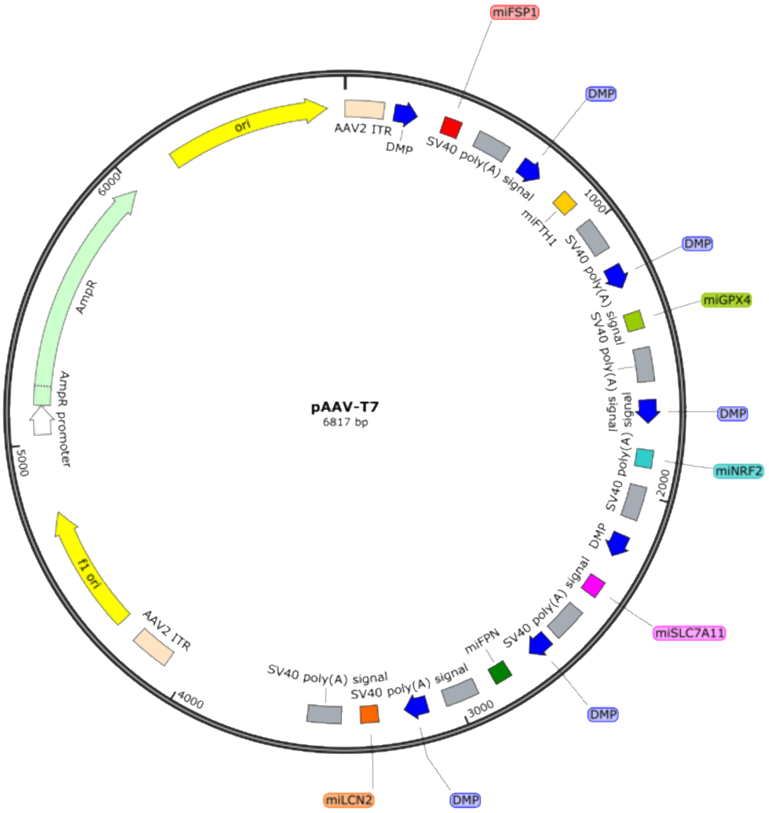
pAAV-T7
6817 bp
AAV2 ITR
DMP
miFSP1
SV40 poly(A) signal
miFTH1
miGPX4
miNRF2
miSLC7A11
miFPN
miLCN2
f1 ori
AmpR promoter
AmpR
ori
1000
2000
3000
4000
5000
6000

**DMP-T7 sequence (human):**

DMP+miFSP1+SV40 poly(A) signal-DMP+miFTH1+SV40 poly(A) signal-DMP+miGPX4+SV40 poly(A) signal DMP-miNRF2+SV40 poly(A) signal-DMP+miSLC7A11+SV40 poly(A) signal-DMP+miFPN+SV40 poly(A) signal-DMP+miLCN2+SV40 poly(A) signal

GGGAATTTCCGGGGACTTTCCGGGAATTTCCGGGGACTTTCCGGGAATTTCCTAGAGGGTATATAA
TGGAAGCTCGACTTCCAGGCTAGCGAATTCGCTAAGCACTTCGTGGCCGTCGATCGTTTAAAGGGA
GGTAGTGAGTCGACCAGTGGATCCTGGAGGCTTGCTGAAGGCTGTATGCTGCAAACAAACAAATA
AAGTGGAGTTTTGGCCACTGACTGACTCCACTTTTTGTTTGTTTGCAGGACACAAGGCCTGTTACTA
GCACTCACATGGAACAAATGGCCCAGATCTGGCCGCACTCGAGATAACTTGTTTATTGCAGCTTAT
AATGGTTACAAATAAAGCAATAGCATCACAAATTTCACAAATAAAGCATTTTTTTCACTGCATTCT
AGTTGTGGTTTGTCCAAACTCATCAATGTATCTTAAGGCGTAAATTGTAAGCGTTGCTTCGCGATG
TACGGGCATTAATGGCCTAACTGGCCGGTACCGGGAATTTCCGGGGACTTTCCGGGAATTTCCGGG
GACTTTCCGGGAATTTCCTAGAGGGTATATAATGGAAGCTCGACTTCCAGGCTAGCGAATTCGCTA
AGCACTTCGTGGCCGTCGATCGTTTAAAGGGAGGTAGTGAGTCGACCAGTGGATCCTGGAGGCTT
GCTGAAGGCTGTATGCTGTAAGGAATCTGGAAGATAGCCGTTTTGGCCACTGACTGACGGCTATCT
CAGATTCCTTACAGGACACAAGGCCTGTTACTAGCACTCACATGGAACAAATGGCCCAGATCTGG
CCGCACTCGAGATAACTTGTTTATTGCAGCTTATAATGGTTACAAATAAAGCAATAGCATCACAAA
TTTCACAAATAAAGCATTTTTTTCACTGCATTCTAGTTGTGGTTTGTCCAAACTCATCAATGTATCT
TAAGGCGTAAATTGTAAGCGTTGACGGATCGGGAGATCTCATTAATGGCCTAACTGGCCGGTACC
GGGAATTTCCGGGGACTTTCCGGGAATTTCCGGGGACTTTCCGGGAATTTCCTAGAGGGTATATAA
TGGAAGCTCGACTTCCAGGCTAGCGAATTCGCTAAGCACTTCGTGGCCGTCGATCGTTTAAAGGGA
GGTAGTGAGTCGACCAGTGGATCCTGGAGGCTTGCTGAAGGCTGTATGCTGTTCAGTAGGCGGCA
AAGGCGGGTTTTGGCCACTGACTGACCCGCCTTTCGCCTACTGAACAGGACACAAGGCCTGTTACT
AGCACTCACATGGAACAAATGGCCCAGATCTGGCCGCACTCGAGATAACTTGTTTATTGCAGCTTA
TAATGGTTACAAATAAAGCAATAGCATCACAAATTTCACAAATAAAGCATTTTTTTCACTGCATTC
TAGTTGTGGTTTGTCCAAACTCATCAATGTATCTTAAGGCGTAAATTGTAAGCGTTCCGATCCCCTA
TGGTGCACATTAATGGCCTAACTGGCCGGTACCGGGAATTTCCGGGGACTTTCCGGGAATTTCCGG
GGACTTTCCGGGAATTTCCTAGAGGGTATATAATGGAAGCTCGACTTCCAGGCTAGCGAATTCGCT
AAGCACTTCGTGGCCGTCGATCGTTTAAAGGGAGGTAGTGAGTCGACCAGTGGATCCTGGAGGCT
TGCTGAAGGCTGTATGCTGTAAAGTAGCAGGTGAGGGCATGTTTTGGCCACTGACTGACATGCCCT
CCTGCTACTTTACAGGACACAAGGCCTGTTACTAGCACTCACATGGAACAAATGGCCCAGATCTGG
CCGCACTCGAGATAACTTGTTTATTGCAGCTTATAATGGTTACAAATAAAGCAATAGCATCACAAA
TTTCACAAATAAAGCATTTTTTTCACTGCATTCTAGTTGTGGTTTGTCCAAACTCATCAATGTATCT
TAAGGCGTAAATTGTAAGCGTTCTGCTCTGATGCCGCATAGATTAATGGCCTAACTGGCCGGTACC
GGGAATTTCCGGGGACTTTCCGGGAATTTCCGGGGACTTTCCGGGAATTTCCTAGAGGGTATATAA
TGGAAGCTCGACTTCCAGGCTAGCGAATTCGCTAAGCACTTCGTGGCCGTCGATCGTTTAAAGGGA
GGTAGTGAGTCGACCAGTGGATCCTGGAGGCTTGCTGAAGGCTGTATGCTGATAACCTGGAGACA
GCAAACAGTTTTGGCCACTGACTGACTGTTTGCTCTCCAGGTTATCAGGACACAAGGCCTGTTACT
AGCACTCACATGGAACAAATGGCCCAGATCTGGCCGCACTCGAGATAACTTGTTTATTGCAGCTTA
TAATGGTTACAAATAAAGCAATAGCATCACAAATTTCACAAATAAAGCATTTTTTTCACTGCATTC
TAGTTGTGGTTTGTCCAAACTCATCAATGTATCTTAAAgcATTAATGGCCTAACTGGCCGGTACCGG
GAATTTCCGGGGACTTTCCGGGAATTTCCGGGGACTTTCCGGGAATTTCCTAGAGGGTATATAATG

GAAGCTCGACTTCCAGGCTAGCGAATTCGCTAAGCACTTCGTGGCCGTCGATCGTTTAAAGGGAG
GTAGTGAGTCGACCAGTGGATCCTGGAGGCTTGCTGAAGGCTGTATGCTGTCTACCTGCAGCTTAC
ATGATGTTTTGGCCACTGACTGACATCATGTACTGCAGGTAGACAGGACACAAGGCCTGTTACTAG
CACTCACATGGAACAAATGGCCCAGATCTGGCCGCACTCGAGATAACTTGTTTATTGCAGCTTATA
ATGGTTACAAATAAAGCAATAGCATCACAAATTTCACAAATAAAGCATTTTTTTCACTGCATTCTA
GTTGTGGTTTGTCCAAACTCATCAATGTATCTTAAGGCGTAAATTGTAAGCGTTCAGGGCTGGCAC
TCTGTCGATTAATGGCCTAACTGGCCGGTACCGGGAATTTCCGGGGACTTTCCGGGAATTTCCGGG
GACTTTCCGGGAATTTCCTAGAGGGTATATAATGGAAGCTCGACTTCCAGGCTAGCGAATTCGCTA
AGCACTTCGTGGCCGTCGATCGTTTAAAGGGAGGTAGTGAGTCGACCAGTGGATCCTGGAGGCTT
GCTGAAGGCTGTATGCTGTAATGTTGCCCAGCGTGAACTGTTTTGGCCACTGACTGACAGTTCACG
GGGCAACATTACAGGACACAAGGCCTGTTACTAGCACTCACATGGAACAAATGGCCCAGATCTGG
CCGCACTCGAGATAACTTGTTTATTGCAGCTTATAATGGTTACAAATAAAGCAATAGCATCACAAA
TTTCACAAATAAAGCATTTTTTTCACTGCATTCTAGTTGTGGTTTGTCCAAACTCATCAATGTATCT
TA

**DMP-T7 sequence (mouse):**

DMP+miFSP1+SV40 poly(A) signal-DMP+miFTH1+SV40 poly(A) signal-DMP+miGPX4+SV40 poly(A) signal DMP-miNRF2+SV40 poly(A) signal-DMP+miSLC7A11+SV40 poly(A) signal-DMP+miFPN+SV40 poly(A) signal-DMP+miLCN2+SV40 poly(A) signal

GGGAATTTCCGGGGACTTTCCGGGAATTTCCGGGGACTTTCCGGGAATTTCCTAGAGGGTATATAA
TGGAAGCTCGACTTCCAGGCTAGCGAATTCGCTAAGCACTTCGTGGCCGTCGATCGTTTAAAGGGA
GGTAGTGAGTCGACCAGTGGATCCTGGAGGCTTGCTGAAGGCTGTATGCTGTTGGCATGCAGGCC
AGCGTGGGTTTTGGCCACTGACTGACCCACGCTGCTGCATGCCAACAGGACACAAGGCCTGTTACT
AGCACTCACATGGAACAAATGGCCCAGATCTGGCCGCACTCGAGATAACTTGTTTATTGCAGCTTA
TAATGGTTACAAATAAAGCAATAGCATCACAAATTTCACAAATAAAGCATTTTTTTCACTGCATTC
TAGTTGTGGTTTGTCCAAACTCATCAATGTATCTTAAGGCGTAAATTGTAAGCGTTGCTTCGCGAT
GTACGGGCATTAATGGCCTAACTGGCCGGTACCGGGAATTTCCGGGGACTTTCCGGGAATTTCCGG
GGACTTTCCGGGAATTTCCTAGAGGGTATATAATGGAAGCTCGACTTCCAGGCTAGCGAATTCGCT
AAGCACTTCGTGGCCGTCGATCGTTTAAAGGGAGGTAGTGAGTCGACCAGTGGATCCTGGAGGCT
TGCTGAAGGCTGTATGCTGATATTCTGCCATGCCAGCTTCGTTTTGGCCACTGACTGACGAAGCTG
GTGGCAGAATATCAGGACACAAGGCCTGTTACTAGCACTCACATGGAACAAATGGCCCAGATCTG
GCCGCACTCGAGATAACTTGTTTATTGCAGCTTATAATGGTTACAAATAAAGCAATAGCATCACAA
ATTTCACAAATAAAGCATTTTTTTCACTGCATTCTAGTTGTGGTTTGTCCAAACTCATCAATGTATC
TTAAGGCGTAAATTGTAAGCGTTGACGGATCGGGAGATCTCATTAATGGCCTAACTGGCCGGTACC
GGGAATTTCCGGGGACTTTCCGGGAATTTCCGGGGACTTTCCGGGAATTTCCTAGAGGGTATATAA
TGGAAGCTCGACTTCCAGGCTAGCGAATTCGCTAAGCACTTCGTGGCCGTCGATCGTTTAAAGGGA
GGTAGTGAGTCGACCAGTGGATCCTGGAGGCTTGCTGAAGGCTGTATGCTGTTTCCTAGGACTTTG
GCGTCCGTTTTGGCCACTGACTGACGGACGCCAGTCCTAGGAAACAGGACACAAGGCCTGTTACT
AGCACTCACATGGAACAAATGGCCCAGATCTGGCCGCACTCGAGATAACTTGTTTATTGCAGCTTA
TAATGGTTACAAATAAAGCAATAGCATCACAAATTTCACAAATAAAGCATTTTTTTCACTGCATTC
TAGTTGTGGTTTGTCCAAACTCATCAATGTATCTTAAGGCGTAAATTGTAAGCGTTCCGATCCCCTA
TGGTGCACATTAATGGCCTAACTGGCCGGTACCGGGAATTTCCGGGGACTTTCCGGGAATTTCCGG
GGACTTTCCGGGAATTTCCTAGAGGGTATATAATGGAAGCTCGACTTCCAGGCTAGCGAATTCGCT

AAGCACTTCGTGGCCGTCGATCGTTTAAAGGGAGGTAGTGAGTCGACCAGTGGATCCTGGAGGCT
TGCTGAAGGCTGTATGCTGAATGTGGGCAACCTGGGAGTAGTTTTGGCCACTGACTGACTACTCCC
ATTGCCCACATTCAGGACACAAGGCCTGTTACTAGCACTCACATGGAACAAATGGCCCAGATCTG
GCCGCACTCGAGATAACTTGTTTATTGCAGCTTATAATGGTTACAAATAAAGCAATAGCATCACAA
ATTTCACAAATAAAGCATTTTTTTCACTGCATTCTAGTTGTGGTTTGTCCAAACTCATCAATGTATC
TTAAGGCGTAAATTGTAAGCGTTCTGCTCTGATGCCGCATAGATTAATGGCCTAACTGGCCGGTAC
CGGGAATTTCCGGGGACTTTCCGGGAATTTCCGGGGACTTTCCGGGAATTTCCTAGAGGGTATATA
ATGGAAGCTCGACTTCCAGGCTAGCGAATTCGCTAAGCACTTCGTGGCCGTCGATCGTTTAAAGGG
AGGTAGTGAGTCGACCAGTGGATCCTGGAGGCTTGCTGAAGGCTGTATGCTGATTACGAGCAGTT
CCACCCAGGTTTTGGCCACTGACTGACCTGGGTGGCTGCTCGTAATCAGGACACAAGGCCTGTTAC
TAGCACTCACATGGAACAAATGGCCCAGATCTGGCCGCACTCGAGATAACTTGTTTATTGCAGCTT
ATAATGGTTACAAATAAAGCAATAGCATCACAAATTTCACAAATAAAGCATTTTTTTCACTGCATT
CTAGTTGTGGTTTGTCCAAACTCATCAATGTATCTTAAAgcATTAATGGCCTAACTGGCCGGTACCG
GGAATTTCCGGGGACTTTCCGGGAATTTCCGGGGACTTTCCGGGAATTTCCTAGAGGGTATATAAT
GGAAGCTCGACTTCCAGGCTAGCGAATTCGCTAAGCACTTCGTGGCCGTCGATCGTTTAAAGGGA
GGTAGTGAGTCGACCAGTGGATCCTGGAGGCTTGCTGAAGGCTGTATGCTGTATACAGACTCACTG
ATTTGCGTTTTGGCCACTGACTGACGCAAATCAGAGTCTGTATACAGGACACAAGGCCTGTTACTA
GCACTCACATGGAACAAATGGCCCAGATCTGGCCGCACTCGAGATAACTTGTTTATTGCAGCTTAT
AATGGTTACAAATAAAGCAATAGCATCACAAATTTCACAAATAAAGCATTTTTTTCACTGCATTCT
AGTTGTGGTTTGTCCAAACTCATCAATGTATCTTAAGGCGTAAATTGTAAGCGTTCAGGGCTGGCA
CTCTGTCGATTAATGGCCTAACTGGCCGGTACCGGGAATTTCCGGGGACTTTCCGGGAATTTCCGG
GGACTTTCCGGGAATTTCCTAGAGGGTATATAATGGAAGCTCGACTTCCAGGCTAGCGAATTCGCT
AAGCACTTCGTGGCCGTCGATCGTTTAAAGGGAGGTAGTGAGTCGACCAGTGGATCCTGGAGGCT
TGCTGAAGGCTGTATGCTGTCAAGTTCTGAGTTGAGTCCTGTTTTGGCCACTGACTGACAGGACTC
ATCAGAACTTGACAGGACACAAGGCCTGTTACTAGCACTCACATGGAACAAATGGCCCAGATCTG
GCCGCACTCGAGATAACTTGTTTATTGCAGCTTATAATGGTTACAAATAAAGCAATAGCATCACAA
ATTTCACAAATAAAGCATTTTTTTCACTGCATTCTAGTTGTGGTTTGTCCAAACTCATCAATGTATC
TTA
